# Novel conserved RNA structural motifs regulate genome cyclization of yellow fever virus

**DOI:** 10.64898/2026.09.23.753880

**Authors:** Yi-Ge Luo, Jing-Yi Zhang, Dan Li, Yi-Tian Xie, Hai-Tao Lu, Xu-Meng Feng, Yu-Zhen Ding, Hang-Yu Zhou, Cheng-Feng Qin, Zhong-Yu Liu

## Abstract

The long-range RNA-RNA interaction between the termini of a flavivirus genome is controlled by multiple local RNA secondary and tertiary structures. However, a complete picture of this essential genome cyclization process is still elusive. Herein, by using selective 2′-hydroxyl acylation analyzed by primer extension (SHAPE), SHAPE and mutational profiling, *in vitro* RNA binding assays, and reverse genetics approaches, we demonstrated that the hairpin 2 in the capsid-coding region (cHP2) is required for efficient replication of yellow fever virus and functionally interacts with the 3′-small hairpin (sHP) during genome cyclization. Interestingly, through a combination of multidimensional chemical probing and functional characterization assays, it was discovered that the sHP forms previously unknown tertiary interactions with an immediately upstream GACG region embedded within the 3′-cyclization sequence. This crucial tertiary RNA motif accounts for a potential negative regulatory mechanism of genome cyclization and is universally conserved among the mosquito-borne-related flaviviruses, which constitute a major phylogenetic branch within the subgenus *Euflavivirus,* genus *Orthoflavivirus*. Our study highlights the intricacy of RNA-based regulation in flaviviruses.

## Introduction

In addition to its crucial role in protein synthesis, RNA performs functions of great diversity in virtually all biological entities, mostly due to its ability to fold into complicated secondary and tertiary structures (Batey et al., 1999; Ganser et al., 2019; Vicens and Kieft, 2022). Single-stranded, positive-sense RNA viruses are excellent examples of the functionality of RNA molecules, as *cis*-acting RNA elements, most of which are structured, exist essentially in all viral genomes of positive polarity (Liu et al., 2009; Nicholson and White, 2014). Indeed, multiple classical RNA structures were identified from the +ssRNA viruses, such as the type I-IV internal ribosome entry sites (IRES) (Jang et al., 1988; Pelletier and Sonenberg, 1988; Jackson et al., 2010; Jaafar and Kieft, 2019), the −1 programmed ribosomal frameshift pseudoknot (Plant et al., 2005; Jaafar and Kieft, 2019) and the Xrn1-resistant RNAs (Pijlman et al., 2008; Silva et al., 2010; MacFadden et al., 2018; Szucs et al., 2020; Vicens and Kieft, 2021).

The mosquito-borne flaviviruses (MBFVs, which throughout the text refers to the true mosquito-borne members), an ecological group that belongs to the family *Flaviviridae*, genus *Orthoflavivirus* (Postler et al., 2023), subgenus *Euflavivirus* (hereafter euflaviviruses) (International Committee on Taxonomy of Viruses, 2025; Simmonds et al., 2025), include a large number of human pathogens that are closely related phylogenetically, such as dengue virus (DENV), Zika virus (ZIKV), West Nile virus (WNV) and yellow fever virus (YFV) (Pierson and Diamond, 2020). Although a long-range RNA-RNA interaction between viral genome termini (usually referred to as genome cyclization) appears to be a conserved feature among the euflaviviruses (Liu et al., 2016; Liu and Qin, 2020), the genome cyclization of the MBFVs shares a similar pattern and has been extensively investigated (Khromykh et al., 2001; Corver et al., 2003; Alvarez et al., 2005a; Alvarez et al., 2005b; Filomatori et al., 2006; Alvarez et al., 2008; Zhang et al., 2008; Friebe and Harris, 2010; Villordo et al., 2010; Friebe et al., 2011; Friebe et al., 2012; Liu et al., 2013; Friedrich et al., 2014; de Borba et al., 2015; Liu et al., 2016; Li et al., 2023). Three pairs of complementary sequences, the 5′-3′ cyclization sequences (CSs) (Khromykh et al., 2001; Corver et al., 2003; Liu et al., 2017), the 5′-3′ upstream AUG regions (UARs) (Alvarez et al., 2005b; Alvarez et al., 2008; Zhang et al., 2008), and the 5′-3′ downstream AUG regions (DARs) (Friebe and Harris, 2010; Friebe et al., 2011; Li et al., 2020) are directly responsible for the genome cyclization event, which is essential for the initiation of synthesis of the negative-strand RNA (Filomatori et al., 2006; Lodeiro et al., 2009; Liu et al., 2016; Fajardo et al., 2020; Oviedo-Rouco et al., 2026). The localizations and primary sequences of the CSs are highly conserved (Hahn et al., 1987; Khromykh et al., 2001; Liu et al., 2013; Liu et al., 2016), whereas the UARs and DARs show lower conservation in their sequences (Liu et al., 2016). Moreover, these cyclization elements are frequently involved in the formation of local RNA structures that can regulate the genome cyclization process in turn (Alvarez et al., 2005b; Villordo et al., 2010; Liu et al., 2016). In all MBFVs investigated so far, the 5′-CS sequences are single-stranded and do not have a defined secondary structure prior to genome cyclization (Hahn et al., 1987; Khromykh et al., 2001; Liu et al., 2013; Liu et al., 2016; Li et al., 2023). The 3′-UAR and the 3′-DAR are embedded in the 3′-sHP and the bottom region of the terminal 3′-SL (Alvarez et al., 2005b; Alvarez et al., 2008; Villordo et al., 2010). Since these secondary structures in the MBFVs’ 3′ termini are extremely conserved (Hahn et al., 1987; Villordo et al., 2016; Liu and Qin, 2020), the 3′-UAR and the 3′-DAR, as well as their complementary partners in the 5′ end, can also be readily recognized.

Interestingly, the local conformations of the 3′-CS, 5′-UAR and 5′-DAR are not universal among the MBFVs. Instead, previous studies have shown that the upstream portion of the 3′-CS forms a pseudoknotted stem with the top loop sequence of the dumbbell (DB) structure in one of the two major phylogenetic branches of the MBFVs (hereafter referred to as the subgroup II, which includes DENV, ZIKV, WNV and many other human/animal pathogenic MBFVs) (Sztuba-Solinska et al., 2013; de Borba et al., 2019; Akiyama et al., 2021; Li et al., 2023). Based on the results in DENV, this pseudoknot is proposed to limit the degree of genome cyclization (Li et al., 2023). By contrast, in the other phylogenetic branch (hereafter subgroup I, including YFV and several lesser-known MBFVs, such as Wesselsbron virus (WESSV)), the 3′-CS is essentially not involved in the formation of the DB’s pseudoknot (Li et al., 2023). The local structure of the 5′-UAR also exhibits a phylogenetically related pattern. In the subgroup I MBFVs, the 5′-UAR is largely single-stranded, whereas it folds into an irregular stem-loop (the 5′-SLB) in the subgroup II MBFVs (Liu et al., 2016; Li et al., 2023). Previous studies suggested that the 5′-DARs in the subgroup II MBFVs are short single-stranded regions locally (Friebe and Harris, 2010; Friebe et al., 2011; Liu et al., 2016; Li et al., 2020). However, the structure and function of the 5′-DAR or its functional equivalent in the subgroup I MBFVs remain largely uncharacterized.

In this study, the YFV 17D vaccine strain was chosen as a model for the subgroup I MBFVs, and through the combination of reverse genetics approaches, RNA structure chemical probing and other biochemical assays, we demonstrated that the hairpin 2 in the capsid-coding region (cHP2), which is localized upstream of the 5′-CS and exists in the linear form of the viral genome, is required for efficient viral RNA (vRNA) replication and viral propagation. Moreover, the results of selective 2′-hydroxyl acylation analyzed by primer extension (SHAPE), SHAPE and mutational profiling (SHAPE-MaP), and electrophoretic mobility shift assay (EMSA) showed that the cHP2 interacts with the 3′-sHP to form a duplexed region in a circularized viral genome, which is stabilized by the cHP2-sHP interaction. This interaction was predicted across multiple members in the subgroup I MBFVs. Thus, the cHP2 can be recognized as the analogue of the 5′-DAR. Interestingly, several atypical phenomena in EMSA and replicon assays led to the discovery that the 3′-sHP is involved in a novel tertiary interaction with the downstream portion of the 3′-CS. This 3′-interlock motif is highly likely to be conserved among all members that form a monophyletic group with the MBFVs and it is suggested to account for a newly revealed negative regulatory mechanism of genome cyclization in these viruses. Our work highlights a delicate communication between specialized secondary structures and a conserved tertiary motif during the life cycle of a major branch of the euflaviviruses.

## Results

### The cHP2 hairpin interacts with the sHP to form a duplex region in a circularized YFV minigenome

In our previous work, two hairpins, the cHP1 and the cHP2, were identified between the 5′-SLB and 5′-CS in the 5′ terminus of the 17D genome by SHAPE-guided RNA structure prediction (Li et al., 2023). This feature was never identified in the subgroup II MBFVs, in which only one hairpin structure, the cHP, was found in the corresponding region (Clyde and Harris, 2006; Clyde et al., 2008; Dong et al., 2008; Polacek et al., 2009; Liu et al., 2016). Interestingly, a prior investigation (Villordo et al., 2010) suggested that the cHP2 is likely to form a long-range interaction with the 3′-sHP structure in the circular form of the 17D genome, which was supported by our previous SHAPE analysis (Li et al., 2023). Accordingly, an early study (Corver et al., 2003) reported that YFV has an extended 5′-CS, and its upstream portion actually overlaps with the cHP2 hairpin. To provide experimental evidence to validate the cHP2-sHP interaction, SHAPE and SHAPE-MaP analyses were performed for the 17D 5′-Ins1BP-312 nt RNA, 3′-UTR RNA, and Ins1BP-minigenome (Figure 1, Figure 1-figure supplement 1 and Figure 1-supplementary files 1 and 2).

**Figure 1.**
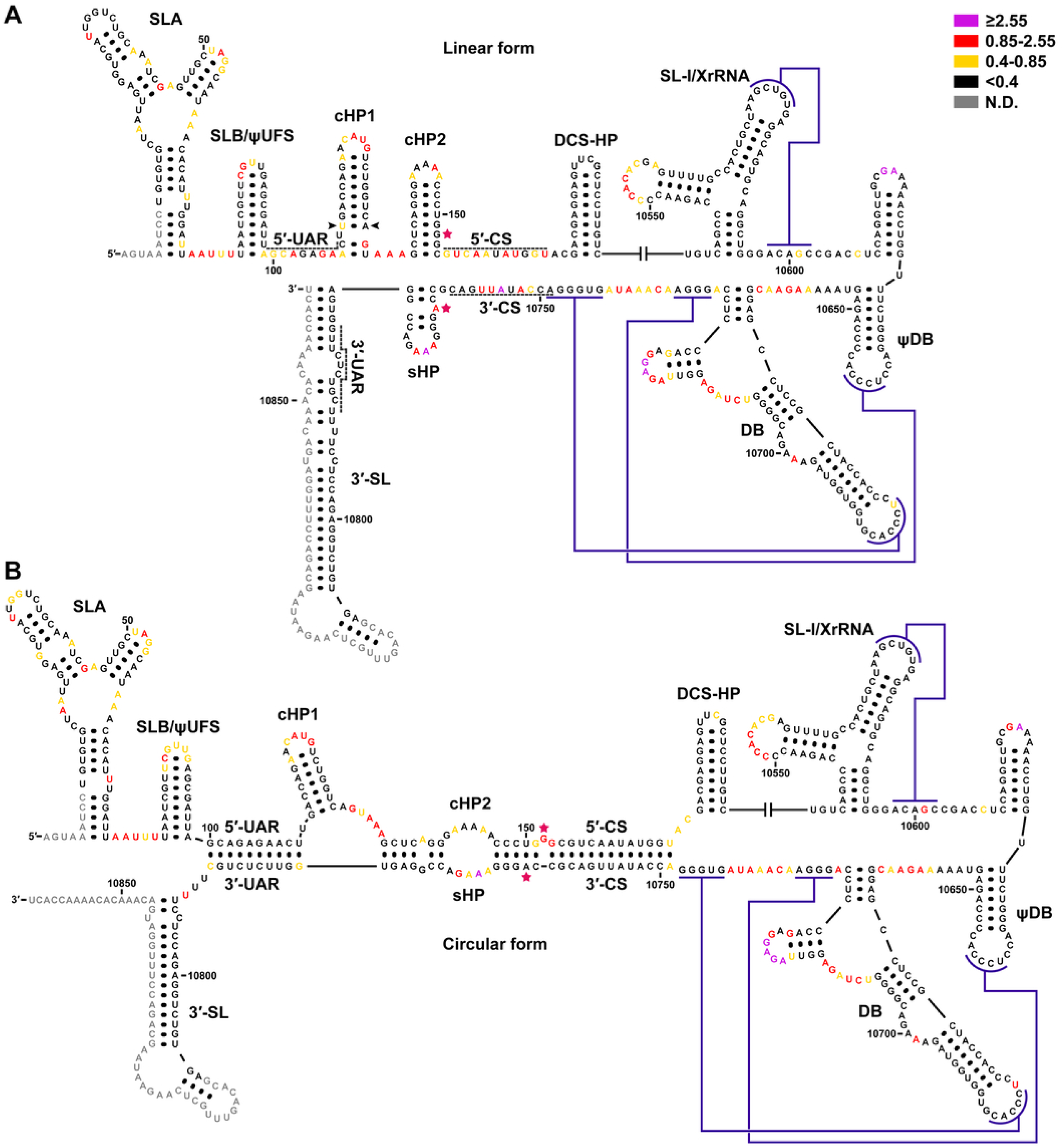
SHAPE analysis of the RNA fragments modeling the linear and circular forms of YFV. (A) SHAPE reactivity values obtained from the 5′-Ins1BP-312 nt RNA and the 3′-UTR RNA were annotated on a linear structure model for the terminal regions of the YFV genome. The base pair inserted into the cHP1 was labeled by arrowheads. (B) The 17D Ins1BP-minigenome was subjected to SHAPE analysis, and the reactivity was annotated on a circular model for the termini of the YFV genome. The RNA structure of the 5′ end in A was generated based on SHAPE-constrained RNA structure prediction by using *RNAstructure* v6.3 (Reuter and Mathews, 2010). For the 3′ end in A, a well-accepted structure model was demonstrated. The structure model in B was generated by SHAPE-constrained RNA structure prediction, except for the xrRNA to DB region, for which their conical structure models were utilized. The shown SHAPE reactivity values were the average of two biological replicates, and the level of SHAPE reactivity was labeled in different colors. Black: <0.4, Golden yellow: ≥0.4 and <0.85, Red: ≥0.85 and <2.55, Purple: ≥2.55, Gray: not determined (N.D.). The numbering was based on the unmodified sequence of the 17D genome. Various RNA structures were labeled and the dark violet lines stand for the pseudoknots. The stars colored in raspberry highlight the nucleotides that exhibited characteristic shifts in SHAPE reactivity during the structural change of the termini of the 17D genome.

We found that the reactivity from both the SHAPE and SHAPE-MaP experiments was in good agreement with the well-established structural model of the 17D 3′-UTR (Figure 1A and Figure 1-figure supplement 1). Then, the reactivity obtained by SHAPE was used to generate RNA structural models for the 5′-Ins1BP-312 nt RNA by using the *RNAstructure* v6.3 package (Reuter and Mathews, 2010). A top-ranked structure similar to that reported previously (Li et al., 2023) was obtained and the cHP2 hairpin was also identified in all suboptimal structures (Figure 1A and Figure 1-supplementary file 1). Next, the Ins1BP-minigenome was analyzed with the reactivity obtained by either SHAPE or SHAPE-MaP as a constraint. Both predictions indicated that the Ins1BP-minigenome is in the circular form, and the panhandle structure formed by long-range RNA-RNA interaction was consistently predicted (Figure 1B, Figure 1-figure supplement 1, Figure 1-supplementary files 1 and 2). Furthermore, the cHP2 and the sHP were shown to form a bulged duplex region with a large internal loop. Importantly, several nucleotides (e.g., G^152^ and A^10763^) exhibited hallmark changes in SHAPE reactivity between the minigenome and the 5′ or 3′ RNAs, consistent with the transitions from paired to unpaired conformations (or *vice versa*). The formation of the cHP2 and its base-pairing relationship with the sHP were conserved in representative strains from all seven established genotypes of YFV (Figure 1-figure supplement 2). These findings suggested functional roles for both the local cHP2 and the cHP2-sHP long-range interactions.

### The local secondary structure of cHP2 is a determinant for efficient YFV propagation

To characterize the function of the cHP2’s local structure, two sets of mutations targeting this hairpin were introduced into the infectious clone of 17D (Figure 2A). Since the cHP2 is localized within the coding sequence of the capsid (C) protein and also participates in long-range RNA-RNA interactions, the mutations were designed with great caution to cause no or minimal effects on the C protein sequence and on the base-pairing relationship between the cHP2 and the sHP. Notably, the G21A and G21A-C31U mutations did not cause any change in the C protein, and RNA structure prediction constrained by SHAPE reactivity showed that the cyclization patterns of these two mutants were essentially the same (Figure 2-figure supplement 1 and Figure 2-supplementary files 1 and 2), with a minor variation in the predicted location of the single G bulge in the cHP2-sHP duplex. However, it is likely that the G^151^-G^152^-G^153^ region in the circular form samples intrinsically mixed conformations, as inferred from the corresponding SHAPE reactivity values (Figure 1-supplementary files 1 and 2 and Figure 2-supplementary file 2). The only obvious difference was that the G21A mutation remodeled and destabilized the cHP2 hairpin, whereas this local structure was restored in the G21A-C31U mutant, as shown by SHAPE-guided RNA structure prediction of the corresponding mutated 5′-Ins1BP-312 nt RNAs (Figure 2-figure supplement 1 and Figure 2-supplementary file 1). The G21A-G22A-A24U, C30U-C31U and G21A-G22A-A24U-C30U-C31U mutants were designed based on a similar strategy. However, this group was designed to have a more profound effect on the disturbance of the secondary structure of the cHP2 (especially for the G21A-G22A-A24U mutation) at the cost of increased, but presumably acceptable influences on the C protein sequence and the cHP2-sHP pairing relationship (Figure 2A and Figure 2-supplementary files 2 and 3).

**Figure 2.**
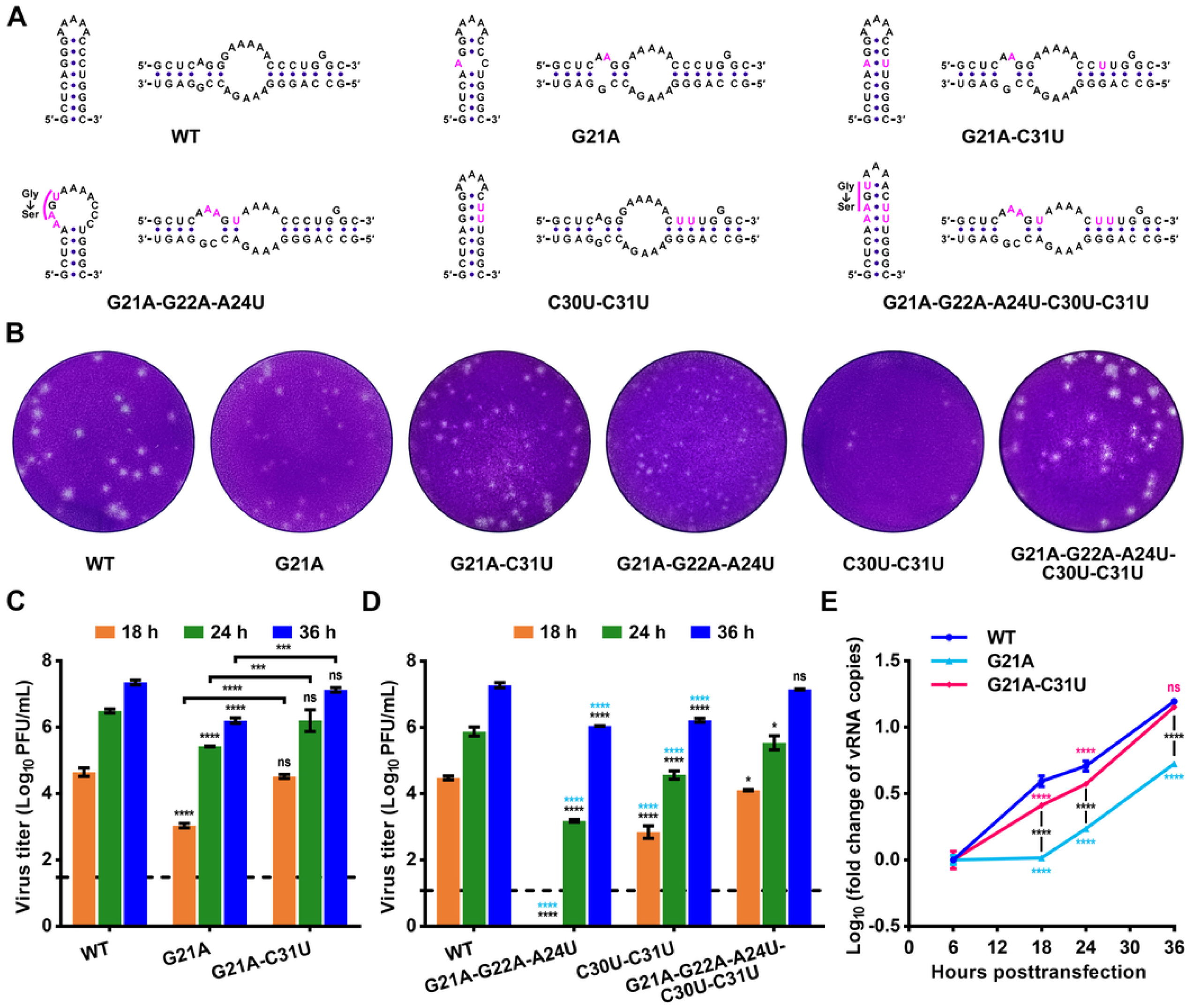
The local hairpin structure of the cHP2 is required for efficient YFV propagation. (A) Design of mutations targeting the local structure of the cHP2. Mutations were highlighted in magenta. The potential effects of the mutations on the complementary relationships between the cHP2 and the sHP were shown for reference. The G21A-G22A-A24U mutation caused a Gly-to-Ser substitution in the C protein sequence. It should be noted that the shown structures were based on a manual inference of the mutations’ effects with reference to the results of *mfold* and *RNAstructure* prediction; they were not meant to represent the most thermodynamically stable ones. Structure demonstration in A was performed with *VARNA v*3.93 (Darty et al., 2009). (B) Plaque morphology of the WT 17D and the cHP2 mutants at 96 h postinfection. (C-D) Titration of the progeny virus in culture supernatants of the vRNA-transfected BHK-21 cells. C and D represented two different experiments that were performed in duplicate. (E) Determination of the efficiency of vRNA amplification in transfected cells by qRT-PCR. The transfection and subsequent experiments were performed in triplicate. The data in C to E were shown as the mean ± SD, and two-way ANOVA and Tukey’s multiple comparisons test were performed. \**P*<0.05, \*\*\**P*<0.001, \*\*\*\**P*<0.0001, ns: not statistically significant. In E, the light blue asterisks for statistical significance indicated the comparison with the G21A-G22A-A24U-C30U-C31U mutant. The numbers in the designations of the cHP2 mutants were determined by the positions in an unmodified C-coding sequence of 17D.

The wild type (WT) and the mutated vRNAs were then transfected into BHK-21 cells. It was shown that the plaques formed by the cHP2-disrupting or destabilizing mutants tended to be smaller and indistinct (Figure 2B). Titration of progeny virus in the culture supernatants demonstrated that the G21A mutant exhibited an apparently lower level of viral propagation compared with the WT and the G21A-C31U mutant at 18, 24 and 36 h posttransfection. By contrast, the G21A-C31U mutant replicated at a similar level to the WT, indicating that reconstitution of the cHP2 hairpin restored the replication deficiency caused by the G21A mutation (Figure 2C). Similar results were obtained from the other panel of mutants. The C30U-C31U mutation, which simultaneously destabilizes the cHP2 hairpin and the cHP2-sHP duplex (Figure 2A and Figure 2-supplementary files 1 to 3), attenuated viral replication, whereas the G21A-G22A-A24U mutant was further attenuated, with no progeny virus detected at 18 h posttransfection. This phenomenon was likely due to its more extensive disruption in local RNA structure and/or long-range RNA interaction, but the glycine-to-serine change in C protein may also contribute to some degree. However, since the replication of the G21A-G22A-A24U-C30U-C31U mutant was substantially more efficient than that of both the C30U-C31U and G21A-G22A-A24U mutants, and only had a modest difference from the WT (Figure 2D), it can be deduced that the replication deficiency of the G21A-G22A-A24U was mainly determined by the disruption/destabilization of the local cHP2 hairpin.

Moreover, cells transfected with the WT, the G21A or the G21A-C31U vRNA were collected at various time points and total RNA was isolated and subjected to quantitative RT-PCR (qRT-PCR). It was found that the vRNA replication of the G21A mutant was also significantly attenuated in comparison with the WT and the G21A-C31U mutant (Figure 2E). In contrast, by monitoring the intracellular metabolism of vRNAs which encode NS5 deficient in RNA-dependent RNA polymerase activity (ΔGDD), we showed that the G21A and the G21A-C31U mutations had no apparent effect on vRNA stability (Figure 2-figure supplement 2). Collectively, the above results demonstrated that the local hairpin structure of the cHP2 plays an important role in YFV RNA replication.

### The function of the long-range base pairing between the cHP2 and the sHP in YFV RNA replication

Next, the role of the long-range interaction between the cHP2 and sHP elements in viral replication was investigated. Because further mutagenesis would be quite likely to cause changes in the C protein sequence, a previously reported IRES-based 17D replicon (Li et al., 2023) was utilized to bypass the restriction on the cHP2 sequence at the viral translational level. A panel of mutants targeting the base-pairing relationships between the cHP2 and the sHP was then generated based on the 17D-Ins1BP-SP-IRES-Rluc-Rep replicon (Figure 3A). By introducing the G21A and the G21A-C31U mutations into the replicon, it was shown that the early replication of the G21A mutant was less efficient than that of the WT, whereas the G21A-C31U mutant replicated at the same level as the WT. Thus, it was further confirmed that the cHP2 hairpin functions as a *bona fide cis*-acting RNA replication element (Figures 3B and 3C). The cHP2-M1C and cHP2-M2C mutants that contain an intact cHP2 secondary structure also replicated more efficiently than the corresponding cHP2-disrupted mutants, the cHP2-M1A and the cHP2-M2A. However, in line with the scenario that the base pairing between these cHP2-intact mutants and the sHP was interrupted, the cHP2-M1C and cHP2-M2C mutants were still attenuated compared with the WT, especially at the early stage (Figures 3B and 3C).

**Figure 3.**
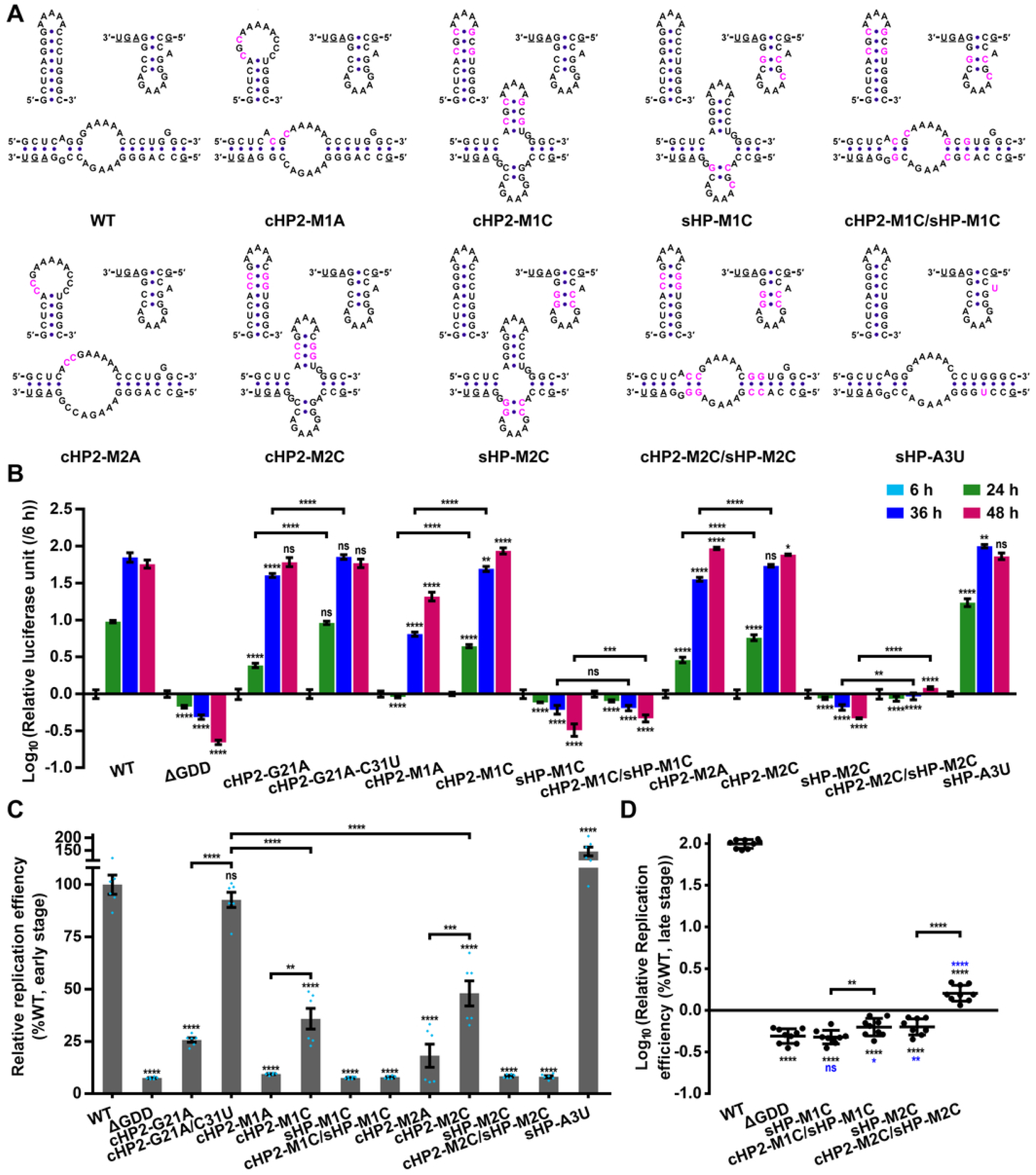
The function of the base pairing between the cHP2 and the sHP in YFV RNA replication. (A) The mutations were highlighted in magenta on the corresponding RNA structures. For each sub-panel, the top left represented the structure of the cHP2, and the structure of the sHP was shown at the top right. The sHP-flanking nucleotides that participate in the interaction with the cHP2 were underlined. The hybridized structure between the cHP2 and the sHP was shown at the bottom. It should be noted that the shown structures of the cHP2/sHP complexes were based on a direct inference of the mutations’ effects; they were not meant to represent the most thermodynamically stable ones. Structure demonstration was performed with *VARNA v*3.93 (Darty et al., 2009). (B) Representative results for the replicon assay. The experiment was performed in triplicate. Data were shown in the logarithmic form of the relative luciferase unit, which was normalized by dividing by the mean raw luciferase activity of the same group at 6 h posttransfection. The error bars stand for standard deviation (SD) in B. Two-way ANOVA and Tukey’s multiple comparisons test were performed. (C) Early replication capacity of the mutants. Data were shown as the relative luciferase unit of a mutant further normalized to the corresponding average value of the WT at 24 h posttransfection (referred to as the relative replication efficiency). Results from two biologically independent experiments, each of which was performed in triplicate transfections, were shown. (D) Replication capacity of the replicons containing mutations in the sHP structure at a late stage of vRNA replication (45 or 48 h posttransfection). Results from three biologically independent experiments, each of which was performed in triplicate transfections, were demonstrated. Data were shown in the logarithmic form of the relative replication efficiency. One-way ANOVA and the two-stage linear step-up procedure of Benjamini, Krieger and Yekutieli were performed for C and D. The error bars stand for standard errors of the mean (SEM) and the individual data points are also displayed. \**P*<0.05, \*\**P*<0.01, \*\*\**P*<0.001, \*\*\*\**P*<0.0001, ns: not statistically significant. Blue asterisks in D indicated the comparisons with the ΔGDD group.

Meanwhile, although mutating the bulged A^10763^ to U did not hinder vRNA replication, the replication capacity of both the sHP-M1C and sHP-M2C mutants was substantially impaired (Figures 3B and 3C). Such results were not fully unexpected, since previous studies (Villordo et al., 2010; Villordo and Gamarnik, 2013) reported that the sHP hairpin is also important for MBFV replication *per se*, and several conserved nucleotides in the loop region of the sHP had been reported to be required for efficient growth of WNV (Davis et al., 2013). The above results further suggested that some specific nucleotides and/or base pairs in the sHP’s stem region are also determinants for its function. Nonetheless, the reconstitution of the cHP2-sHP base pairing resulted in a weak, but consistent recovery of vRNA replication at the late time points (the cHP2-M1C/sHP-M1C versus the sHP-M1C, and the cHP2-M2C/sHP-M2C versus the sHP-M2C, Figures 3B and 3D). The above results provided evidence supporting that the long-range interaction between the cHP2 and the sHP is also required for vRNA replication of YFV.

### The interaction between the cHP2 and the sHP promotes YFV genome cyclization

An *in vitro* EMSA that models the long-range interaction between flavivirus genomic termini (Alvarez et al., 2005b; Dong et al., 2008; Friebe et al., 2012; Liu et al., 2013; Liu et al., 2016; Li et al., 2020; Li et al., 2023) was then utilized to investigate the effects of various cHP2 and sHP mutations on the efficiency of YFV genome cyclization. It was shown that both the cHP2-M1C and the cHP2-M2C mutations reduced the binding efficiency between the corresponding 5′-Ins1BP RNAs and the WT 3′-UTR RNA, and the binding efficiency was restored to a level similar to the WT control group if the corresponding sHP-M1C or sHP-M2C mutation was introduced into the 3′-UTR RNA (Figure 4). These results indicated that the base pairing between the cHP2 and the sHP is required for efficient long-range RNA-RNA interactions between the termini of the YFV genome.

**Figure 4.**
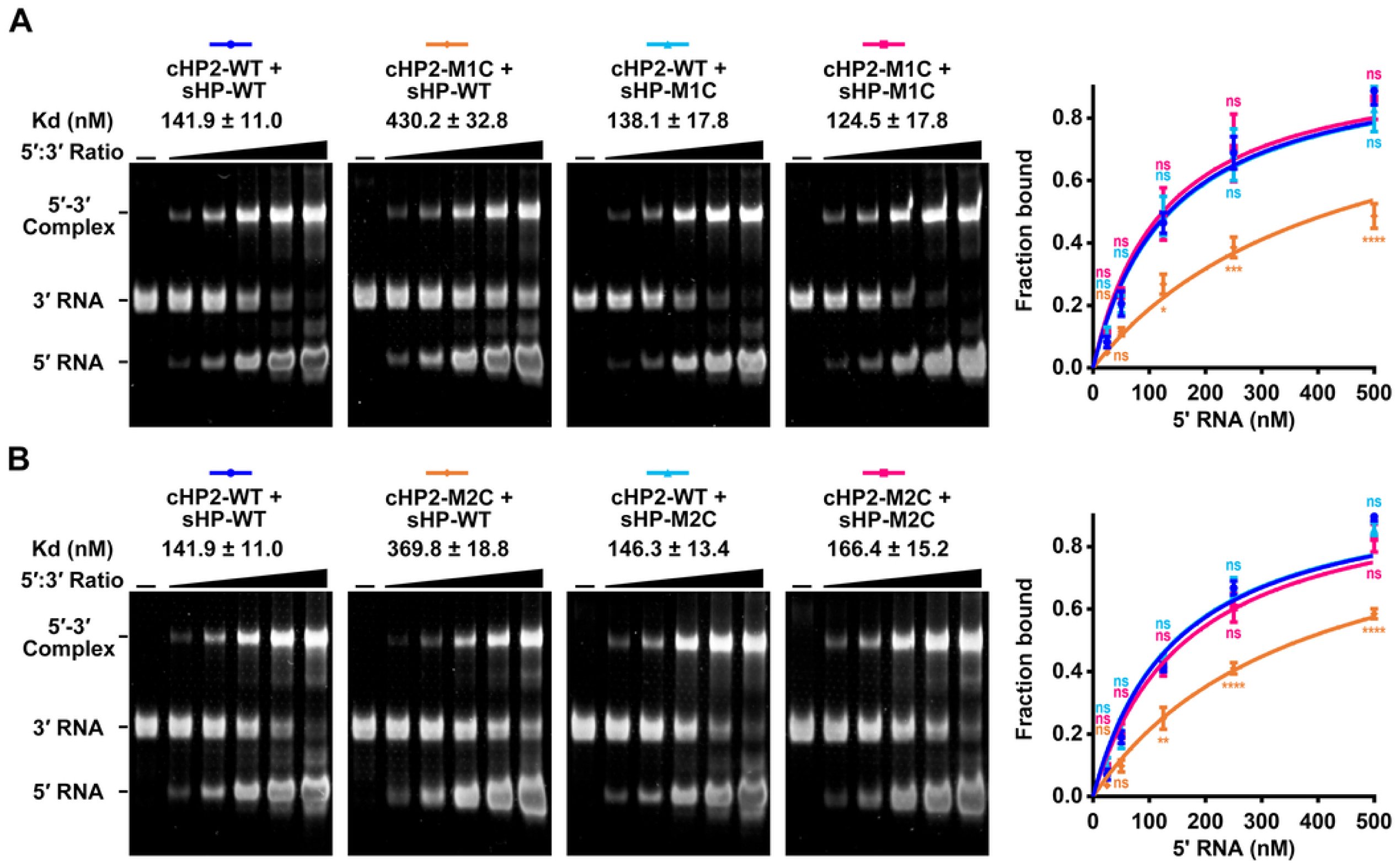
The role of the cHP2-sHP interaction in regulating the binding between YFV 5′ and 3′ RNAs. (A) EMSA for the M1C-series of mutants. (B) EMSA for the M2C-series of mutants. Right: representative results. Left: the summary of three independent biological replicates. Data were shown as the mean ± SEM. Two-way ANOVA and Tukey’s multiple comparisons test were performed for A and B. \**P*<0.05, \*\**P*<0.01, \*\*\**P*<0.001, \*\*\*\**P*<0.0001, ns: not statistically significant. The calculated apparent dissociation constant (*Kd.app*) values were labeled in A. The *Kd.app* for the WT group was calculated from the six biological replicates shown in Figure 4. We noted that additional linear adjustment of contrast and brightness had been made to balance variations between different experiments for demonstration purposes. For more information, please refer to the Materials and Methods section and the source data of Figure 4.

Unexpectedly, the sHP-M1C and sHP-M2C mutations by themselves only slightly affected the 3′-UTR RNA’s binding efficiency with the WT 5′-Ins1BP-312 nt RNA (Figure 4), which was seemingly contrary to the sHP’s well-accepted function in the long-range interactions between MFBV genome termini (Friebe and Harris, 2010; Villordo et al., 2010; Villordo and Gamarnik, 2013; Li et al., 2020). Since a similar phenomenon was consistently observed in these two sHP mutants, it was speculated that there were some hidden mechanisms rather than experimental artifacts behind these results.

### SHAPE analysis suggested that the sHP is likely involved in tertiary interactions

To understand the underlying mechanisms for the results observed by the EMSAs, the Ins1BP-minigenomes containing the “M2C” group of mutations were analyzed by SHAPE (Figure 5A). The SHAPE reactivity of the 5′/3′-CS and 5′/3′-UAR regions was low in general, suggesting that the minigenomes predominantly adopted circularized conformations (Figure 5A and Figure 5-supplementary file 1).

**Figure 5.**
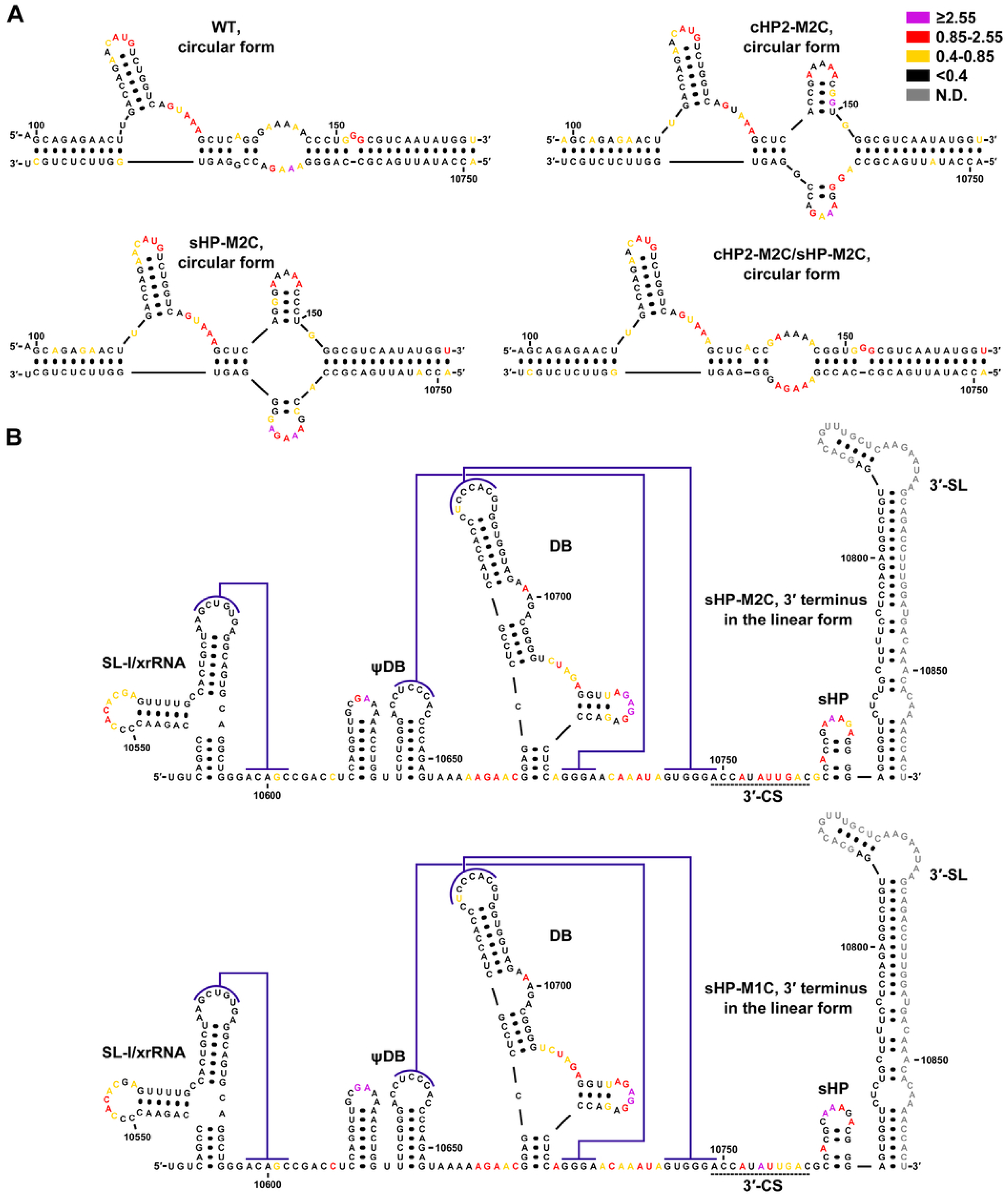
SHAPE analysis suggested that the sHP is involved in a local tertiary interaction. (A) The cHP2-M2C, sHP-M2C and cHP2-M2C/sHP-M2C-containing Ins1BP-minigenomes were subjected to SHAPE analysis, and the obtained SHAPE reactivity was used as constraints for RNA structure prediction. Only the top-ranked predicted structures formed between the various 5′ and 3′ cyclization elements were shown. The averaged SHAPE reactivity from two biological replicates was annotated on the structures, following the rules described in Figure 1. The WT minigenome was demonstrated in parallel and the same SHAPE reactivity in Figure 1B was annotated. (B) The averaged SHAPE reactivity values from two biological replicates were annotated on the structure models for the sHP-M1C and the sHP-M2C-containing 17D 3′-UTRs. The DR regions were not shown due to space limitations. The numbering was based on the unmodified sequence of the 17D genome.

Then, RNA structure prediction for the M2C-set of minigenomes was performed by using the obtained SHAPE reactivity as constraint files. As expected, it was demonstrated that the cHP2-M2C, the sHP-M2C and the cHP2-M2C/sHP-M2C minigenomes were all circularized, as the WT was. From the most stable predicted structures, we found that the cyclization elements in the cHP2-M2C/sHP-M2C mutant formed a panhandle-like structure very similar to that in the WT Ins1BP-minigenome. By contrast, in the cHP2-M2C and sHP-M2C minigenomes, a four-way junction structure was predicted to form between the cHP2 and the sHP (Figure 5A). The four-way junction was nearly identical between the cHP2-M2C and the sHP-M2C mutants, although marginal differences were identified in the folds of the residual sHP structures. Some nucleotides in the stem region of the partially opened cHP2 exhibited moderate to high SHAPE reactivity, suggesting that the structures of the cHP2-sHP complex in these mutants were likely to be flexible and locally heterogeneous. However, the overall folding of these mutated minigenomes was essentially the same as the WT, and no apparent changes in SHAPE reactivity pattern were observed, except for the above-described cHP2-sHP region (Figure 5A, Figure 5-supplementary files 1 and 2). Furthermore, the pseudo-Gibbs free energy changes (pseudo-ΔG) to form the circularized cHP2-M2C and sHP-M2C minigenomes were very close to each other, suggesting that the differences in the stability of the circularized structure cannot account for the unexpected binding efficiency between the WT 5′-Ins1BP-312 nt RNA and the sHP-M2C-containing 3′-UTR RNA. Thus, the reason behind the atypical binding behavior is not the changes in the structure and stability of the circular form, raising the possibility that the sHP mutations may cause structural changes in the corresponding 3′-UTRs.

Then, the 3′-UTR RNAs carrying the sHP-M1C or the sHP-M2C mutation were subjected to SHAPE analysis (Figure 5B, Figure 5-supplementary files 3 and 4). By comparing the SHAPE reactivity patterns of the sHP-M1C and sHP-M2C mutants with that of the WT (Figure 1A and Figure 1-supplementary file 1), no apparent changes of biological significance were found for the direct repeats (DRs), the xrRNA and the ψDB-DB regions (Figure 5B, Figure 5-supplementary files 3 and 4). From the available data (which could not cover the entire 3′-SL), the SHAPE reactivity of the 3′-SL region was also not changed. Moreover, the nucleotides forming the stem region of the sHP also exhibited low SHAPE reactivity, whereas the bulged A^10763^ remained highly reactive. These results indicated that the secondary structures of the sHP-M1C and sHP-M2C mutants are the same as the WT. From the above results, it can be inferred that these mutations changed neither the local secondary structure of the sHP nor the overall folding of the 3′-UTR.

Instead, we noticed that the SHAPE reactivity of some nucleotides in the loop region of the sHP (G^10770^ and A^10771^, referred to as the L5 and L6 nucleotides, or L5 and L6 frequently in the following text) changed from low in the WT to medium/high in the sHP-M2C mutant, and the L6 nucleotide of the sHP-M1C mutant also exhibited a dramatic elevation in SHAPE reactivity. Furthermore, the G^10757^-A^10758^-C^10759^-G^10760^ tetranucleotide (referred to as the GACG motif below), which is immediately upstream of the sHP and is a part of the 3′-CS, exhibited a near-zero level of SHAPE reactivity in the WT 3′-UTR, but the SHAPE reactivity of this region was elevated in both the sHP-M2C and sHP-M1C mutants. For the sHP-M2C mutant, three nucleotides in the GACG motif exhibited medium to high SHAPE reactivity. Since SHAPE reactivity is a reflection of the local dynamics of a nucleotide in an RNA strand, and the changes of SHAPE reactivity in the GACG motif and the sHP loop were identified in both of the two sHP mutants, we deduced that there are unrecognized tertiary interactions between the downstream portion of the 3′-CS and the sHP.

### SHAPE-based characterization of molecular signatures required for the GACG-sHP interaction

To further investigate the presence of the tertiary interaction between the 3′-CS and the sHP, SHAPE-based mutagenesis was employed. First, the sHP-M1C and the sHP-M2C were split into three constituent mutations, the sHP-1.4CG, the sHP-1.5CG and the sHP-L1C. By performing SHAPE analysis of 17D 3′-UTR RNAs containing the above mutations, it was found that multiple nucleotides in the GACG and the sHP shifted from low to medium/high SHAPE reactivity in the sHP-1.5CG-carrying 3′-UTR, whereas the sHP-1.4CG and sHP-L1C mutations had slight-to-modest effects on the SHAPE reactivity of the GACG-sHP region (Figure 6A and Figure 6-supplementary file 1). However, since the sHP-M1C that is composed of the sHP-1.4CG and the sHP-L1C actually caused shifts of SHAPE reactivity of the GACG motif and the L6 position in the sHP’s loop (Figure 5B), the roles of the 1.4 base pair and the L1 position in maintaining the tertiary interaction could not be ruled out. Next, the L2 to L6 positions were individually mutated and SHAPE assays were performed. It was shown that the G^10757^ in the GACG motif exhibited an elevation in the SHAPE reactivity in the sHP-L2C, sHP-L4U and sHP-L6G mutants (Figure 6A), and two additional nucleotides in the sHP-L6G mutant’s stem region also became more reactive to the SHAPE reagent. The sHP-L3G mutant did not cause elevated SHAPE reactivity in the GACG motif and the sHP loop region. In fact, the hyper-reactivity of the L3 position was greatly reduced by this mutation. The most dramatic change in SHAPE reactivity of the GACG-sHP region, however, was caused by the sHP-L5U mutation. In the corresponding mutated 3′-UTR RNA, all four nucleotides in the GACG motif exhibited medium or high SHAPE reactivity, and both the L5 and L6 positions became highly reactive. By examining the conservation of the GACG-sHP region in the mosquito-borne-related flaviviruses (the MBr-FVs, a distinct group including the MBFVs and their close relatives, such as most of the dual host-affiliated insect-specific flaviviruses (dISFVs) and the bat-associated flaviviruses with no known vectors (NKVs)) with available full-length genome sequences, it was clearly shown that the L3 is the only variable position, whereas other nucleotides in the loop region are absolutely conserved (Figure 6-figure supplement 1). The consensus sequence for the YFV GACG motif is GACR, with the first three nucleotides being invariant. Interestingly, although the sequence of the GACG motif is also constrained by the requirement to maintain the base pairing with the 5′-CS, this motif is obviously more conserved than the remaining sequence in the 3′-CS (Figure 6-figure supplement 1).

**Figure 6.**
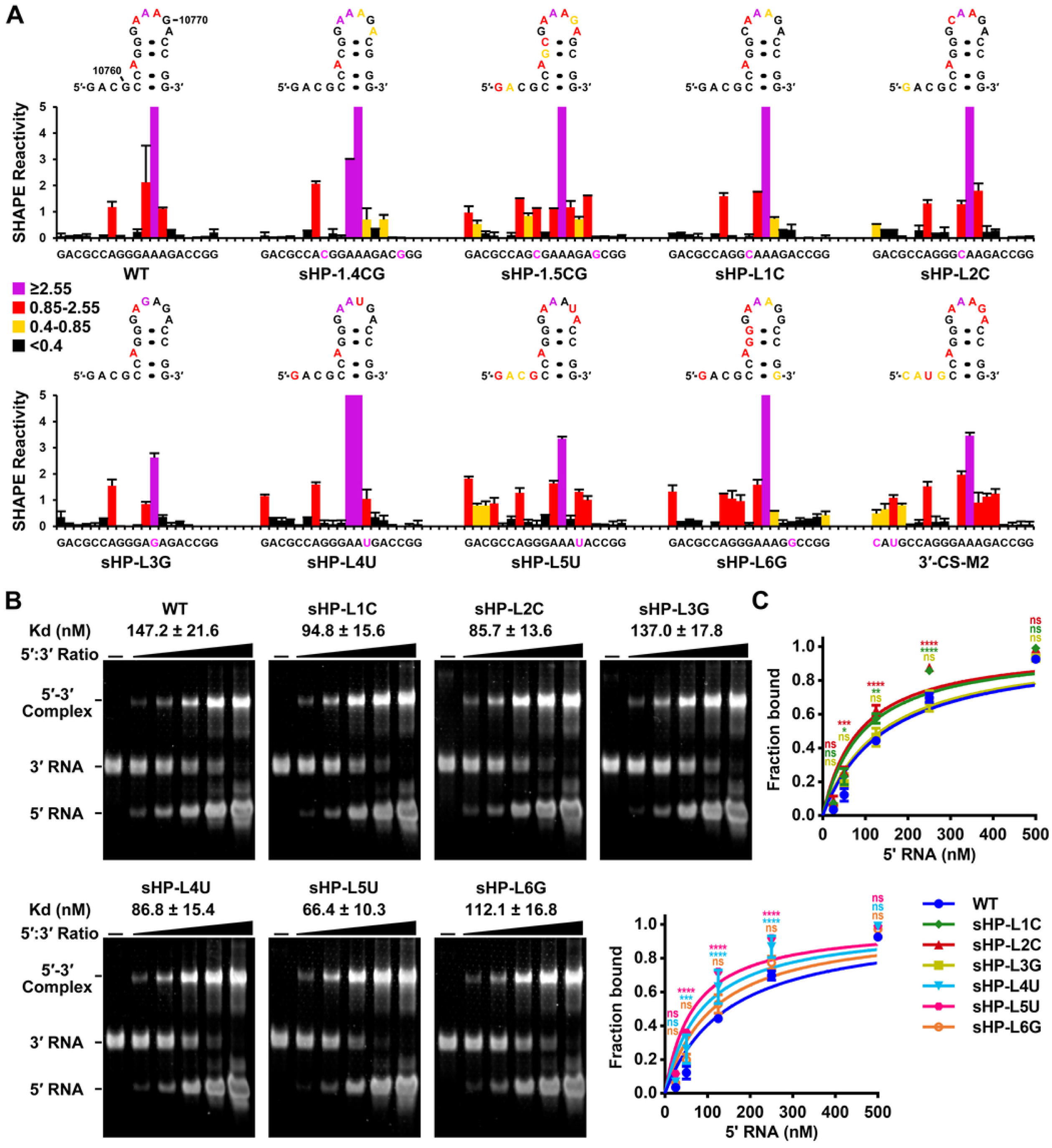
Characterization of nucleotides required for the potential tertiary motif and their effects on *in vitro* binding between 5′ and 3′ RNAs. (A) SHAPE analysis of the mutated 3′-UTRs. Top: SHAPE reactivity was annotated on the corresponding secondary structure. Bottom: SHAPE reactivity for the corresponding region in the upper secondary structure was shown as means plus SD. Mutated nucleotides were highlighted in magenta. Data shown were obtained from two biological replicates. The SHAPE reactivity values of the L3/A^10768^ usually exceeded the range of the Y axis greatly, and the exact numeric values were provided in Figure 6-supplementary file 1 and the corresponding source data. The SHAPE reactivity of the WT group was the same as that in Figure 1A. (B-C) EMSAs were performed to characterize the effects of mutations targeting sHP’s loop sequence on the binding between YFV 5′ and 3′ RNAs. (B) Representative results. (C) Summary of three biological replicates. Data were shown as the mean ± SD. Two-way ANOVA and Tukey’s multiple comparisons test were performed for C. \**P*<0.05, \*\**P*<0.01, \*\*\**P*<0.001, \*\*\*\**P*<0.0001, ns: not statistically significant. The binding reactions for the six mutants and WT were performed in parallel in each of the replicates, and the results were split into two panels for demonstration.

The GACG motif was then mutated into CAUG in the background of the 17D 3′-UTR. It was shown that the mutated tetranucleotide became apparently more SHAPE reactive, and so did the L5 and the L6 positions in the loop of the sHP (Figure 6A, the 3′-CS-M2 group). It has been reported that the loop of the sHP may form a pseudoknot with a pyrimidine-rich internal loop in the bottom to middle region of the 3′-SL in several MBFVs (Shi et al., 1996), but no apparent change in SHAPE reactivity of that internal loop in the 3′-SL was observed in any of the above characterized 17D 3′-UTR mutants (Figure 5, Figure 5-supplementary files 3 and 4, Figure 6A and Figure 6-supplementary files 1 and 2). Notably, the existence of the reported pseudoknot interaction was not consistently supported in subsequent investigations (Davis et al., 2013; Zhu et al., 2024).

The 3′-UTRs of the flaviviruses are well-known to form complicated pseudoknots (Olsthoorn and Bol, 2001; Silva et al., 2010; Sztuba-Solinska et al., 2013; Villordo et al., 2015; Akiyama et al., 2016; Villordo et al., 2016). This feature hinders standard RNA structure prediction based on Zuker’s MFE algorithm (utilized by both the *mfold* and *RNAstructure* programs) (Zuker and Stiegler, 1981; Walter et al., 1994; Zuker and Jacobson, 1998; Mathews et al., 1999; Reuter and Mathews, 2010). In our experience, the secondary structure of the sHP was not successfully predicted in the context of a WT 17D 3′-UTR by using the *RNAstructure* program without additional constraints on the formation of the pseudoknots, even when a SHAPE constraint file was applied. Thus, instead of *de novo* RNA structure prediction, the SHAPE reactivity data of the 3′-UTRs were mapped onto the established model to evaluate the consistency in the above sections. To avoid possible limitations of this strategy, an additional uracil nucleotide was inserted into the WT sHP to constitute a Watson-Crick base pair with the bulged A^10763^. The rationale for this design is that the bulge is not a conserved feature of the sHPs, which usually have a stem region of 4 to 5 base pairs (Figure 6-figure supplement 1). The U-insertion resulted in the sHP’s secondary structure being readily predicted in the sHP-InsU-containing 3′-UTR (Figure 6-supplementary file 3), indicating that this variant can serve as a basis to generate mutants suitable for direct RNA structure prediction. Indeed, the results of SHAPE-guided RNA structure prediction showed that the sHP-InsU hairpin, or its slightly altered version, was present in the top-ranked structures of most mutants, except for the sHP-InsU-L6G-containing 3′-UTR (Figure 6-supplementary files 1 and 3).

Then, the influences of the mutations on the SHAPE reactivity of the GACG-sHP-InsU region were evaluated (Figure 6-figure supplement 2 and Figure 6-supplementary file 1). It was shown that the sHP-InsU-L5U mutation caused a substantial increase in SHAPE reactivity of the GACG motif and the L5-L6 positions, strikingly resembling that of the L5U mutation in the background of the parental sHP. The SHAPE reactivity profile of the sHP-InsU-L6G also exhibited similarity to that of the sHP-L6G, as they shared elevated SHAPE reactivity at the G^10757^ and several positions in the stem region of the sHP. The above phenomenon suggested that the A-to-G mutation in L6 had a destabilization effect on the sHP’s secondary structure, which was also reflected by the above RNA structure prediction (Figure 6-supplementary file 3). The L6 position showed elevated SHAPE reactivity of different degrees in the sHP-1.4CG, the sHP-InsU-1.4CG and the sHP-InsU-1.4UA mutants, and the other nucleotides of the GACG-sHP region in these three mutants had SHAPE reactivity profiles similar to those of the corresponding WT controls, agreeing with a minor role for this base pair in the tertiary interaction (Figure 6A and Figure 6-figure supplement 2). The sHP-InsU-1.1GC and the sHP-InsU-1.2GC mutations did not apparently affect the SHAPE reactivity profiles of the GACG-sHP region, suggesting that their contributions to the higher-order interaction are negligible. Moreover, the 1.5GC base pair may not be the central component to establish the GACG-sHP interaction, as the effects of the corresponding mutations were less profound in the stabilized sHP-InsU compared with those in the parental sHP (Figure 6A and Figure 6-figure supplement 2, sHP-InsU-1.5CG/1.5UA versus sHP-1.5CG). Collectively, an explicit interpretation that the L5 (G^10770^) is an important molecular signature in the sHP for the maintenance of the interaction with the GACG motif can be made, even though a systematic understanding of how tertiary interaction information is reflected by SHAPE reactivity profiles is still lacking.

### The GACG-sHP interaction negatively controls YFV genome cyclization

If the evaluated SHAPE reactivity observed in the sHP loop mutant truly reflected a disruption of the tertiary interaction with the GACG motif, then these mutations would enhance the binding efficiency between the 5′ RNA and the corresponding 3′-UTR RNAs, since many of them were localized in the internal loop of the cHP2-sHP duplex structure and thus had little effect on the base-pairing pattern of the circularized form. Agreeing with this deduction, it was shown that the sHP-L5U-containing 3′-UTR bound to the 5′-Ins1BP-312 nt RNA with the highest efficiency by EMSA, and the sHP-L2C, sHP-L4U and sHP-L6G also enhanced the binding between the 5′-Ins1BP-312 nt RNA and the corresponding 3′-UTR mutants (Figures 6B and 6C). The potential destabilization of the sHP’s secondary structure by the sHP-L6G mutation may result in partial misfolding of the GACG-sHP region, which could explain its less efficient binding to 5′ RNA than that of the sHP-L2C and the sHP-L4U. The sHP-L1C mutation also enhanced the binding efficiency between YFV terminal RNAs to a degree close to the sHP-L2C and the sHP-L4U. Thus, given the combined effects of the sHP-L1C and sHP-1.4CG mutations on the SHAPE reactivity of the GACG-sHP region (Figure 5B and Figure 6A), the L1 position is also likely to participate in the formation of the tertiary motif. By contrast, the sHP-L3G mutation did not apparently affect the binding efficiency. Thus, the results from SHAPE, RNA binding assay, and consensus structure analysis were broadly consistent.

In addition, SHAPE-MaP analyses were performed for the 17D Ins1BP-minigenomes containing the sHP-L3G and sHP-L5U mutations, and RNA structure prediction was performed with SHAPE reactivity as constraints. It was shown that the above two mutants shared essentially the same circularization pattern as the WT (Figure 6-figure supplement 3 and Figure 6-supplementary file 4). From the ranked 1^st^ predicted results, the pseudo-ΔG of the sHP-L5U minigenome was moderately less negative than that of the sHP-L3G and the WT minigenomes, thereby excluding the possibility that the differences in binding efficiency were caused by variations in the structure and stability of the circularized forms. The above data suggested that the GACG motif and the sHP, both of which are essential components of MBr-FV genome cyclization, interact with each other by the formation of a higher-order, “interlock” motif and provide an additional level of control of genome cyclization.

### The sHP-L5/G^10770^ plays a crucial role in YFV RNA replication

A panel of mutations (Figure 7A) that largely overlapped with those in Figure 6A was introduced into the IRES-based 17D replicon and the replication efficiency of the mutants was evaluated in BHK-21 cells. We were unable to generate a replicon carrying the sHP-L6G after repeated attempts, possibly due to the instability of the aimed construct. All the tested mutants showed certain levels of attenuation in vRNA replication except the sHP-L3G, agreeing with the results of SHAPE, EMSA and structural conservation analysis (Figure 6 and Figure 6-figure supplement 1). The sHP-1.4CG mutant exhibited a moderate attenuation characteristic; at 24 h posttransfection, the relative luciferase expression in the cells transfected with this mutant was approximately 40% of that of the WT. By contrast, all the other mutants replicated poorly, with at least a 6.7-fold reduction in relative luciferase expression level at 24 h posttransfection (Figures 7B and 7E). Both the sHP-L1C and sHP-L1A mutants were greatly attenuated, consistent with the suggested participation in the interlock interaction by the L1 position. Importantly, the two mutations targeting the L5 position resulted in the most deleterious effects on vRNA replication (Figures 7B and 7E), and again, this finding was in line with the corresponding results of the SHAPE and EMSA experiments (Figure 6).

**Figure 7.**
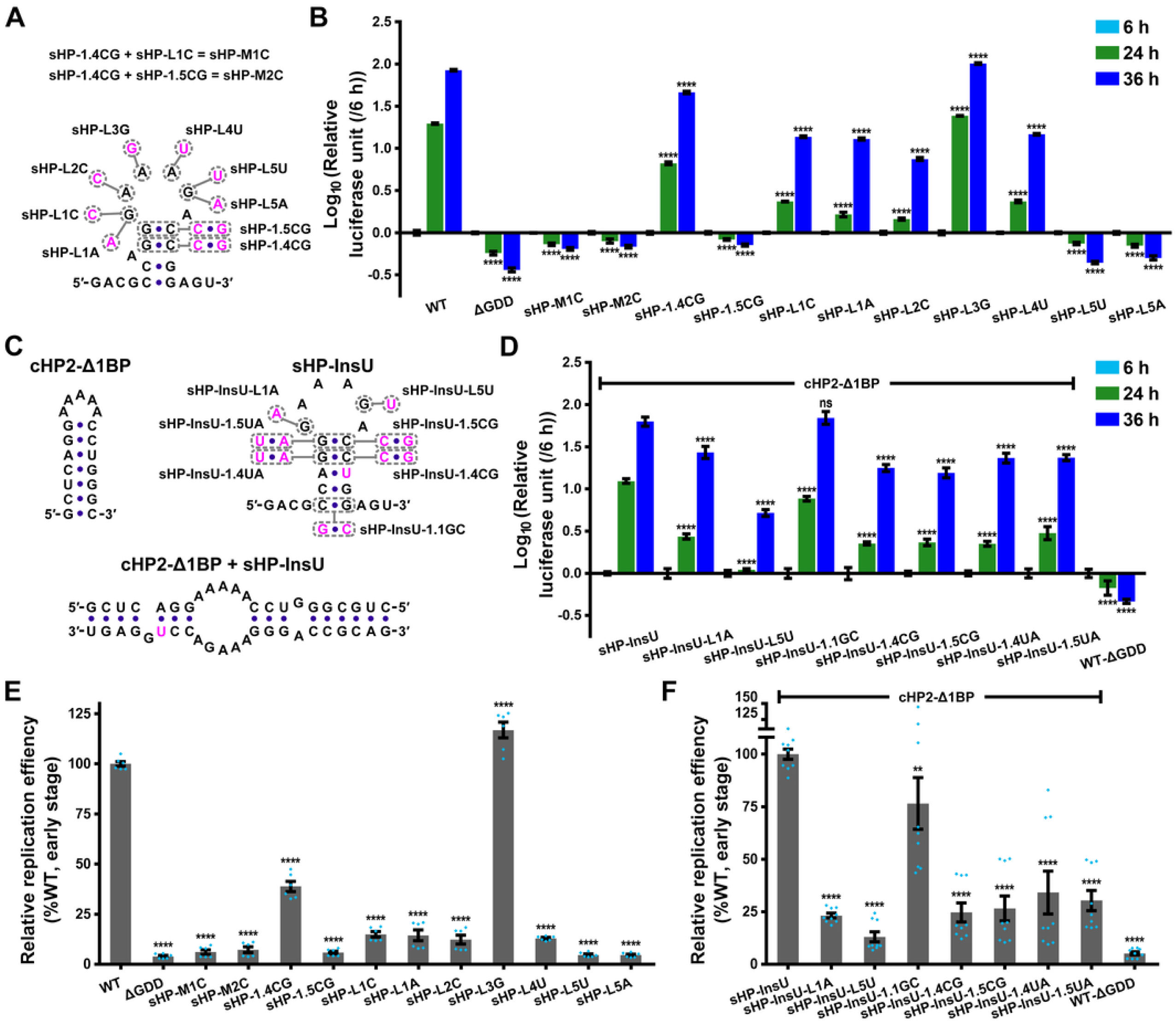
The roles of various structural components of the sHP in YFV RNA replication. (A) Design of the mutations that were introduced into a parental sHP. (B) Representative results of replicon assays for the mutants in A. (C) Design of the mutations that were introduced into a replicon carrying the cHP2-Δ1BP and the sHP-InsU. The duplex formed by the cHP2-Δ1BP and the sHP-InsU was demonstrated according to the prediction results of *mfold* v3.0. (D) Representative results of replicon assays for the mutants in C. Data were shown as the mean ± SD, and two-way ANOVA and Tukey’s multiple comparisons test were performed for B and D. (E) Summary of two independent biological replicates for the mutants in A, each of which was performed in triplicate transfections. (F) Summary of three independent biological replicates for the mutants in C, each of which was performed in triplicate. For E and F, relative replication efficiency calculated at 24 h posttransfection was shown and one-way ANOVA and the two-stage linear step-up procedure of Benjamini, Krieger and Yekutieli were performed. The error bars stand for SEM in E and F and the individual data points are also plotted. \*\**P*<0.01, \*\*\*\**P*<0.0001, ns: not statistically significant. It should be noted that the ΔGDD control used in D and F contained the parental cHP2 and sHP structures.

Mutations were also introduced into a modified version of the 17D-Ins1BP-SP-IRES-Rluc-Rep, which carried a cHP2 shortened by one base pair (cHP2-Δ1BP) and the sHP-InsU variant (Figure 7C). The initial purpose of this design was to maintain the balance between linear and circular forms of vRNA, although these two variants actually had only slight effects on vRNA replication (Figure 7-figure supplement 1). Most tested mutants were attenuated (Figures 7D and 7F), except for the sHP-InsU-1.1GC mutant, which only replicated slightly less efficiently than the control replicon (cHP2-Δ1BP/sHP-InsU). This result was also consistent with its SHAPE reactivity profile, and the slight reduction of vRNA replication was likely caused by the moderate interference with the base pairing between the cHP2 and the sHP. The mutations targeting the 1.4 and the 1.5 base pairs of the sHP reduced vRNA replication by approximately 3- to 4-fold, but the detailed effects somehow varied from the results of the mutagenesis based on the native YFV sHP (Figures 7B and 7E). However, since the effects of mutations targeting the sHP stems that also directly participated in the long-range RNA-RNA interactions were almost unavoidably mixed, the most appropriate interpretation for the above data should be that the 1.4 and 1.5 pairs are involved in the stabilization of the interlock interaction, but not by providing the directly interacting moieties. Importantly, the mutation targeting the L5 position caused the largest degree of attenuation (Figures 7D and 7F), highlighting the crucial role of the G^10770^ in the establishment of the interlock motif.

### Further evidence supporting the existence of the interlock motif was revealed by differential SHAPE analysis

SHAPE reagents, such as N-methylisatoic anhydride (NMIA) and 1-methyl-7-nitroisatoic anhydride (1M7), are thought to have little nucleotide-biased modification (Wilkinson et al., 2006; Mortimer and Weeks, 2007; Busan et al., 2019). However, to exclude the possibility that the above SHAPE results based on NMIA modification were caused by any potential sequence-biased modifications, the WT and the 3′-CS-M2-mutated 17D 3′-UTR RNAs were modified by three different SHAPE reagents, 1M7, NMIA and 1-methyl-6-nitroisatoic anhydride (1M6), and the modification was detected by SHAPE-MaP (Siegfried et al., 2014; Smola et al., 2015; Busan and Weeks, 2018). The WT GACG and the sHP L5-L6 sites were unreactive to all of the three reagents, resulting in low SHAPE reactivity (<0.4) for these positions. By contrast, in the 3′-CS-M2 mutant, most of the corresponding nucleotides exhibited medium to high SHAPE reactivity regardless of the reagents used (Figure 8A, Figure 8-figure supplement 1 and Figure 8-supplementary file 1). Meanwhile, the sHP-L5U-mutated 3′-UTR and two mutants targeting the pyrimidine-rich internal loop of the 3′-SL (3′-SL-M1 and 3′-SL-M2) were modified with 1M7 followed by SHAPE-MaP analysis (Figure 8-figure supplement 1). It was shown that the SHAPE reactivity profile of the sHP-L5U mutant obtained by SHAPE-MaP matched the corresponding data of SHAPE, whereas the mutations in the 3′-SL’s internal loop did not induce any notable changes in the SHAPE reactivity profiles of the GACG-sHP region. These results further demonstrated that the formation of the GACG-sHP interlock does not require the participation of the pyrimidine-rich internal loop in the 3′-SL.

**Figure 8.**
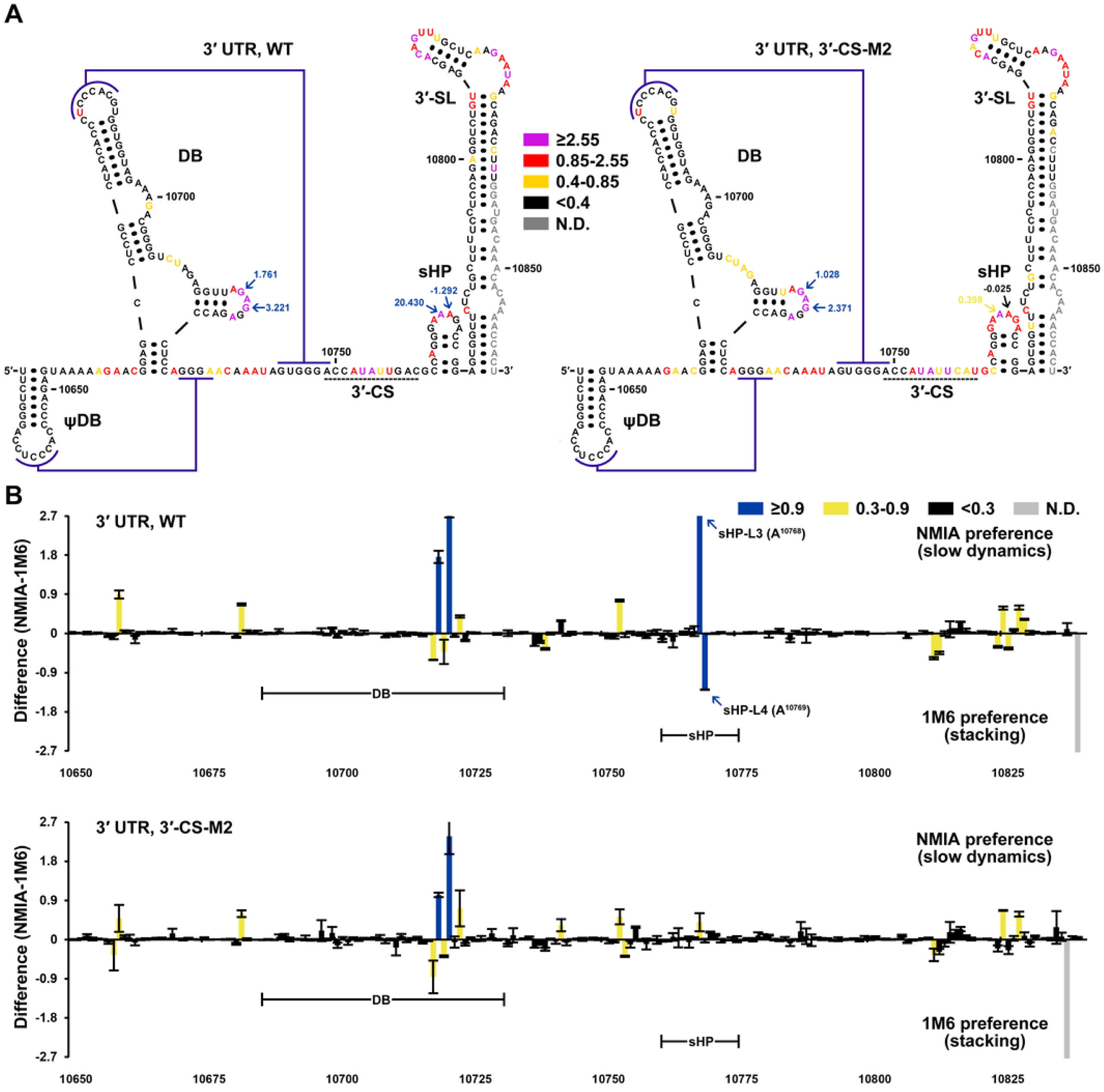
Differential SHAPE reveals a crucial role of the GACG motif in the maintenance of the interlock interaction. (A) Secondary structures of the WT and 3′-CS-M2-containing 3′-UTR RNAs that were annotated with SHAPE reactivity obtained by 1M7 modification and SHAPE-MaP. Only the ψDB to 3′-SL regions were shown for clarity. The dark violet lines indicate the pseudoknot interactions. The nucleotides with a strong differential signal (with an absolute value ≥ 0.9) in the WT RNA were annotated with the values in blue letters. The differential SHAPE values of the corresponding positions in the 3′-CS-M2 RNA were also annotated for comparisons. (B) Differential SHAPE profiles of the WT and 3′-CS-M2-containing 3′-UTR RNAs. Only the data for the 10650-10840 nt region (based on the sequence in the parental 17D genome) were shown. Two biologically independent experiments were performed and the differential SHAPE profiles were generated individually by using the differential SHAPE pipeline (Rice et al., 2014a). The data were shown as the mean ± SD. For the colors of the differential SHAPE values, the following rules were applied. Black: <0.3; Maize yellow: ≥0.3 and <0.9; Cobalt blue: ≥0.9. Gray: N.D. The numerical comparison was based on the absolute values of the differential SHAPE reactivity. The corresponding regions of the DB and the sHP were labeled and the signals for the L3/A^10768^ and L4/A^10769^ in the WT were indicated. The differential SHAPE values in A were based on the results shown in B. The complete differential SHAPE profiles of these RNAs were provided in Figure 8-supplementary file 3.

Differential SHAPE analysis, which is a powerful tool for the hunt of RNA tertiary interactions (Steen et al., 2012; Rice et al., 2014b), was then performed. Three nucleotides were found to have a high NMIA preference (≥0.9) within the core functional region (the xrRNA-ψDB-DB-3′-CS-sHP-3′-SL region) of the WT 3′-UTR (Figure 8 and Figure 8-supplementary file 2). Two of them were localized within the loop region of the small stem-loop in the DB structure. The site with the highest NMIA preference among the three was the L3/A^10768^ of the sHP. Meanwhile, the L4/A^10769^ was the only nucleotide with a high 1M6 preference (≤-0.9) in the WT 3′-UTR. In the 3′-CS-M2-mutated 3′-UTR, the two nucleotides in the DB still had high NMIA preferences. Interestingly, the 1M6 preference of the L4/A^10769^ completely vanished, and the NMIA preference of the L3/A^10768^ was also greatly reduced in the 3′-CS-M2 (Figures 8A and 8B). Since the 3′-CS-M2 mutation did not cause direct changes in the loop sequence of the sHP and thus avoided the possibility of sequence biases in chemical modification, the above findings demonstrated that the GACG motif is able to affect the conformation of the sHP’s loop, consistent with the existence of the interlock motif. Especially, a high 1M6 preference had been shown to be a property of a nucleotide that participates in a stacking interaction using one side of its nucleobase (Steen et al., 2012; Rice et al., 2014b). For the L4/A^10769^, such a base stacking is more likely to be established with the neighboring L5/G^10770^, which is conformationally constrained by the interlock interaction.

To further validate the above finding, the WT Ins1BP-minigenome and 5′-Ins1BP-312 nt RNA were subjected to differential SHAPE analysis (Figure 8-figure supplements 2 and 3 and Figure 8-supplementary file 3). Indeed, it was demonstrated that the differential SHAPE values of the L3/A^10768^ and the L4/A^10769^ were reduced by more than 10-fold in the minigenome in comparison with those in the free 3′-UTR, whereas the G^10719^ and G^10721^ in the DB retained their high NMIA preferences, confirming that the transition in differential SHAPE characteristics of the L3/A^10768^ and the L4/A^10769^ was determined by specific changes in higher-order RNA structures.

Other than the sites presented above, few nucleotides exhibited a notable preference in reactivity. Among these, the U^10682^ exhibited a high NMIA preference in the circular form (Figure 8-figure supplements 2 and 3). Accordingly, this site was predicted to form the closing base pair of the DB-PK (the pseudoknot formed between the top-loop of the DB and downstream sequence) in the linear form, but became single-stranded after genome cyclization. Only the G^103^ in the 5′-UAR was found to exhibit a relatively stable high NMIA preference in the free 5′-Ins1BP RNA, and this preference virtually diminished in the minigenome (Figure 8-figure supplements 2 and 3 and Figure 8-supplementary file 3), echoing its structural transition during the circularization of YFV RNA. There were also a few sites of high 1M6 preference found in the minigenome (Figure 8-supplementary file 3). However, since their 1M6 preferences were not as strong as that of the L4/A^10769^ (in the free 3′-UTR) and these sites were not relevant to the scope of this study, they were not further investigated.

### The GACG motif is crucial for YFV RNA replication beyond its participation in the long-range base pairing during genome cyclization

Since the importance of the loop of the sHP in YFV RNA replication was demonstrated by replicon assays (Figure 7), we then investigated the role in vRNA replication of the GACG motif, the other component of the interlock interaction. Two pairs of mutants targeting different regions of the 5′-CS/3′-CS duplex were generated (Figure 9A). It was shown that the disruption of the base pairing between 5′-CS and 3′-CS severely impaired vRNA replication (Figure 9B, 5′-CS-M1 and 3′-CS-M2), and the introduction of complementary mutations into the 5′-CS-M1 mutant restored vRNA replication to a similar level to the WT. However, the defect in replication of the 3′-CS-M2 targeting the GACG motif was unable to be rescued by introducing complementary mutations in the 5′-CS (Figure 9B, 3′-CS-M2 and 5′-CS-M2/3′-CS-M2). We further showed that the circularization pattern of the 5′-CS-M2/3′-CS-M2 could actually be restored in the background of the Ins1BP-minigenome by SHAPE and RNA structure prediction (Figure 9-figure supplement 1). Collectively, these results highlighted that the GACG motif plays a critical role in YFV RNA replication independent of its direct participation in genome cyclization.

**Figure 9.**
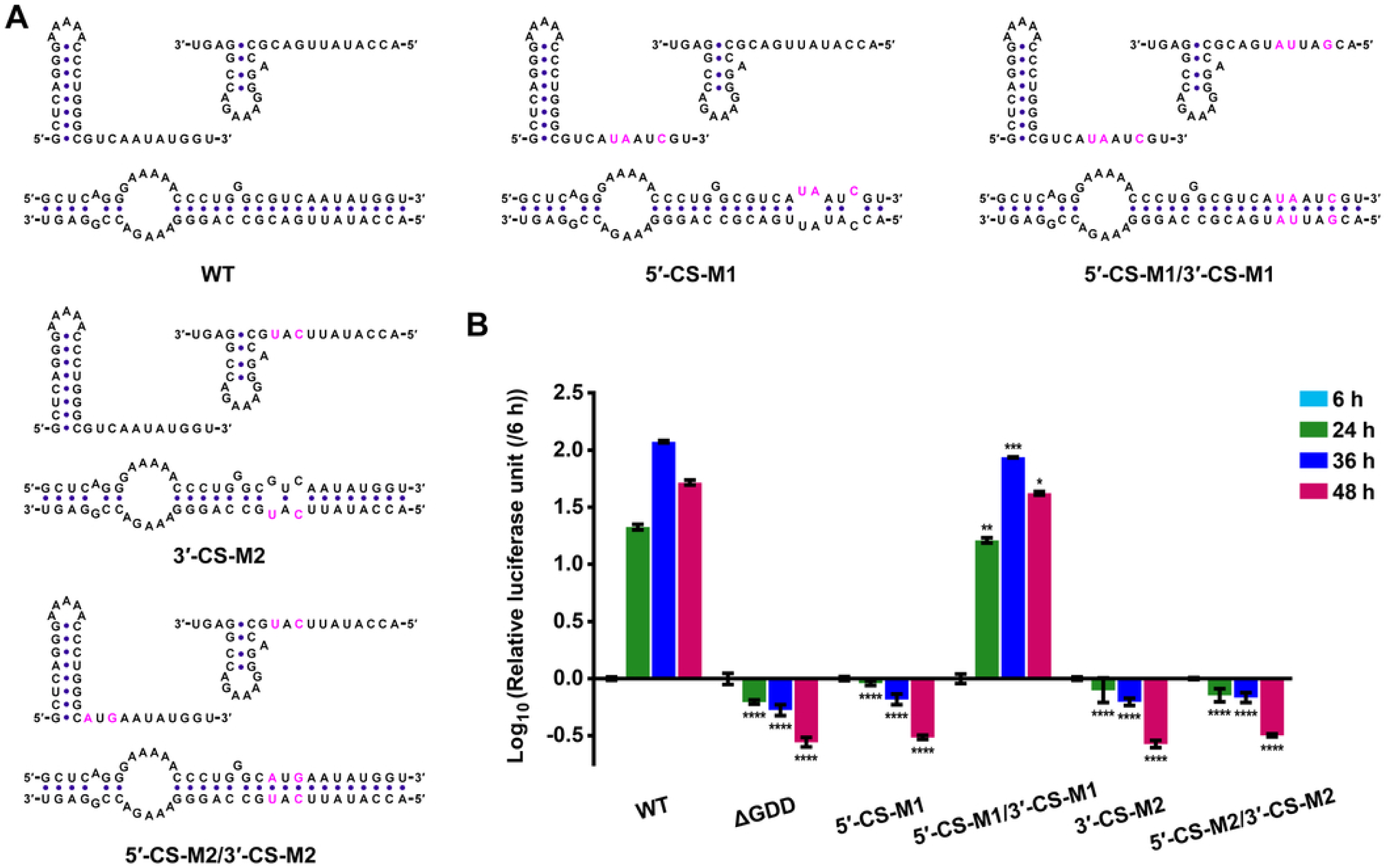
The GACG motif is crucial for YFV RNA replication independent of its direct role in genome cyclization. (A) Design of mutations targeting the 5′- and 3′-CS regions. The mutated sites were highlighted in magenta, and the directly inferred effects on the interaction between the 5′-CS and the 3′-CS were displayed. (B) Results of replicon assay. Data were demonstrated as in Figures 3B, 7B and 7D. Two-way ANOVA and Tukey’s multiple comparisons test were performed. \**P*<0.05, \*\**P*<0.01, \*\*\**P*<0.001, \*\*\*\**P*<0.0001.

### The significance of the interaction between the cHP2 and the sHP was further illustrated by abolishing the interlock motif

As the interlock tertiary motif provides a restriction of the genome cyclization event, if it is abolished, the disruption of the cHP2-sHP base pairing by targeting either secondary structure would have negative effects of similar degree on the *in vitro* binding between 5′ and 3′ RNAs. Based on the available structural information of the interlock motif, a modified sHP was designed. It was shown that the SHAPE reactivity of the GACG region and the L1, L5, L6 positions in the loop of the sHP-PxP (for Pair-cross-Pair) increased to medium or high levels (Figure 10A), in contrast to the low SHAPE reactivity of these nucleotides in the WT GACG-sHP region (Figure 6A). In addition, to avoid the formation of mutation-caused alternative 5′-3′ complex structures, the sHP-PxP was engineered so that each base pair of it would form two base-pairings with a base pair of the cHP2-Δ1BP. On this basis, mutants targeting the base pairing between the cHP2-Δ1BP and the sHP-PxP were further generated (Figure 10B). By EMSAs, we found that the binding between the cHP2-Δ1BP-containing 5′-Ins1BP RNA and the sHP-PxP-containing 3′-UTR RNA was considerably higher than that between the parental RNAs, in agreement with the absence of the interlock motif. Moreover, the introduction of mutations that disrupted the long-range base pairing between the cHP2-Δ1BP and the sHP-PxP into either local structure had comparable inhibitory effects on the efficiency of RNA binding (Figure 10C). As expected, the 5′-Ins1BP-cHP2-Δ1BP-M RNA and the 3′-sHP-PxP-M UTR RNA bound efficiently, with a *Kd.app* value nearly equivalent to that of the corresponding WT RNAs (Figure 10C). These results not only supported the presence and significance of the GACG-sHP interlock motif, but also highlighted the role of the cHP2-sHP interaction in YFV genome cyclization.

**Figure 10.**
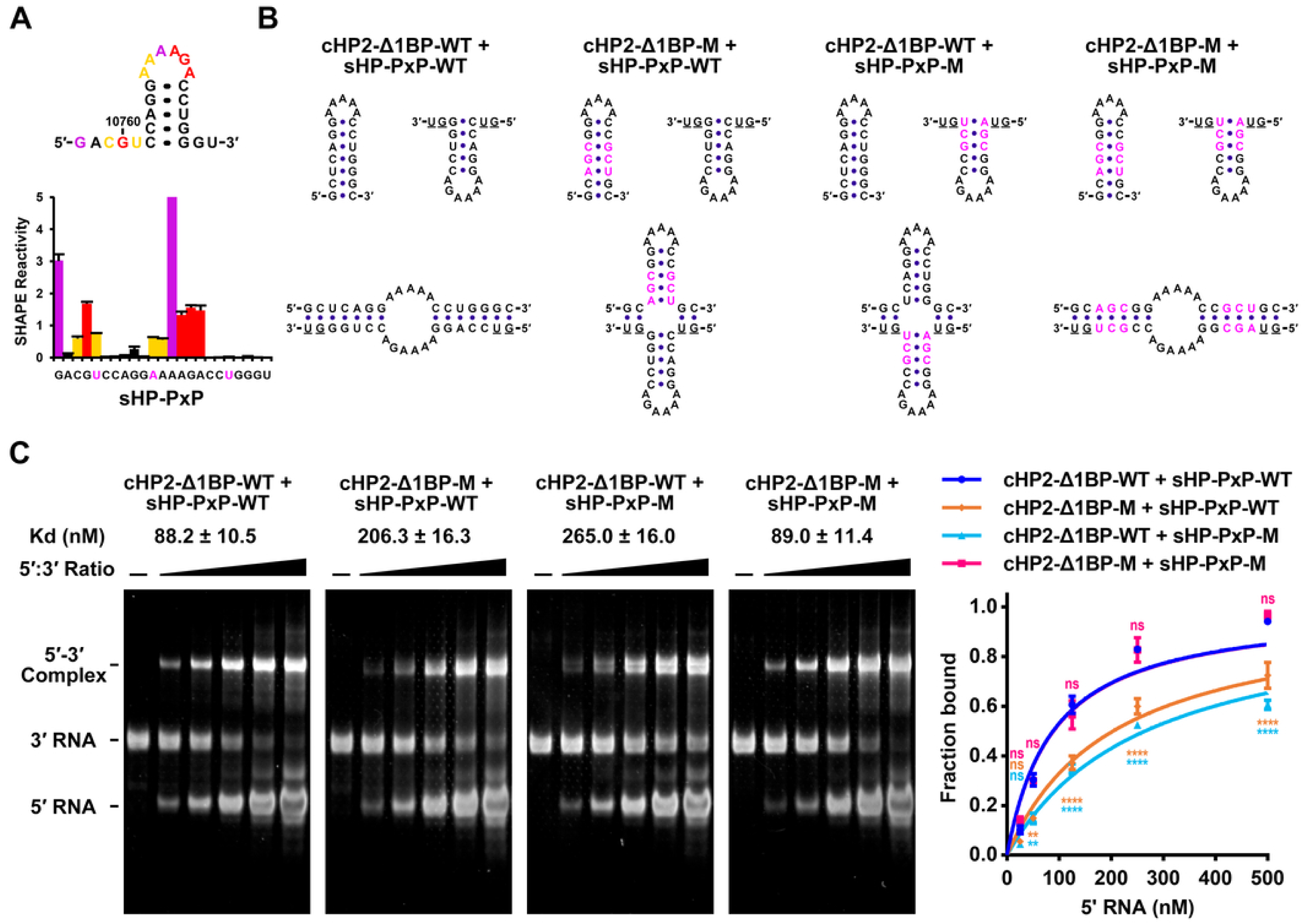
RNA binding assays performed under a condition where the interlock motif was disrupted. (A-B) Design of the cHP2-Δ1BP/sHP-PxP interacting pair. (A) SHAPE reactivity profile of the sHP-PxP variant. The insertion of a U after G^10760^ and the mutation of L1/G^10766^ into an A were aimed at disrupting the interlock motif, as evidenced by the substantial increase in the SHAPE reactivity of the GACG and the L1, L5, and L6 positions. (B) In combination with the above two mutations, the deletion of A^10777^ immediately downstream of the sHP further created a pair-cross-pair relation between the sHP-PxP and the cHP2-Δ1BP structures. Then mutations (highlighted in magenta) targeting the engineered interaction were introduced into the cHP2-Δ1BP or the sHP-PxP. We noted that the shown structures of the cHP2/sHP complexes were for illustration purposes mainly and were not necessarily meant to represent the most thermodynamically stable ones, especially for the interaction-disrupting mutants. (C) Left panel: representative results of the EMSA experiments. Right panel: summary of the results of three biological replicates. Data were shown as the mean ± SEM. Two-way ANOVA and Tukey’s multiple comparisons test were performed. \*\**P*<0.01, \*\*\*\**P*<0.0001, ns: not statistically significant. The calculated *Kd.app* values were shown in the left panel.

### Distribution of the cHP2 and the GACG/sHP-interlock among the euflaviviruses

The phylogenetic distribution of *cis*-acting RNA elements varies across flaviviruses (Villordo et al., 2016; Liu and Qin, 2020). We constructed a phylogenetic tree spanning most available members of the euflaviviruses (Figure 11A and Figure 11-figure supplement 1). The phylogenetic tree clearly showed that the euflaviviruses diversified into two major clusters: the first corresponded to the MBr-FVs and included the MBFVs, two clades of dISFVs and the bat-associated NKVs; while the second was composed of the TBFVs, the rodent-associated NKVs and the recently reported potential tick-specific flaviviruses (Harima et al., 2021; Wang et al., 2024). Consistent with previous works (Alkan et al., 2015; Moureau et al., 2015; Colmant et al., 2016), the ecological characteristics did not precisely reflect the phylogenetic positions of flaviviruses. Therefore, we referred to the second major group as the TBr-FVs (tick-borne-related flaviviruses), in correspondence with the MBr-FVs. The phylogenetic analysis also confirmed that there are two sub-lineages in the MBr-FVs. They are referred to as subgroups I and II MBr-FVs, since these sub-lineages represent expansions of the MBFV branches mentioned above.

**Figure 11.**
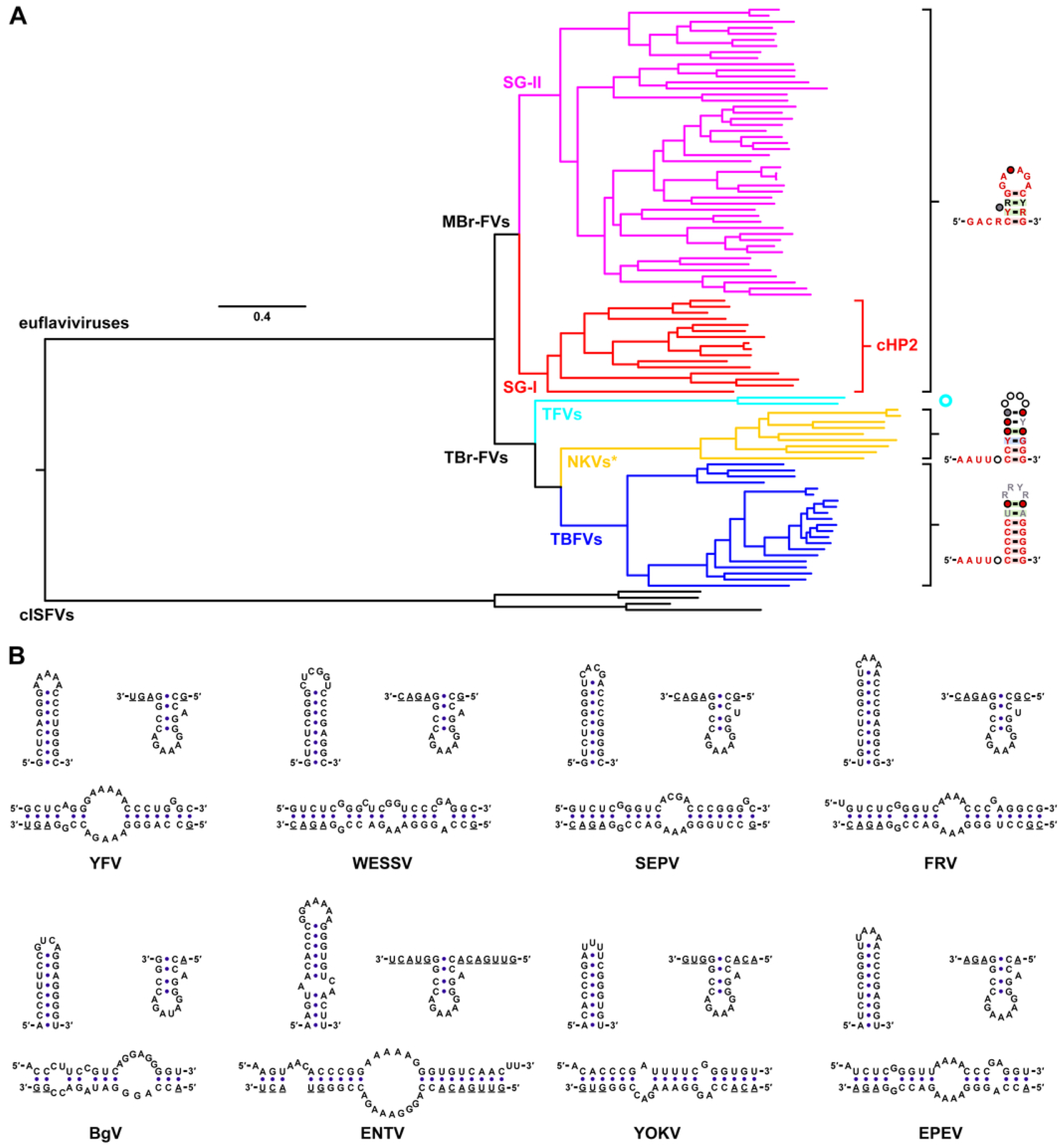
Conservation of the cHP2 and sHP/interlock among different lineages of the flaviviruses. (A) The cHP2 is only present in the subgroup I (SG-I, red) MBr-FVs, but not in the subgroup II (SG-II, magenta) MBr-FVs. The GACR-sHP interlock motif is highly conserved in the entire MBr-FV lineage, regardless of the viruses’ natural host-range, whereas a stem-loop which lacks both an upstream GACR motif and a conserved hexaloop sequence is found in the homologous position in the 3′-UTRs of the TBr-FVs, which include true TBFVs (blue), the rodent-associated NKVs (the NKVs* clade in yellow), and the potential tick-specific flaviviruses (TFVs, highlighted in cyan). The consensus RNA structures were generated by using R2R v1.0.6 (Weinberg and Breaker, 2011). For the rodent-associated NKVs, the upstream AAUU sequence was manually aligned. The bat-associated NKVs, which clustered within the subgroup I MBr-FVs, were highlighted in red. The sHP structures of two “tick-specific” flaviviruses with available 3′-UTR sequences were provided in Figure 11-figure supplement 1. The phylogenetic tree was constructed with the complete ORF sequences of the flaviviruses by using IQ-TREE2 (Lanfear et al., 2020). Representative cISFVs were included as the outgroup to root the tree. (B) Potential patterns of interaction between the cHP2/cHP2-like hairpins and the sHP among subgroup I MBr-FVs with at least one complete genome sequence available. The most stable RNA structures predicted by *RNAstructure* v6.3 for the cHP2/sHP duplex were demonstrated, except for the Bamaga virus (BgV), for which additional constraints were applied in the RNA prediction parameters to generate a duplex structure with all the possible base pairs formed between the cHP2 and the sHP. The structures were illustrated by using *VARNA* v3.93 (Darty et al., 2009). Abbreviations: WESSV, Wesselsbron virus; SEPV, Sepik virus; FRV, Fitzroy River virus; ENTV, Entebbe bat virus; YOKV, Yokose virus; EPEV, Ecuador Paraiso Escondido virus.

Then the presence of the cHP2 and the interlock motif was examined. The cHP2 was found in all members with available complete genome sequences in the subgroup I MBr-FVs, including true MBFVs, e.g., YFV, WESSV, Sepik virus (SEPV), Fitzroy River virus (FRV), as well as several dual-host-affiliated members, Bamaga virus (BgV), Ecuador Paraiso Escondido virus (EPEV), Entebbe bat virus (ENTV) and Yokose virus (YOKV) (Figure 11B, top right in each panel). Since there were only complete ORF sequences for a branch of the subgroup I MBr-FVs (Grard et al., 2010), and these viruses (e.g., Edge Hill virus, Banzi virus, etc.) likely express a N-terminal-truncated C protein (Grard et al., 2010), the presence of a cHP2 in these flaviviruses remains inconclusive. However, it is clear that a cHP2-like element is not present in the subgroup II MBr-FVs, in which a single cHP is flanked by the linear/unstructured 5′-DAR and 5′-CS, as shown by a consensus structure prediction by *LocaRNA* (Figure 11-supplementary file 1) and several previous studies (Clyde et al., 2008; Liu et al., 2016; Liu and Qin, 2020).

On the contrary, the primary sequence for the constitution of the GACG/sHP-interlock motif was identified in all members of the MBr-FV group with full-length genome sequences available (Figure 6-figure supplement 1). As a proof of principle, DENV4 WT and sHP-L5U-mutated 3′-UTRs were subjected to SHAPE-MaP, and the SHAPE reactivity patterns of the GACG-sHP regions in the two RNA species resembled the corresponding 3′-UTRs of 17D (Figure 8-figure supplement 1 and Figure 11-figure supplement 2), the sHP-L5U mutation also resulted in a substantial attenuation in a DENV4 replicon assay (Figure 11-figure supplement 3). These data supported the idea that the GACR-sHP interlock motif is highly conserved among virtually all MBr-FVs. Interestingly, the lack of a GACG motif upstream of the sHP and the distinct pattern of its loop sequences in the TBr-FV group strongly argued that the interlock motif does not exist in these viruses (Figure 11A and Figure 11-figure supplement 1), and this interpretation was supported by the SHAPE-MaP and differential SHAPE analysis of a synthetic 3′-UTR of tick-borne encephalitis virus (TBEV), which showed that all of the loop nucleotides of the TBEV sHP were reactive to the SHAPE reagents with no high differential SHAPE values identified (Figure 11-figure supplements 4 and 5). The above findings highlighted the diversification of *cis*-acting structural elements and regulatory mechanisms of viral replication among the euflaviviruses. Finally, by using conventional single-sequence RNA structure prediction, we found that the base pairing between the cHP2/cHP2-like structures and the sHP in a circularized vRNA is a conserved feature among the subgroup I MBr-FVs (Figure 11B, bottom in each panel), suggesting the functional importance of this interaction.

## Discussion

Although pioneering studies have indicated the presence of the cHP2 in the YFV genome (Hahn et al., 1987) and/or its participation in genome cyclization (Khromykh et al., 2001) solely by *in silico* RNA structure prediction, no further functional investigations had been reported for more than two decades. In our recent work (Li et al., 2023), it was found that the G^151^-G^152^-G^153^ trinucleotide in the cHP2’s stem region exhibited elevated SHAPE reactivity in the circularized 17D Ins1BP-minigenome compared with that in the free 5′ terminal RNA, suggesting a structural change for the hairpin during genome cyclization. This finding led us to systematically characterize the function of the cHP2 herein. By a precise destabilization of the cHP2’s secondary structure while keeping an intact complementary relationship required for YFV genome cyclization, it was clearly demonstrated that the cHP2 hairpin *per se* functions as a *cis*-acting RNA replication element (Figure 2, Figure 2-figure supplement 1, Figure 2-supplementary files 1 and 2). In combination with phylogenetic analysis and various RNA structure prediction approaches, a novel divergent feature in the 5′ terminal RNA structures between the subgroup I and subgroup II MBr-FVs was eventually discovered (Figure 11).

The direct participation of the sHP in MBFV genome cyclization has been well studied (Villordo et al., 2010). Previous studies (Davis et al., 2013; Villordo and Gamarnik, 2013) also showed that the loop sequences of the DENV and WNV sHPs are functionally important, consistent with our findings based on YFV. Interestingly, a lot of the previously reported SHAPE data of flaviviruses (whether obtained from an sfRNA, a free 3′-UTR or a viral genome, which was mostly in the linear form within infected cells) consistently exhibited a property that low SHAPE reactivity was dominant for the GACR motif and the L5-L6 positions of the sHP (Chapman et al., 2014; Dethoff et al., 2018; Li et al., 2018; Dethoff and Weeks, 2019; Zhang et al., 2019; Li et al., 2023; Huston et al., 2024), and this phenomenon showed little dependence on the modification reagents used (either NMIA, 1M7 or 2-methylnicotinic acid imidazolide (NAI)), or the source and preparative methods of the RNA samples. In agreement with our results, these facts together strongly suggested that the GACR motif is structured. Herein, through the combination of multifaceted SHAPE methodologies, an *in vitro* RNA binding assay and a replicon assay, the information about this enigmatic interlock motif was revealed for the first time. Previous studies showed that the upstream portion of the 3′-CS is sequestered by the pseudoknot interaction with the top-loop of the DB in most of the subgroup II MBr-FVs (de Borba et al., 2019; Li et al., 2023), highlighting the presence of negative control mechanisms of genome cyclization among the flaviviruses. Our findings further suggested a conserved mechanism among all MBr-FVs that the GACR portion of the 3′-CS forms tertiary interactions with the sHP to establish a tight restriction on genome cyclization. These discoveries also challenged the traditional description that the 3′-CS is unstructured in a linear genome, and implied the complexity of the dynamic flavivirus RNA structurome.

Although we analyzed the importance of the GACG and sHP Loop in a sub-viral replicon system of YFV, their functional significance in a fully-competent viral system has been indicated by previous studies. In addition to the above-mentioned works about the sHP’s loop in WNV and DENV (Davis et al., 2013; Villordo and Gamarnik, 2013), Basu et al. reported that the WNV mutants containing a single substitution in the GACA motif formed pinpoint-sized plaques in BHK-21 cells, whereas single mutations targeting other sites in the 5′-CS and 3′-CS had smaller effects on the reduction of plaque size. The GACA-targeting mutation also resulted in the largest attenuation in viral propagation as determined by plaque assay (Basu and Brinton, 2011), which was supportive of the crucial role of the GACR motif as suggested in this study.

Through a comprehensive mutagenesis and SHAPE analysis (Figure 6, Figure 6-figure supplement 2, Figure 8 and Figure 8-figure supplement 1), we proposed that the maintenance of the GACG-sHP interlock does not require the direct participation of the 3′-SL, or at least the pyrimidine-rich internal loop. In addition, disruption, restoration and re-localization of the DB-UCS-PK did not have apparent effects on the SHAPE reactivity profile of the GACG-sHP region (Li et al., 2023). These pieces of evidence suggested that the existence of the interlock motif does not rely on specific interactions with other RNA structures in the 3′-UTR. If this speculation is correct, the interlock motif could be one of the smallest folding units containing non-nested tertiary interactions to our knowledge.

Moreover, we showed that the L4/A^10769^ exhibited a strong 1M6 preference, which also required the presence of the GACG motif and a linear conformation of the viral genome (Figure 8 and Figure 8-figure supplements 2 and 3). Based on the theoretical framework of the differential SHAPE (Steen et al., 2012; Rice et al., 2014b), we hypothesized that one side of the adenine base of the L4/A^10769^ is involved in a base stacking interaction with the L5/G^10770^. The rationale is that the L5/G^10770^ is highly likely to form a direct contact with nucleotides in the GACG motif, whereas the L3/A^10768^ is both functionally variable and structurally flexible. The strong NMIA preference of the L3/A^10768^ may reflect a slow dynamics property of an adenosine under this specific chemical environment, since the L3 nucleotide in the sHP-L3G mutant showed a much lower reactivity with NMIA (Figure 6A), yet this mutation did not have an obvious effect on *in vitro* RNA binding and vRNA replication (Figures 6B and 7).

Base stacking is a fundamental interaction for the stabilization of RNA structures (Batey et al., 1999; Butcher and Pyle, 2011). Based on the corresponding SHAPE reactivity profiles (Figure 6, Figure 6-figure supplement 2, Figure 8 and Figure 8-figure supplement 1), both the L1/G^10766^ and L6/A^10771^ should be structured. Therefore, there is a considerable probability that the first and last nucleotides in the hexaloop of the sHP constitute a *trans* Sugar-Hoogsteen GA base pair (also known as the GA sheared base pair) (Leontis and Westhof, 2001; Chen et al., 2005), as *the* trans Sugar-Hoogsteen GA pair is quite stable, prevailing among the GA base pair families (https://www.nakb.org/ndbmodule/bp-catalog/), and also consistent with the strand orientation of the loop of the sHP (Leontis et al., 2002; Stombaugh et al., 2009). Moreover, the formation of this base pair would enable a continuous stacking interaction from the sHP’s stem to the L4/A^10769^, which further stabilizes the interlock motif. However, the verification of the above arguments and the complete unveiling of the structural details of the interlock motif would require more direct biophysical approaches, such as nuclear magnetic resonance spectroscopy and X-ray crystallography.

It is plausible to hypothesize that the interlock motif also functions through mechanisms other than the negative regulatory role in genome cyclization, since the deleterious effects of the sHP-M1C and sHP-M2C mutants on vRNA replication were not very likely due to unbalanced genome cyclization, although differences between *in vitro* and *in vivo* conditions could theoretically cause such discrepancies. In fact, the 3′-SL immediately downstream of the interlock motif serves as an important target for the interactions with host factors (Davis et al., 2007; Davis et al., 2013; Friedrich et al., 2014; Friedrich et al., 2018; Mazeaud et al., 2018). It was reported that host eukaryotic translation elongation factor 1A (eEF1A) interacts with the 3′-SL and sHP in several flaviviruses, but the experiments were performed with a truncated RNA which lacks the GACR motif (Davis et al., 2007; Davis et al., 2013). In addition, the reported eEF1A major binding site was located within the top stem region of the 3′-SL. Thus, its specific affinity with the 3′-UTR should be less related to the interlock motif. However, tertiary motifs are important targets for the recognition by RNA binding proteins (Hainzl et al., 2002; Oubridge et al., 2002; Huang and Lilley, 2018; Choksupmanee et al., 2021; Meier-Stephenson, 2022; Chen et al., 2023), and it would also be of interest to identify if any host factors are recruited by the GACG-sHP interlock in further work.

In summary, this study identified a 5′ terminal *cis*-acting RNA element conserved in the subgroup I MBr-FVs. Using YFV 17D as a model, we showed that the cHP2 operates in a dual-mode fashion: acting as a replicational enhancer by its local structure and regulating genome cyclization via its interaction with the even more conserved sHP-interlock motif. The presence of these elements is closely associated with flavivirus phylogeny rather than with host range, suggesting that the underlying mechanisms are crucial for the replication of the corresponding MBr-FVs.

## Materials and Methods

### Cell culture and viruses

*Mesocricetus auratus* BHK-21 cells (ATCC CCL-10) were cultured in Dulbecco’s Modified Eagle’s Medium (DMEM, Thermo Fisher Scientific) containing 2 g/L NaHCO_3_, 5% fetal bovine serum (FBS), 100 U/mL penicillin and 100 μg/mL streptomycin. Species identity of the BHK-21 cells was verified by Sanger sequencing of a 1389-bp house-keeping gene coding region. The cells are mycoplasma-free as determined by routine DNA staining with 4′,6-diamidino-2-phenylindole or Hoechst 33342. 17D and the cHP2-mutated viruses were produced by the transfection of *in vitro* transcribed vRNAs into the BHK-21 cells. Viral stocks for the plaque morphology assay were also generated by using BHK-21 cells.

### Molecular cloning

#### 17D Ins1BP-minigenomes and the TBEV minigenome

The WT 17D Ins1BP-minigenome has been reported previously (Li et al., 2023). The cHP2-M1C, the sHP-M1C, the cHP2-M1C/sHP-M1C, the cHP2-M2C, the sHP-M2C, the cHP2-M2C/sHP-M2C, and the cHP2-Δ1BP mutants were generated by overlapping PCR, and the corresponding mutated replicon plasmids and the WT 17D Ins1BP-minigenome plasmid served as the templates. The other minigenome constructs, including the sHP loop mutants, the sHP-1.4CG and sHP-1.5CG mutants, the 5′-CS-M2, the sHP-InsU and the corresponding loop/stem mutants, the cHP2-G21A and cHP2-G21A-C31U mutants, as well as the sHP-PxP, cHP2-Δ1BP-M and the sHP-PxP-M minigenomes were generated by site-directed mutagenesis based on the WT Ins1BP-minigenome template. The 5′-CS-M2/3′-CS-M2 mutant was also generated by site-directed mutagenesis using the 5′-CS-M2-containing minigenome plasmid as the template DNA.

The TBEV minigenome cDNA, which contains the 5′ first 300 nt, the last 17 nt in the NS5 coding region and the 3′-UTR sequence containing a 23-nt deletion in the long internal A-tract, was chemically synthesized based on the strain Neudoerfl (U27495.1) and cloned into pUC57 by Sangon Biotech.

#### 17D-Ins1BP-SP-IRES-Rluc-Rep replicons

The 17D-Ins1BP-SP-IRES-Rluc-Rep replicon system has been described previously (Li et al., 2023). For the cHP2-M1A, cHP2-M1C, cHP2-M2A, cHP2-M2C, cHP2-G21A, cHP2-G21A-C31U, cHP2-Δ1BP, and 5′-CS-M1 replicon plasmids, subcloned Not I-SP6-17D-Ins1BP-5′-238 nt-SnaB I fragments containing the above mutants were first generated by a cassette-cloning strategy similar to that described before (Li et al., 2023), and a specialized cassette vector, in which the 135- to 166-nt region from a parental 17D genome was replaced by the following sequence 5′-<u>GAGACG</u>AACACATACCTAATGGGA<u>CGTCTC</u>-3′ (BsmB I sites were underlined), was utilized. The sHP-M1C, the sHP-M2C, the sHP-A3U, the 3′-CS-M1, the sHP-InsU, and the sHP-InsU-L1A mutations were first introduced into a subcloned Blp I-Xho I fragment which contained the last 293 nt of the 17D genome by site-directed mutagenesis. The corresponding replicon plasmids were then generated by routine restriction endonuclease-based cloning procedures. For the replicons containing mutations in both the 5′ and the 3′ regions, two rounds of molecular cloning were performed.

For the other replicon plasmids, the mutation-containing 3′ end fragments were amplified by high-fidelity PCR using the corresponding minigenomes as the templates. Then the DNA products were subjected to restriction endonuclease digestion and subcloned into the replicon backbone as described above. To generate the 5′-CS-M2/3′-CS-M2 replicon plasmid, the 5′-CS-M2 mutation was first introduced into the Not I-Apa I fragment derived from the WT replicon plasmid through site-directed mutagenesis. Then, the mutation-containing Not I-Apa I fragment was subcloned into the pre-generated 3′-CS-M2 replicon plasmid by using *Not* I/*Apa* I double digestion.

#### 17D infectious clones

The cHP2-targeting mutations were introduced into a subcloned Not I-Nsi I fragment derived from the pACNR-FLYF17Dx infectious clone (Bredenbeek et al., 2003) by site-directed mutagenesis. Then the full-length infectious clones containing the mutations were constructed through restriction endonuclease digestion followed by ligation. The ΔGDD-containing plasmids were generated using similar approaches.

#### p4-Rluc-Rep replicons and the corresponding mutants

Since we want to characterize the function of the sHP’s loop in a replicon system closely resembling the original organization of a flavivirus genome, the p4-Rluc-Rep was generated based on the p4 infectious clone (Lai et al., 1991; Durbin et al., 2001) and the p4-DualStop-SP-IRES-Rluc-Rep replicon (Liu et al., 2016) of the DENV4 814669 strain. The resulting replicon RNA was organized as follows: the 5′-UTR, the first 147 nt of the C-coding region, the coding sequences for the Renilla luciferase reporter and foot and mouth disease virus (FMDV) 2A autocleavage peptide, the coding sequences of the last 24 residues in the envelope protein and of the nonstructural proteins, and the 3′-UTR. The p4-Rluc-Rep-GVD plasmid was generated similarly by using the p4-DualStop-SP-IRES-Rluc-Rep-GVD plasmid as the backbone. The sHP-L5U mutant was first generated by site-directed mutagenesis based on a recombinant plasmid containing a DENV4 3′-UTR, and the corresponding mutated replicon plasmid was then constructed by Mlu I/Age I-directed subcloning.

### RNA preparations

All DNA templates for *in vitro* transcription experiments were produced by high-fidelity PCR with Pyrobest DNA polymerase (Takara Biotech) or Q5 Hot Start High-Fidelity 2× Master Mix (New England Biolabs) and recovered by using a Wizard SV Gel and PCR Clean-Up System (Promega). In most cases, the PCR cleanup protocol was employed. Replicon and full-length RNA preparations were generated by using a RiboMAX SP6 Large Scale RNA Production System (Promega) and purified by using a Total RNA Kit I (Omega) as described previously (Li et al., 2023). Uncapped 5′, 3′ and minigenome RNAs were also generated following a similar protocol for the SHAPE and SHAPE-MaP experiments, while the RNAs for EMSA were first recovered by LiCl precipitation, and then further purified by urea PAGE and the “Crush & Soak” method. All RNA preparations were quantified by using a NanoDrop One UV-Vis spectrophotometer and stored at −80°C until use. For RNA preparations used in the SHAPE-MaP experiments, TURBO DNase (Thermo Fisher Scientific) was utilized to digest the DNA template instead of RQ1 RNase-Free DNase (Promega).

### Replicon assay

The 17D-Ins1BP-SP-IRES-Rluc-Rep replicon assays were performed in 48-well format as described previously (Li et al., 2023). For the p4-Rluc-Rep replicon assays, 250 ng per well of RNA was transfected in BHK-21 cells at approximately 50% confluency by using Lipofectamine 2000 (Thermo Fisher Scientific), and the cell lysates were collected at 6, 24, 48 and 72 h posttransfection. The measurement of Renilla luciferase activity was conducted as described previously (Li et al., 2023).

### vRNA quantitation by qRT-PCR

Transfection of vRNAs into BHK-21 cells and qRT-PCR were performed in triplicate as described before (Li et al., 2023). In brief, total cellular RNA was isolated by using a Total RNA Kit I (Omega) at the indicated time points and qRT-PCR was performed by using One Step PrimeScript RT-PCR Kit (Perfect Real Time) (Takara Biotech) on a QuantStudio 3 Real-Time PCR system (Applied Biosystems). The temperature for the annealing-extension step was set to 56°C, and the following primer/probe sets targeting the 17D NS5 sequence were utilized: forward: 5′-GACTTATGCTCTGAACAC-3′; reverse: 5′-TCATCACAATCTTGAACAT-3′; probe: 5′-6-FAM-CACCATCTCTGCTTCTGCCAT-TAMRA-3′. A ten-fold serial-dilution of the WT 17D-Ins1BP-SP-IRES-Rluc-Rep replicon RNA was used to set up a standard curve. The RNA stability assay using the ΔGDD-containing vRNAs was performed similarly, with the exceptions that the cells were treated by three cycles of washing with 1 mL of fresh DMEM containing 2% FBS at 6 h posttransfection and the reactions were performed on a QuantStudio 1 system (Applied Biosystems).

### Virus propagation assay

Culture supernatants of the transfected BHK-21 cells were collected at 18, 24 and 36 h posttransfection. The progeny virus in the supernatants was quantified by using plaque assay. For the observation of plaque morphology, plaque assay was performed using the Passage 1 stocks and the infected cells were cultured for 4 days before the fixation step.

### EMSA

The EMSA experiments were performed and analyzed as described (Li et al., 2023). 15-well native PAGE gels (8.3 cm × 7.3 cm × 0.75 mm) were utilized for the resolution of RNA bands. Prior to band quantification by using ImageJ 1.52a, all electrophoretic images were subjected to the same contrast adjustment, which was performed in ImageJ 1.52a, and this adjustment was also kept in the demonstrated representative images. Moreover, probably due to the inherent attenuation of the UV light source, we found an obvious difference in signal strength among the experiments which were performed within a time span of approximately 1.5 years, despite the fact that the same imaging parameters were used. Although such a variation only slightly affects the results of quantification, it could result in misunderstandings about the interpretation of the results if the images from different experiments were shown in parallel. On the condition that the representative images of all the groups belonging to the same panel were from a single experiment (which we had followed strictly in previous studies), a further linear and consistent adjustment of contrast and brightness was performed for the representative images shown in Figure 4B. The original and adjusted images were provided in the corresponding source data.

### SHAPE

Typically, 10 pmol per reaction of RNAs were denatured and refolded in 1×RNA folding buffer (100 mM HEPES, pH 8.0, 100 mM NaCl, 6 mM MgCl_2_) as described previously (Li et al., 2023). The refolded RNAs were split and modified with 13 mM of NMIA (Sigma-Aldrich) or incubated with bulk DMSO solvent at 37°C for 45 min, followed by an RNA cleanup step by using RNA Clean and Concentrator 5 (Zymo Research). Reverse transcription was then performed for the SHAPE (+) and SHAPE (−) groups along with the ddATP sequencing reaction by using Superscript III reverse transcriptase (Thermo Fisher Scientific). Primers for the 5′ reactions were 5′-FAM-TGGGCTGTGATCTTTTTTC-3′ (for the SHAPE (+)/(−) reactions) and 5′-NED-TGGGCTGTGATCTTTTTTC-3′ (for the dideoxy sequencing reactions), and primers used for the 3′ reactions were 5′-FAM-AGTGGTTTTGTGTTTGTCATCC-3′ (for the SHAPE (+)/(−) reactions) and 5′-NED/HEX-AGTGGTTTTGTGTTTGTCATCC-3′ (for the sequencing reactions). The sequencing products were split equally and added into the SHAPE (+) and SHAPE (−) reactions respectively. The mixtures were then purified by ethanol/EDTA precipitation and sent to Sangon Biotech (Shanghai) or Tsingke Biotech (Guangzhou) for capillary electrophoresis. QuShape v1.0 software (Karabiber et al., 2013) was used for the analysis of the SHAPE data following the authors’ instructions. The signal decay correction mode was set to the exponential method and negative values were set to zero automatically. For very long reads (the 3′ reactions for the Ins1BP-minigenomes), the “reactive scale by windowing” option was selected. In addition, there was a region (mostly spanning the xrRNA) with an obvious baseline drift in some of the 3′ reactions, and the baseline window parameter was set to 20 under such a situation.

### SHAPE-MaP

The refolded 17D-derived RNAs were modified with 10 mM of 1M7 (Aladdin), 1M6 (TCI) or NMIA (Sigma-Aldrich) at 37°C for 3 min, 3 min or 22 min respectively. The RNA cleanup procedure was performed as described above and the products were then subjected to reverse transcription reactions by using the Superscript II enzyme (Thermo Fisher Scientific) in the presence of Mn^2+^ as reported previously (Siegfried et al., 2014; Smola et al., 2015; Liao et al., 2026). The RT primers were as follows: 5′-ATCCTTGAACACCTCTTG-3′ for the 5′-Ins1BP-312 nt RNAs, and 5′-AGTGGTTTTGTGTTTGTCATCC-3′ for the 3′-UTR and minigenome RNAs. High-fidelity PCR was then performed by using Q5 Hot Start High-Fidelity 2× Master Mix (New England Biolabs) as described in the original protocol (Smola et al., 2015). A single primer pair (F/R: 5′-AGTAAATCCTGTGTGCTAAT-3′/5′-ATCCTTGAACACCTCTTG-3′) was used for the 5′-Ins1BP-312 nt WT RNA, and two overlapping PCR reactions were performed for the 508-nt 3′-UTR (5′-AACACCATCTAACAGG-3′/5′-CAACCTGGAGGTCGGCTGTC-3′; 5′-CTAAGCTGTGAGGCAGTG-3′/5′-AGTGGTTTTGTGTTTGTCATCC-3′). For the 830-nt Ins1BP-minigenomes, a total of four overlapping PCR reactions (5′-AGTAAATCCTGTGTGCTAAT-3′/5′-ATCCTTGAACACCTCTTG-3′; 5′-GAACATGTCTGGTCAGT-3′/5′-CCCGGTTTCAGGTTGTGG-3′; 5′-AACACCATCTAACAGG-3′/5′-CAACCTGGAGGTCGGCTGTC-3′; 5′-CTAAGCTGTGAGGCAGTG-3′/5′-AGTGGTTTTGTGTTTGTCATCC-3′) were performed. Barcodes were included in the forward primers for multiplexing. The PCR products were quantified and pooled, then sent to GENEWIZ (Nanjing) for NGS by using a paired-end 150 bp strategy on Illumina NovaSeq sequencing platforms. Data were demultiplexed by using fastq-multx v1.02.684 (Aronesty, 2013), and internal primer binding regions were trimmed by using Trimmomatic v0.39 (Bolger et al., 2014). The terminal primer binding regions were usually left untrimmed (except where part of them was used as the barcode), since they could be readily recognized during the ShapeMapper analysis by setting them to lowercase in the .fasta/.fa template file. ShapeMapper v2.2.0 (Busan and Weeks, 2018) was utilized for the data analysis and differential SHAPE reactivity was generated as reported previously (Rice et al., 2014a; Rice et al., 2014b). Two biologically independent replicates were performed for all SHAPE-MaP experiments. For the differential SHAPE experiments, the three modification reactions and two control reactions (a 3-min control for the 1M7 and 1M6 reactions, and a 22-min control for the NMIA reactions) were performed in parallel, except for one replicate of the minigenome RNA, where the 1M7, 1M6 and the 3-min control were performed simultaneously, while the NMIA modification and the 22-min control reactions were performed together at another time point.

The SHAPE-MaP experiments of the DENV4 WT and sHP-L5U-mutated 3′-UTR, and the TBEV 3′-UTR were performed similarly with the following primers. RT primers: 5′-AGAACCTGTTGGATCAAC-3′ for the DENV4 3′-UTRs and 5′-AGCGGGTGTTTTTCCGAGTCACACATCAC-3′ for the TBEV 3′-UTR. For the DENV4 3′-UTRs, two amplicons were prepared by using the following primer sets: 5′-TTACCAACAACAAACACC-3′/5′-ATGCTGTTTTTGTGTTGG-3′; 5′-AGGCGTAATAATCCCCAG-3′/5′-AGAACCTGTTGGATCAAC-3′. Three pairs of primers were used to cover the TBEV 3′-UTR: amplicon A: 5′-CTAAACCCAGACTGTGAC-3′/5′-GCTATGAAGCAGTCCCGTAG-3′; amplicon B: 5′-GAGGCTGAGCTAAAAGTTCC-3′/5′-TTTTTCAGAGTGCCCTAC-3′; amplicon C: 5′-GAGTGGCGACGGGAAAATG-3′/5′-TGTTTTTCCGAGTCACACATCAC-3′. The complete sequences of barcode-containing primers were provided in Supplementary Table S1. We noted that the actual TBEV 3′-UTR RNA for the SHAPE-MaP experiments also contained the upstream 17-nt sequence from the NS5-coding region. In addition, due to a trimming parameter setting, all of the “R1” reads of the TBEV amplicon C groups were trimmed by one additional nucleotide at the 5′ end, which did not affect any potential inter-group comparisons. The SHAPE-MaP sequencing data have been deposited in Sequence Read Archive (PRJNA1516880).

### RNA structure prediction and demonstration

#### SHAPE-guided RNA structure prediction

The structure of the WT 5′-Ins1BP-312 nt RNA was predicted by using *RNAstructure* v6.3 (Reuter and Mathews, 2010) with the average values of SHAPE reactivity for the 10^th^-255^th^ region (with the insertions in the cHP1 structure counted in) obtained by NMIA-probing-based SHAPE experiments as a constraint. The first three nucleotides in the 5′ terminus were forced to be single-stranded. The obtained RNA structure model of the WT 5′-Ins1BP-312 nt RNA served as a template for the linear 5′ end structures in Figure 1 and Figure 1-figure supplement 1. The G21A and G21A-C31U-mutated 5′-Ins1BP-312 nt RNAs were analyzed following the same procedures.

The structures of the 17D Ins1BP-minigenomes were predicted by using *RNAstructure*v6.3 (Reuter and Mathews, 2010) with the averaged SHAPE reactivity obtained from SHAPE or SHAPE-MaP experiments as constraints correspondingly. We noted that for the reactivity obtained from SHAPE experiments, the values for the 10^th^-775^th^ region were applied (the values for the 10^th^-260^th^ region were from the 5′ reactions and the values left were obtained by the 3′ reactions) for the prediction. Two rounds of prediction were performed for the minigenomes. The first round was conducted with only the SHAPE reactivity applied as pseudo-ΔG constraints, whereas in the second round the predicted results were refined with an additional constraint that the three pseudoknot-forming linear regions in the 3′-UTR were forced to be single-stranded. The results of the second round were used for RNA structure demonstration in Figure 1B, Figure 1-figure supplement 1, Figure 2-figure supplement 1, Figure 5A, Figure 6-figure supplement 3, Figure 8-figure supplement 2 and Figure 9-figure supplement 1. We noted that minor variations of the terminal base pairs of the UAR stem existed among different groups or the same RNA with different SHAPE constraints.

As stated above, the WT and the sHP-mutated 3′-UTRs were not subjected to SHAPE-constrained RNA structure prediction, mostly due to the fact that the WT sHP cannot be successfully predicted by *RNAstructure*, and if multiple constraints were applied, the prediction procedure would become less meaningful. Thus, a direct mapping strategy was mainly used for the validation of the structures of the various 3′-UTRs. For the sHP-InsU series of the 3′-UTRs, however, a basic RNA structure prediction using the averaged SHAPE reactivity (of the 10^th^-456^th^ region) as pseudo-ΔG constraints was performed to support the formation of the sHP-InsU stem-loop on the secondary structure level.

#### RNA structure prediction without SHAPE constraints and structure demonstration

For the prediction of the circularized structures formed between the terminal regions of YFV strains from different genotypes, *RNAstructure* v6.3 was utilized with no constraints applied first. Then, the G nucleotide corresponding to the G^152^ of 17D in the other genotypes was forced to be single-stranded, and the ranked 1^st^ results were used for demonstration in Figure 1-figure supplement 2.

For reference, the WT 5′-Ins1BP-312 nt RNA and those containing the cHP2-G21A, cHP2-G21A-C31U, cHP2-G21A-G22A-A24U, cHP2-C30U-C31U or cHP2-G21A-G22A-A24U-C30U-C31U mutation were also analyzed by using *mfold* v2.3, and the corresponding Ins1BP-minigenomes were analyzed with *mfold* v3.0. For the minigenomes, the G^153^ was forced to be double-stranded. In addition, for the cHP2-G21A-G22A-A24U-C30U-C31U-mutated minigenome, both the C^147^ and the G^153^ were set to be double-stranded. The cHP2-G21A-G22A-A24U, cHP2-C30U-C31U or cHP2-G21A-G22A-A24U-C30U-C31U minigenomes were also subjected to *RNAstructure* v6.3 following the procedure for the other two cHP2 mutants, except that no SHAPE constraints were available.

The minigenome sequences containing the cHP2-Δ1BP/sHP-InsU, the cHP2-Δ1BP/sHP-PxP and its derivatives were also analyzed by using *mfold* v3.0 with no additional constraints.

For the prediction of the consensus structure of the NKV and TBFV 3′-sHP, the last 900-nt sequences of the viruses with available full-length 3′-UTRs were subjected to the *LocaRNA* online server (Smith et al., 2010) and the default parameters were used. The consensus structure for the sHP in the MBr-FVs was generated by a manual alignment of the available 3′-CS-sHP sequences. The secondary structural units of the sHP were preferred to be aligned together. The consensus structures were demonstrated by using R2R v1.0.6 (Weinberg and Breaker, 2011). Default parameters were used for the 3′-sHP structures of the NKVs and the TBFVs, whereas for the MBr-FV sHP, the GSC weight was disabled, and the nucleotide identity coloring cutoffs were set to 0.999, 0.95, 0.9. The conservation of the cHP in the subgroup II MBr-FVs was investigated by an analysis of the 5′-500-nt sequences using the *LocaRNA* online server (Smith et al., 2010).

For the prediction of the 5′-cHP2-like structures in the subgroup I MBr-FVs, 5′ terminal sequences of the WESSV, SEPV, FRV, BgV, YOKV, ENTV and EPEV were analyzed by using the *mfold* v2.3 online server (Zuker, 2003). For the prediction of the cyclization patterns in these viruses, the 5′-300-nt and the 3′-UTR sequences were joined together, and subjected to both the *mfold* online server (v3.0) (Zuker, 2003) and *RNAstructure* v6.3 (Reuter and Mathews, 2010). No additional constraints were applied except for the BgV, in which a sub-optimal interaction between the cHP2 and the sHP was deduced.

In addition to the R2R v1.0.6 software (Weinberg and Breaker, 2011), RNA structure demonstration was also conducted by using *RNAcanvas* (Johnson et al., 2019) and *VARNA* v3.93 (Darty et al., 2009).

### Phylogenetic analysis

The ORF sequences of the corresponding flaviviruses (a total of 100 representative sequences) were retrieved from GenBank. Sequence alignment was performed by using MAFFT v7.490 (Katoh et al., 2009; Katoh and Standley, 2013) and phylogenetic analysis was done by IQ-TREE v2.0.7 (Lanfear et al., 2020). 1000 replicates of ultrafast bootstraps were performed and the best-fit model according to BIC was GTR+F+R8. The consensus tree was used for demonstration and a branch containing four classical insect-specific flaviviruses (cISFVs, cell fusing agent virus, Kamiti River virus, Quang Binh virus and *Culex* flavivirus) was set as the outgroup. The visualization of the tree was conducted by using FigTree v1.4.4 (https://tree.bio.ed.ac.uk/software/figtree/).

### Statistical analysis, definition of replicates and principle of data exclusion

Statistical analysis was performed in GraphPad Prism 7.0 (GraphPad Software). The detailed information was included in the corresponding figure legends and source data.

For the EMSA, SHAPE and SHAPE-MaP assays, experiments started from the re-folding of aliquots from the RNA stocks were defined as biological replicates. In some EMSA experiments, the same RNA binding reactions were loaded onto different native polyacrylamide gels, and since these experiments virtually had no independence with each other, they were treated as technical replicates. As such an approach was mainly to avoid accidental defects in the preparation, loading and running of the gels, only one of the two parallel technical replicates was included in the final analyses.

For the replicon and vRNA-transfection assays, mutually independent experiments performed at different times were defined as biological replicates. We also treated parallel transfections or infections within the same experiment as biological replicates.

Similar to other small-scale mechanistic studies, the small sample size did not support the identification of statistically aberrant values. Thus, no data were excluded on statistical grounds. However, experiments that failed technically (e.g., flawed capillary electrophoresis traces in the SHAPE experiments) were discarded. For the SHAPE-MaP results, in addition to the above-described selection of reactivity profiles based on the nucleotide range, if the reactivity value for a given nucleotide was only available in one of the two replicates, then the corresponding nucleotide was treated as “N.D.” by discarding the orphan reactivity.

## Material availability

Plasmids generated in this study are available from the corresponding author upon request, subject to approval by the original providers, institutional material transfer agreements, and applicable local and international regulations.

## Acknowledgments

This work is supported by the National Natural Science Foundation of China (No. 32070183) and the Guangdong Basic and Applied Basic Research Foundation/Natural Science Foundation of Guangdong Province (2024A1515013211).

## Author Contributions

Yi-Ge Luo*, Investigation, Conceptualization, Methodology, Validation, Data Curation, Formal Analysis, Visualization, Writing-Original Draft, Writing-Review & Editing*.

Jing-Yi Zhang*, Investigation, Validation, Conceptualization, Data Curation, Formal Analysis, Writing-Original Draft, Writing-Review & Editing*.

Dan Li*, Investigation, Conceptualization, Data Curation, Writing-Review & Editing*.

Yi-Tian Xie*, Investigation, Conceptualization, Methodology, Writing-Review & Editing*.

Hai-Tao Lu*, Investigation, Methodology, Writing-Review & Editing*.

Xu-Meng Feng*, Investigation, Writing-Review & Editing*.

Yu-Zhen Ding*, Methodology, Writing-Review & Editing*.

Hang-Yu Zhou*, Methodology, Writing-Review & Editing*.

Cheng-Feng Qin*, Resources, Supervision, Writing-Review & Editing*.

Zhong-Yu Liu*, Conceptualization (lead), Funding Acquisition, Supervision, Data Curation, Formal Analysis, Visualization, Writing-Original Draft, Writing-Review & Editing*.

## Data Availability Statement

SHAPE-MaP sequencing data were deposited in SRA (accession number PRJNA1516880). Source data files (including but not limited to raw gel images) have been provided for all figures and figure supplements as supplementary files.

---

**Figure 1-figure supplement 1** SHAPE-MaP analysis of the RNA fragments modeling the linear and circular forms of YFV.

SHAPE reactivity values obtained by SHAPE-MaP analyses were annotated on the corresponding structure models of the YFV 17D. Two biological replicates were performed and the generation and demonstration of RNA structure models and SHAPE reactivity profiles followed the same rules as Figure 1, except for the model of the linear 5′ end RNA, for which the structure in Figure 1A was directly used. In addition, we noted that the utilization of different SHAPE constraints (NMIA modification followed by SHAPE analysis in Figure 1 versus 1M7 modification followed by SHAPE-MaP in Figure 1-figure supplement 1) resulted in minor differences between the demonstrated panhandle structures formed by the 5ʹ and 3′ cyclization elements.

**Figure 1-figure supplement 2** Conservation of the cHP2 and the sHP among the seven genotypes of YFV.

The structures of the cHP2, the sHP, and the duplexes formed by the two elements were shown for the representative strains (with the GenBank accession numbers annotated) from the seven different genotypes of YFV. The structures were generated by *RNAstructure v*6.3. For the duplexed structures, the G nucleotide corresponding to G^152^ was forced to be single-stranded based on the SHAPE reactivity information obtained by chemical probing analysis of the 17D strain. The underlined sequences indicated the nucleotides that flanked the sHP and participated in the interaction with the cHP2.

**Figure 1-supplementary file 1** SHAPE reactivity values of three 17D-derived WT RNAs: the 5′-Ins1BP-312 nt RNA, the 3′-UTR and the Ins1BP-minigenome RNA. The datasets were obtained by SHAPE analyses based on NMIA modification and provided as zipped Excel files. The files related to the prediction by *RNAstructure* were also included.

**Figure 1-supplementary file 2** SHAPE reactivity values of three 17D-derived WT RNAs: the 5′-Ins1BP-312 nt RNA, the 3′-UTR and the Ins1BP-minigenome RNA. The datasets were obtained by SHAPE-MaP analyses based on 1M7 modification and provided as zipped Excel files. The files related to the prediction by *RNAstructure* were also included.

**Figure 1-source data** Source data for Figure 1 and Figure 1-figure supplements 1 and 2.

**Figure 2-figure supplement 1** SHAPE analysis of the G21A and G21A-C31U mutants.

The 5′-Ins1BP-312 nt RNAs and Ins1BP-minigenome RNAs containing the G21A and the G21A-C31U mutations were subjected to SHAPE analysis followed by SHAPE-guided RNA structure prediction. Detailed information about experimental setup and SHAPE reactivity annotation can be referred to in Figure 1. The most thermodynamically stable structures for the cHP2 regions and the panhandle structures formed by the 5′ and 3′ cyclization elements were illustrated by using *RNAcanvas*. The WT structures were shown in parallel with the same set of SHAPE data in Figure 1 annotated.

**Figure 2-figure supplement 2** The effects of cHP2 mutations on the intracellular vRNA stability.

The corresponding ΔGDD-containing vRNAs were transfected into BHK-21 cells in triplicate and intracellular vRNA levels were determined by qRT-PCR. The data were shown as the mean ± SD, and two-way ANOVA and Tukey’s multiple comparisons test were performed. \*\**P*<0.01, ns: not statistically significant.

**Figure 2-supplementary file 1** SHAPE reactivity values of the WT, the cHP2-G21A and the cHP2-G21A-C31U-mutated 5′-Ins1BP-312 nt RNAs. The datasets were obtained by SHAPE analyses based on NMIA modification and provided as a zipped Excel file archive. The related *RNAstructure* prediction files are also included.

**Figure 2-supplementary file 2** SHAPE reactivity values of the WT, the cHP2-G21A and the cHP2-G21A-C31U-mutated Ins1BP-minigenome RNAs. The datasets were obtained by SHAPE analyses based on NMIA modification and provided as a zipped Excel file archive. The related *RNAstructure* prediction files are also included. We noted that the dataset of the WT group is identical to that in Figure 1-supplementary file 1.

**Figure 2-supplementary file 3** RNA structure prediction results of the cHP2-mutated 5′-Ins1BP-312 nt RNAs and Ins1BP-minigenomes by *mfold*.

**Figure 2-source data** Source data for Figure 2 and Figure 2-figure supplements 1 and 2.

**Figure 3-source data** Source data for Figure 3.

**Figure 4-source data** Source data for Figure 4.

**Figure 5-supplementary file 1** SHAPE reactivity values of the WT, the cHP2-M2C, the sHP-M2C and the cHP2-M2C/sHP-M2C-mutated Ins1BP-minigenome RNAs. The datasets were obtained by SHAPE analyses based on NMIA modification and provided as a zipped Excel file archive. The related *RNAstructure* prediction files are also included. We noted that the dataset of the WT group is identical to that in Figure 1-supplementary file 1.

**Figure 5-supplementary file 2** The ranked 1^st^ predicted structures for the WT and the M2C-series of minigenome mutants by *RNAstructure* were demonstrated by using *RNAcanvas*. The zipped file includes *.rnacanvas* files and converted vector image files.

**Figure 5-supplementary file 3** SHAPE reactivity values of the sHP-M1C and the sHP-M2C-mutated 3′-UTR RNAs. The datasets were obtained by SHAPE analyses based on NMIA modification and provided as an Excel file.

**Figure 5-supplementary file 4** The SHAPE reactivity values were mapped onto the structural models of the mutated 17D 3′-UTRs by using *RNAcanvas*. The zipped file contains *.rnacanvas* files and converted vector image files.

**Figure 5-source data** Source data for Figure 5.

**Figure 6-figure supplement 1** The conservation of the 3′-CS-sHP regions in various MBr-FVs.

The shown alignment was generated manually following a structure-first principle. The coloring of the nucleotides was based on the identity and a highlighted background indicated absolute conservation. The order of the sequences was based on the similarity in the sHP structure instead of the phylogenetic distance between the viruses. For the 3′-CS sequences of Entebbe bat virus and Yokose virus, only the portions corresponding to the YFV 3′-CS were shown.

**Figure 6-figure supplement 2** SHAPE analysis of the sHP mutants generated on the basis of the sHP-InsU variant.

The data were obtained and demonstrated following the same rules as in Figure 6A. The SHAPE reactivity values of the L3/A^10768^ usually exceeded the range of the Y axis greatly, and the exact numeric values were provided in Figure 6-supplementary file 1 and the corresponding source data.

**Figure 6-figure supplement 3** SHAPE-MaP analysis of the sHP-L3G and the sHP-L5U-mutated Ins1BP-minigenomes.

The average values of the SHAPE reactivity obtained from two biologically independent SHAPE-MaP experiments were used as constraints for RNA structure prediction. The ranked 1^st^ structures from the results predicted by *RNAstructure* v6.3 were used for the demonstration. Only the panhandle structures formed by the 5′ and 3′ cyclization elements were shown along with the pseudo-ΔG for the whole minigenome structures. The corresponding region of the WT Ins1BP-minigenome was shown in parallel and annotated with the same data in Figure 1-figure supplement 1.

**Figure 6-supplementary file 1** SHAPE reactivity values of the WT and various mutated 17D 3′-UTR RNAs in Figure 6A and Figure 6-figure supplement 2. The datasets were obtained by SHAPE analyses based on NMIA modification and provided as an Excel file. We noted that the dataset of the WT 3′-UTR is identical to that in Figure 1-supplementary file 1.

**Figure 6-supplementary file 2** The SHAPE reactivity values for the groups shown in Figure 6A were mapped onto the structural models of the 17D 3′-UTR by using *RNAcanvas*. The zipped file contains *.rnacanvas* files.

**Figure 6-supplementary file 3** Structural mapping and structure prediction results for the sHP-InsU variants. The SHAPE reactivity values for the sHP-InsU variants were mapped onto the structural models of the 17D 3′-UTR by using *RNAcanvas*. The results of SHAPE-constrained RNA structure prediction by *RNAstructure* were also provided together.

**Figure 6-supplementary file 4** SHAPE reactivity values of the WT, the sHP-L3G and the sHP-L5U-mutated Ins1BP minigenome RNAs. The datasets were obtained by SHAPE-MaP analyses based on 1M7 modification and provided as zipped Excel files. The files related to the prediction by *RNAstructure* were also included. The dataset for the WT group is the same as that provided in Figure 1-supplementary file 2.

**Figure 6-source data** Source data for Figure 6 and Figure 6-figure supplements 1-3.

**Figure 7-figure supplement 1** Characterization of YFV replicons carrying modified cHP2 and/or sHP structures.

(A) Design of the cHP2-Δ1BP, the sHP-InsU and the cHP2-Δ1BP/sHP-InsU-carrying replicon variants. (B) Replicational profile of the variants. WT 17D-Ins1BP-SP-IRES-Rluc-Rep and the corresponding ΔGDD mutant were transfected in parallel. Data were expressed as described in Figures 3B, 7B and 7D. Two-way ANOVA and Tukey’s multiple comparisons test were performed. \*\*\*\**P*<0.0001, ns: not statistically significant.

**Figure 7-source data** Source data for Figure 7 and Figure 7-figure supplement 1.

**Figure 8-figure supplement 1** SHAPE-MaP analysis of the WT 3′-UTR and representative mutants.

The annotations were based on the average values of two biological replicates. Only the 3′-CS to 3′-SL regions were demonstrated. The complete datasets were included in Figure 8-supplementary file 1.

**Figure 8-figure supplement 2** Summary of the strong differential SHAPE reactivity in WT 5′, 3′ and minigenome RNAs.

The nucleotides that exhibited a strong differential SHAPE signal either in the 5′-Ins1BP-312 nt and 3′-UTR RNAs (A) or in the Ins1BP-minigenome (B) were indicated in the structural models of the 5′ and 3′ termini of the YFV genome. For these sites, the corresponding differential SHAPE reactivity values in both the free 5′/3′ RNAs and the minigenome were annotated for comparison. The colors for the differential SHAPE values followed the rules in Figure 8. A pair of black arrowheads indicated the location of the artificially inserted base pair into the cHP1 hairpin. The SHAPE reactivity annotation for the structures was based on 1M7 modification and SHAPE-MaP analysis, and the same data in Figure 1-figure supplement 1 were utilized. The numbering was based on the locations in the unmodified parental 17D genome. We noted that the variation of the differential SHAPE values of the C^87^ in the 5′-Ins1BP RNA was too large for a clear interpretation. Please refer to Figure 8-figure supplement 3 and Figure 8-supplementary file 3 for more information.

**Figure 8-figure supplement 3** Differential SHAPE profiles of the WT 5′, 3′ and minigenome RNAs.

(A) Differential SHAPE reactivity profiles for the 5′ 25-215 nt regions. The inserted base pair into the cHP1 structure was counted in the numbering, which was different from that in Figure 8-figure supplement 2. Upper panel: the differential SHAPE profile of the 5′-Ins1BP RNA. Lower panel: the differential SHAPE profile of the 5′ region in the Ins1BP-minigenome. (B) Differential SHAPE reactivity profiles for the 10650-10840 nt regions. Upper panel: the differential SHAPE profile of the corresponding region in the 3′-UTR RNA (the same data from Figure 8B were shown). Lower panel: the differential SHAPE profile of the corresponding region in the Ins1BP-minigenome. Results from two biological replicates were shown as the mean ± SD. The coloring was the same as in Figure 8B. The corresponding regions of the cHP2, the DB and the sHP were labeled and the signals for the L3/A^10768^ and L4/A^10769^ in the free 3′-UTR were indicated. The complete datasets were included in Figure 8-supplementary file 3.

**Figure 8-supplementary file 1** SHAPE reactivity values of the WT and various mutated 17D 3′-UTR RNAs in Figure 8 and Figure 8-figure supplement 1. The datasets were provided as an Excel file. We noted that the 1M7 dataset of the WT group is identical to that in Figure 1-supplementary file 2.

**Figure 8-supplementary file 2** Differential SHAPE values for the WT and 3′-CS-M2-mutated 3′-UTRs. The datasets were provided as an Excel file.

**Figure 8-supplementary file 3** SHAPE reactivity and differential SHAPE values of the WT 5′-Ins1BP-312 nt RNA and the WT Ins1BP-minigenome RNA. The datasets were obtained by SHAPE-MaP analyses based on 1M7, 1M6 and NMIA modifications and were provided as a zipped Excel file archive. Differential SHAPE values for the WT 3′-UTR were also included for reference.

**Figure 8-source data** Source data for Figure 8 and Figure 8-figure supplements 1-3.

**Figure 9-figure supplement 1** SHAPE analysis of the 5′-CS-M2/3′-CS-M2-mutated Ins1BP-minigenome.

The averaged SHAPE reactivity from two biological replicates was annotated on the predicted structure of the 5′-CS-M2/3′-CS-M2 minigenome.

**Figure 9-source data** Source data for Figure 9 and Figure 9-figure supplement 1.

**Figure 10-source data** Source data for Figure 10.

**Figure 11-figure supplement 1** Phylogenetic tree of the flaviviruses.

The bootstrap values were listed along with the nodes. The coloring of the lineages was the same as in Figure 11A. The positions of the viruses analyzed in Figure 11B were labeled with arrows. The sHP structures and the upstream sequences from two “tick-specific” flaviviruses with available 3′-UTR sequences were presented. The 3′-UTR structure of Mpulungu flavivirus, which was provided in the corresponding source data, was generated with reference to the reported 3′-UTR structures of the “tick-specific” flaviviruses. The structure of the Xinyang flavivirus sHP was drawn according to a previous study (Wang et al., 2024).

**Figure 11-figure supplement 2** SHAPE-MaP analysis of the WT DENV4 3′-UTR and the sHP-L5U mutant.

The averaged SHAPE reactivity of two biological replicates was annotated on the structure models of the WT DENV4 3′-UTR and the sHP-L5U mutant. The level of SHAPE reactivity was labeled in different colors as in Figure 1, Figure 5 and other relevant figures. Black: <0.4, Golden yellow: ≥0.4 and <0.85, Red: ≥0.85 and <2.55, Purple: ≥2.55. Gray: N.D. The numbering was based on the genome sequence of the DENV4 814669 strain. Various RNA structures were labeled and the dark violet lines represented pseudoknots.

**Figure 11-figure supplement 3** The sHP-L5G is also important for DENV4 RNA replication.

Organization of the DENV4-Rluc-Rep. The structural protein coding region was replaced with the coding sequences of the Renilla luciferase reporter and the FMDV 2A autocleavage peptide, except for the first 147 nt of the C-coding region and the last 72 nt of the E-coding region. (B) Representative results of the replicon assay. The assay was performed in triplicate and data were expressed as the mean ± SD. Two-way ANOVA and Tukey’s multiple comparisons test were performed. \*\*\*\**P*<0.0001. (C) Summary of two biologically independent experiments. One-way ANOVA and the two-stage linear step-up procedure of Benjamini, Krieger and Yekutieli were performed. \*\*\*\**P*<0.0001. The NS5-GVD control contains a GDD-to-GVD mutation in the catalytic motif of viral NS5. The error bars in C stand for SEM and the individual data points are also plotted.

**Figure 11-figure supplement 4** SHAPE-MaP analysis of the TBEV 3′-UTR.

SHAPE reactivity of the TBEV sHP region. The results based on the modification by different SHAPE reagents were shown. Two biological replicates were performed and the data were expressed as the mean ± SD. (B) The averaged SHAPE reactivity of the 1M7-modification experiments was annotated onto the structural model of the TBEV 3′-UTR, and the nucleotides with a high differential SHAPE value were annotated. The numbering was based on the synthetic TBEV 3′-UTR sequence, and the corresponding positions in the genome of the TBEV strain Neudoerfl (U27495.1) were also annotated. We noted that the long adenosine tract was shortened by 23 nt due to the technical limit of chemical synthesis, and the SHAPE resolution of this region was poor, likely due to errors in the reverse transcription and biases in the mutational counting procedure.

**Figure 11-figure supplement 5** Differential SHAPE profiles of the TBEV 3′-UTR.

Data were expressed as in Figure 8B and Figure 8-figure supplement 3. The numbering was based on the synthetic TBEV 3′-UTR sequence.

**Figure 11-supplementary file 1** The output files of a *LocARNA* analysis using the 5′-500 nt sequences from 30 representative subgroup II MBr-FVs.

**Figure 11-source data** Source data for Figure 11 and Figure 11-figure supplements 1 to 5.

