## Supplementary figures and images for "Novel conserved RNA structural motifs regulate genome cyclization of yellow fever virus"

### 5-Ins1BP-21-22-24-30-31.pdf

Output of sir_graph (©)
mfold_util 4.7

Created Sat May 9 17:11:47 2026

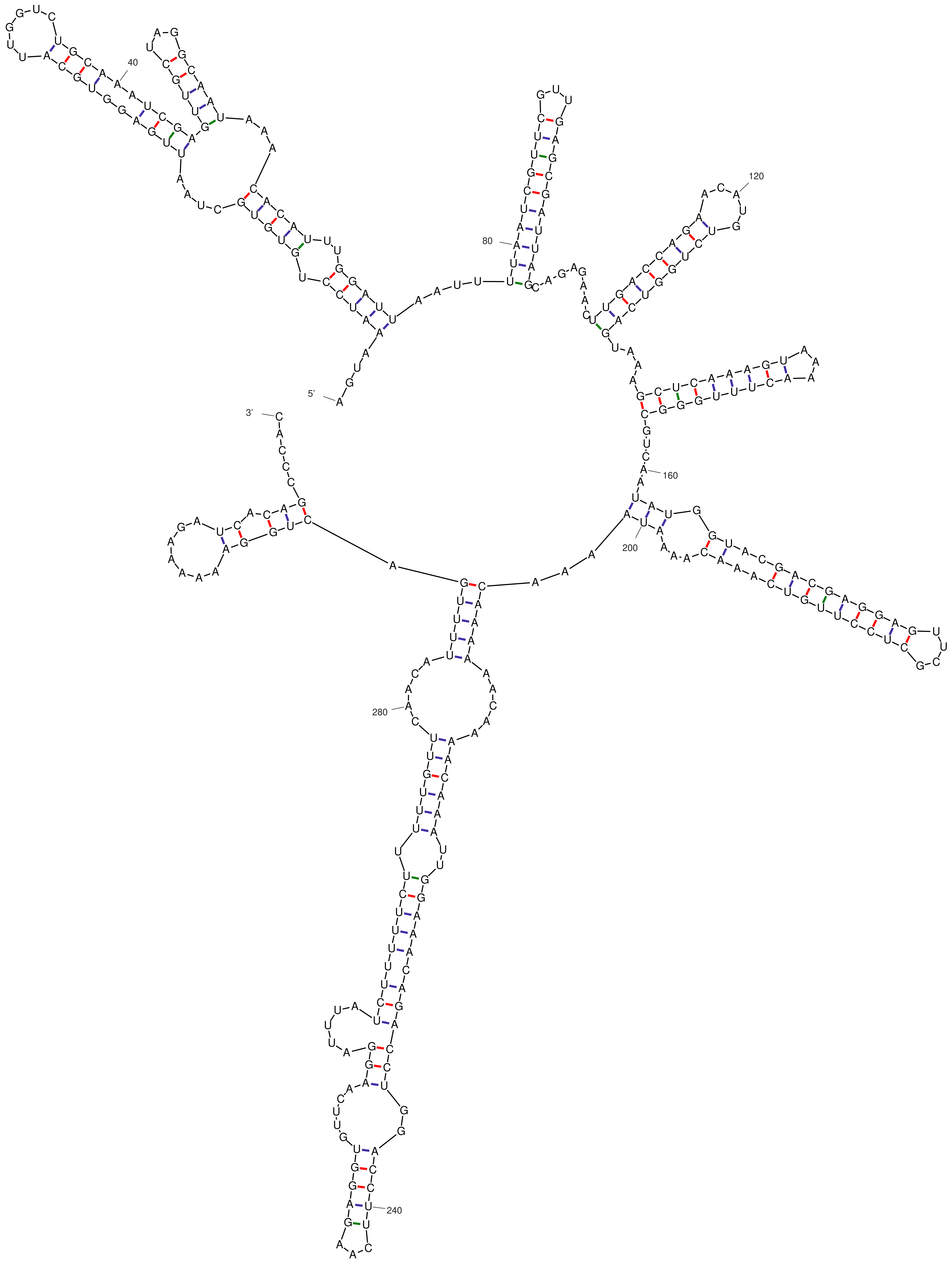


*dG = -85.80 26May09-17-11-46*

### 5-Ins1BP-21-22-24.pdf

Output of sir_graph (©)
mfold_util 4.7

Created Sat May 9 17:08:49 2026

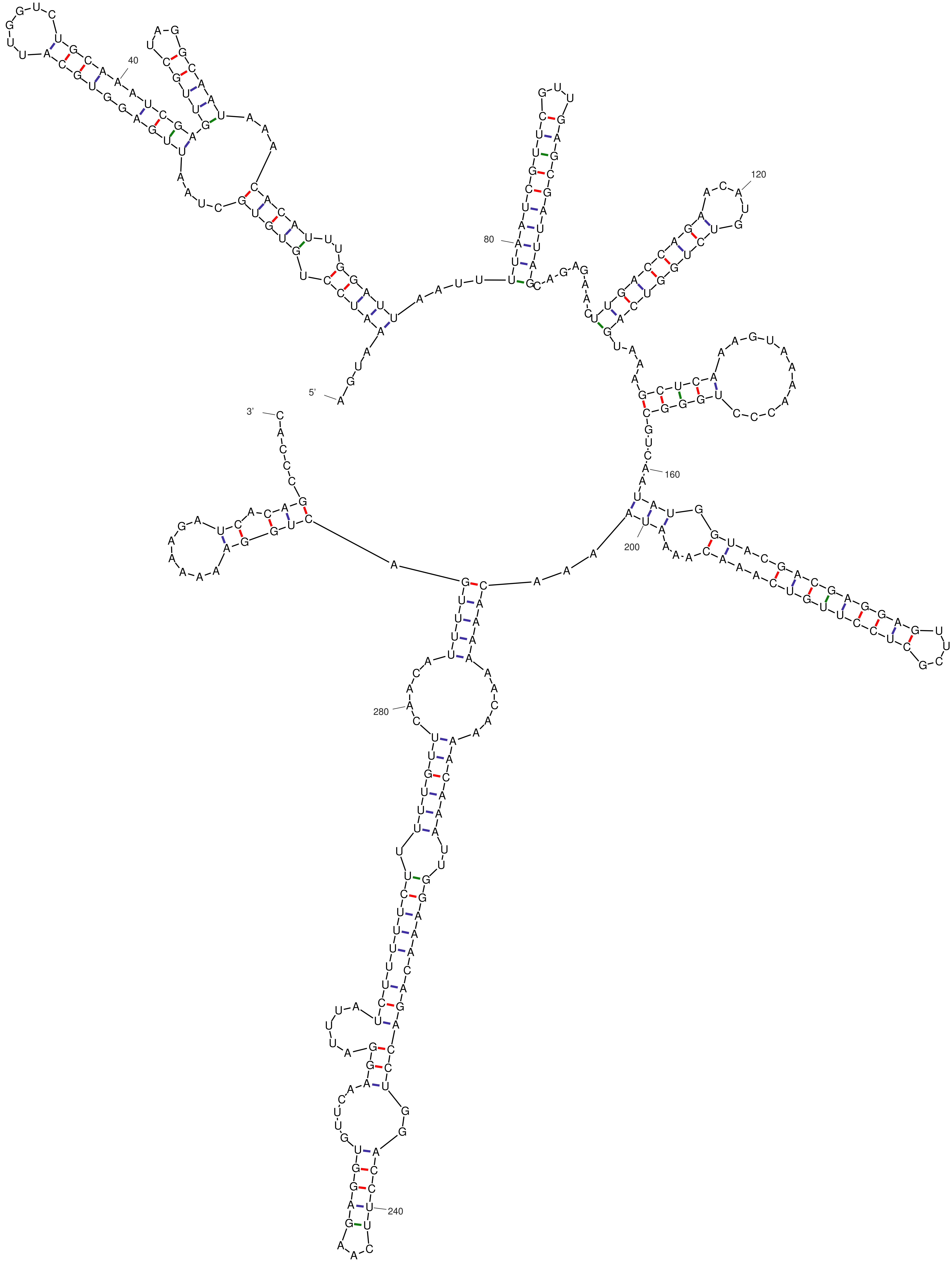


dG = -79.90 26May09-17-08-48

### 5-Ins1BP-30-31.pdf

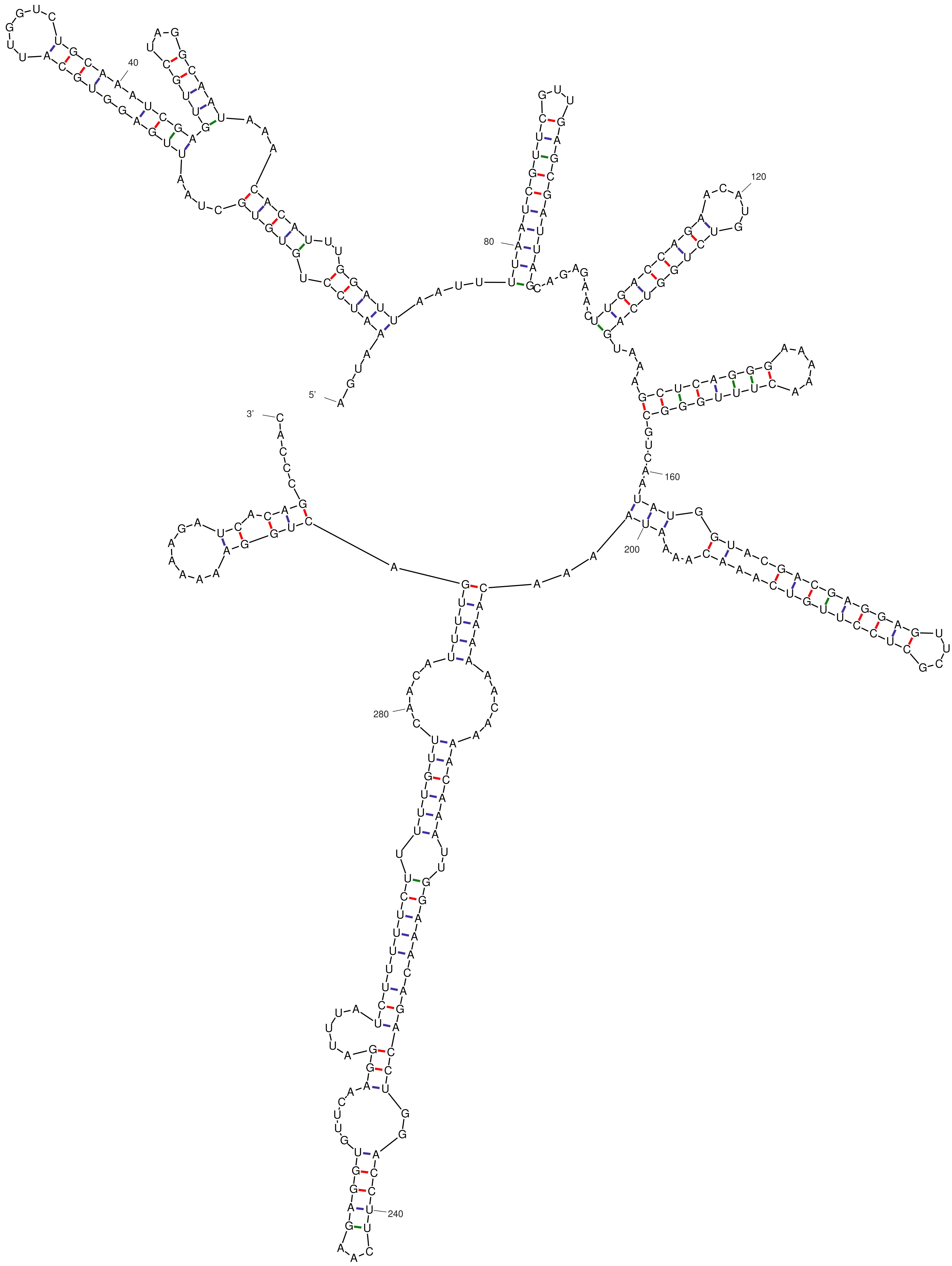

*dG = -83.80 26May09-17-10-34*

### 5-Ins1BP-G21A-C31U.pdf

Output of sir_graph (©)
mfold_util 4.7

Created Sat May  9 17:07:36 2026

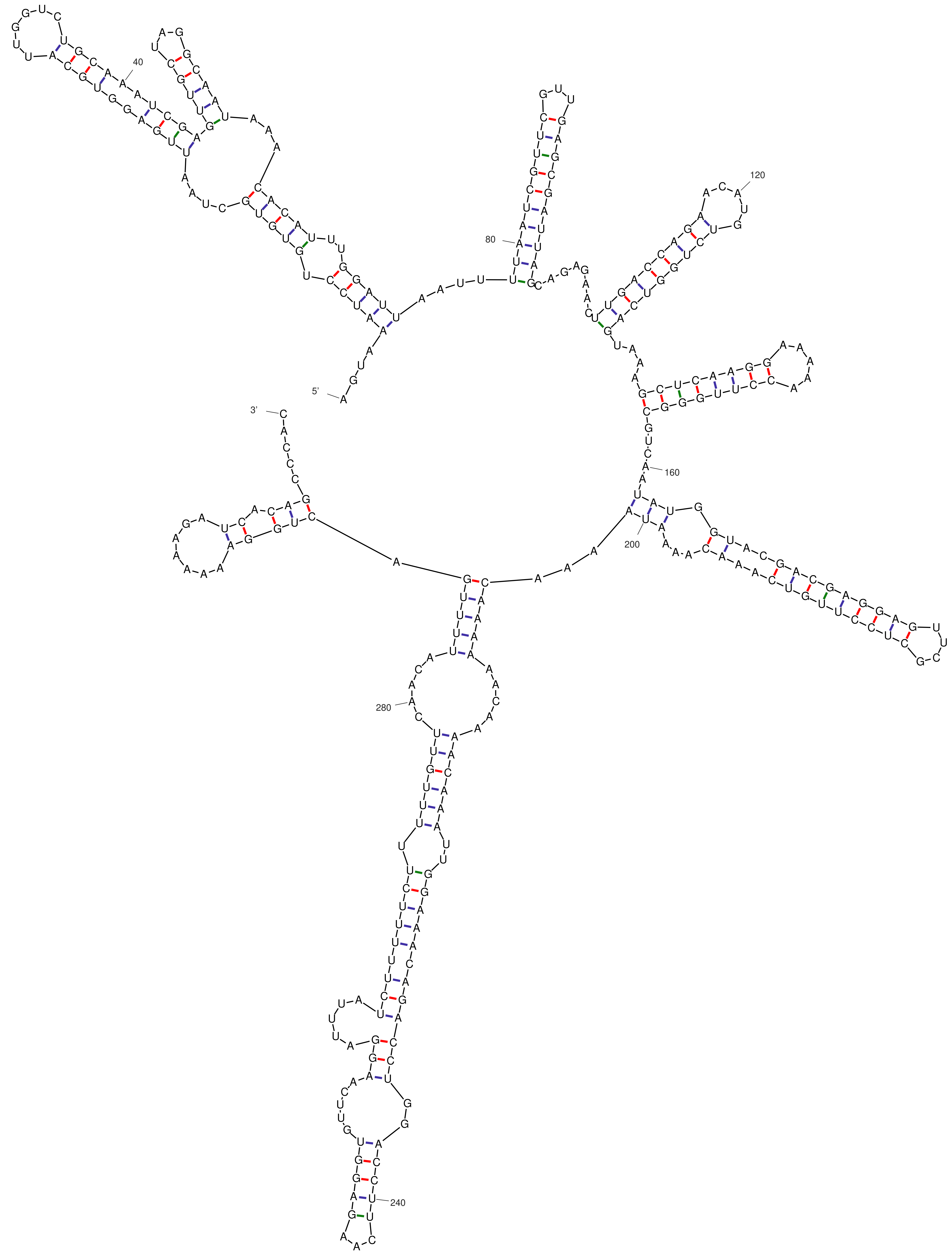


*dG = -86.50 26May09-17-07-35*

### 5-Ins1BP-G21A.pdf

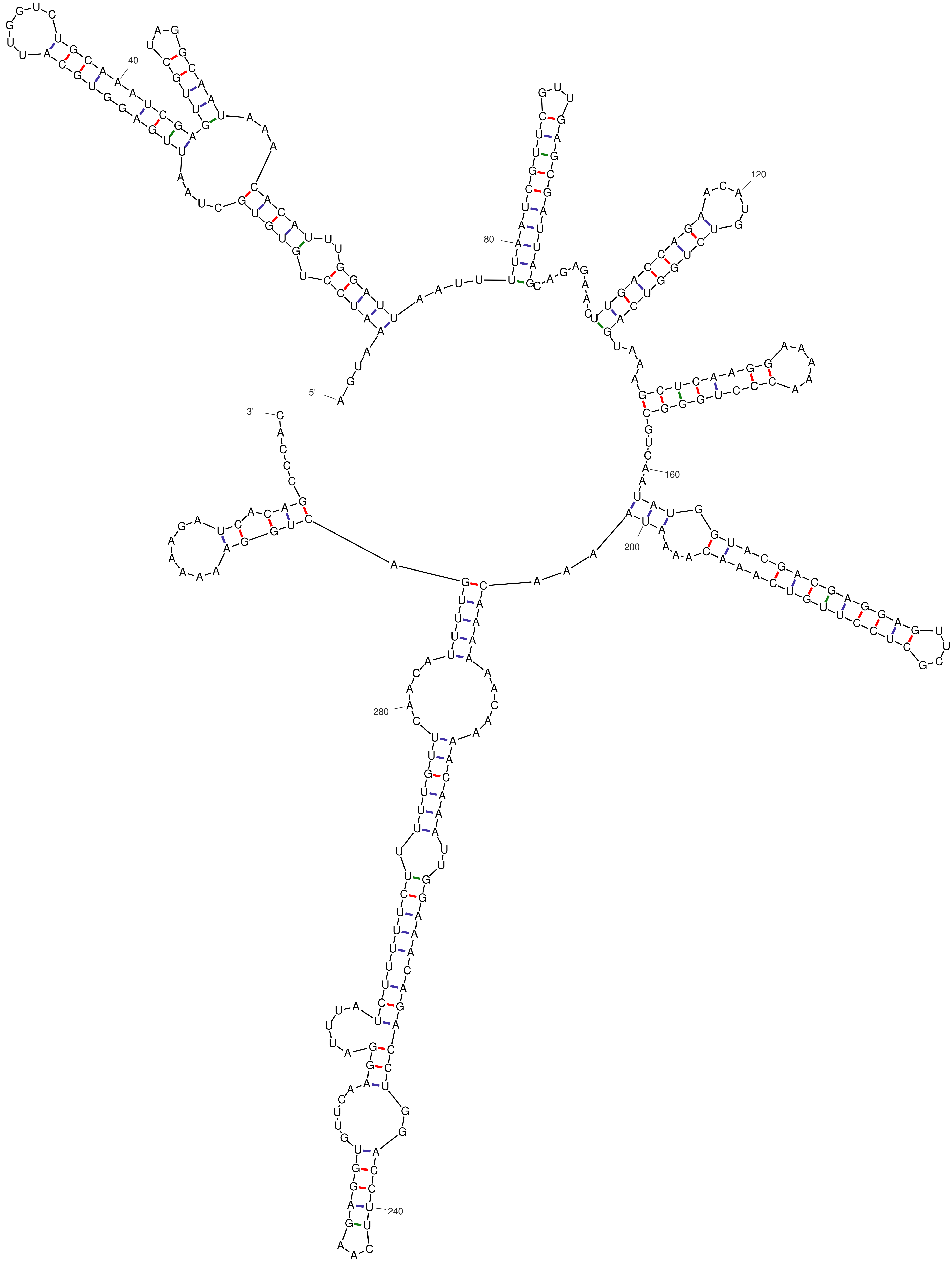


*dG = -83.10 26May09-17-06-28*

### 5-Ins1BP-WT.pdf

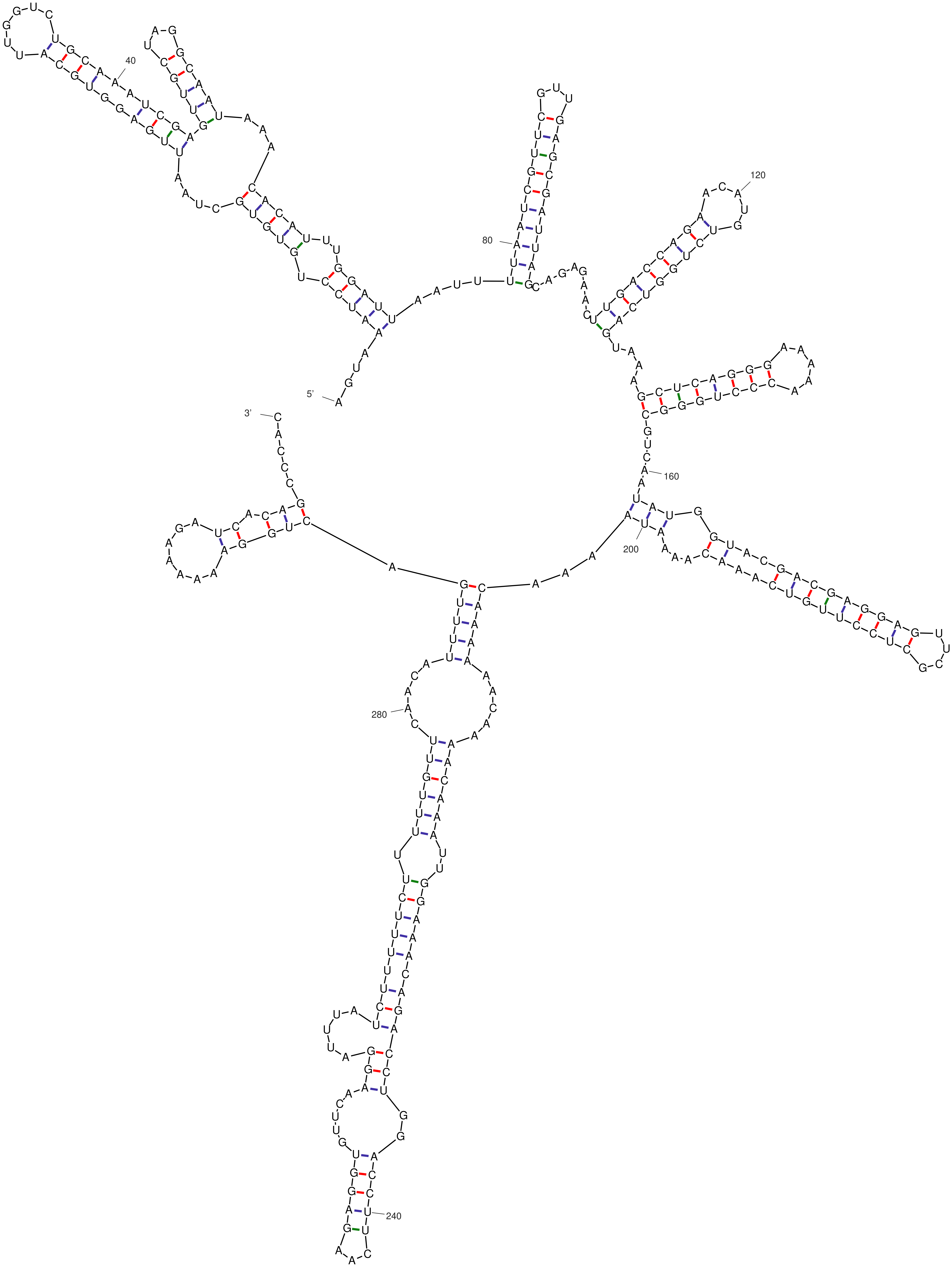


*dG = -88.50 26May09-17-05-02*

### alirna.pdf

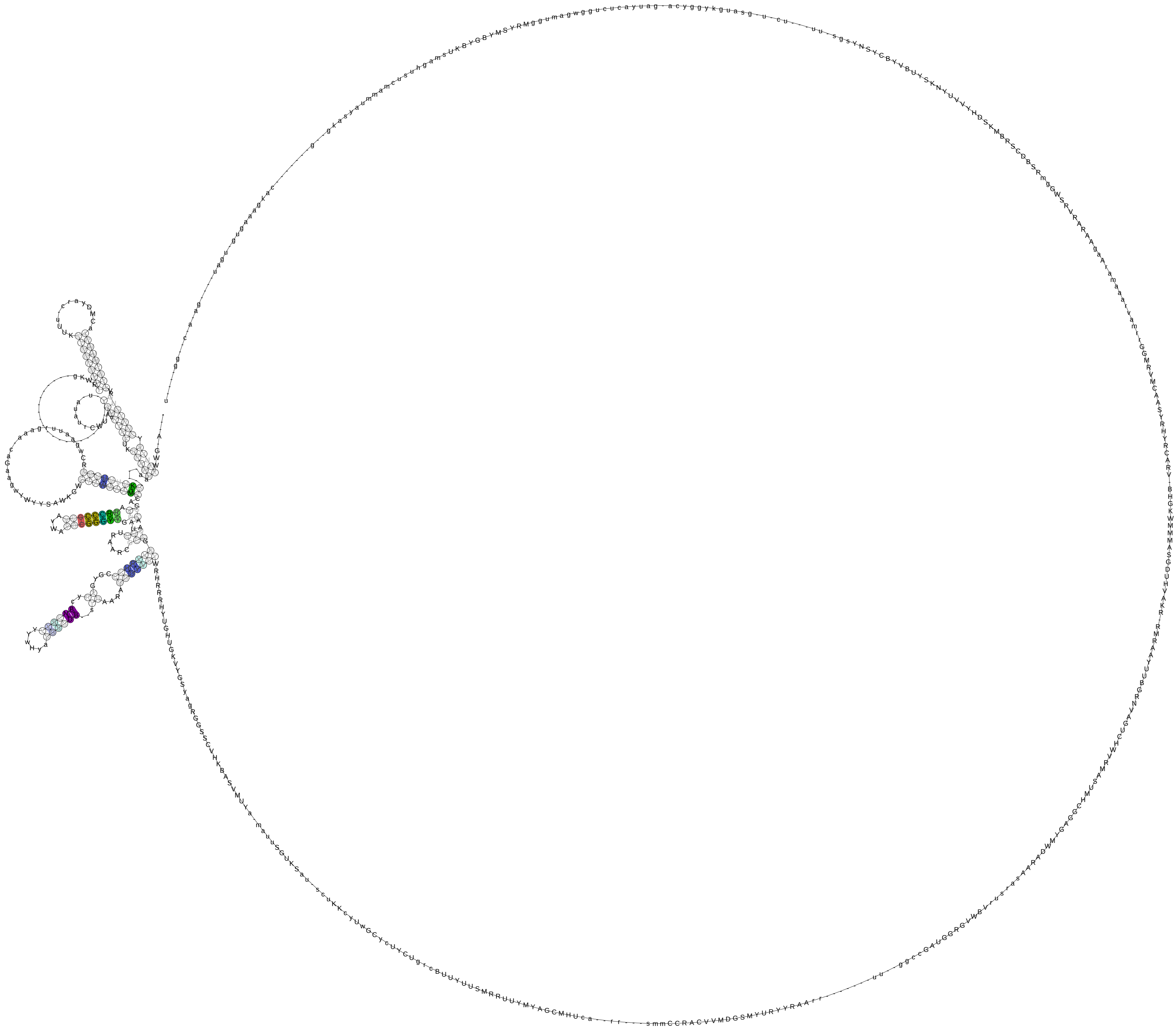

### alirna.png

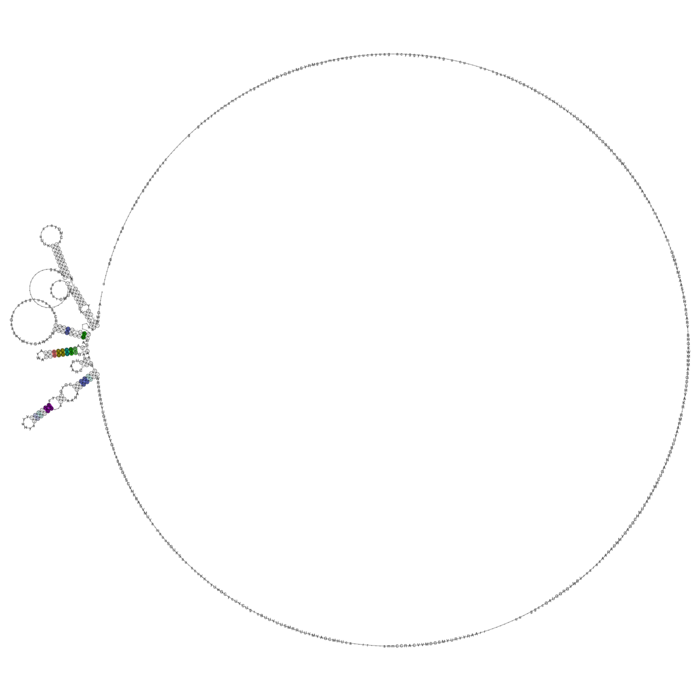

### aln.pdf

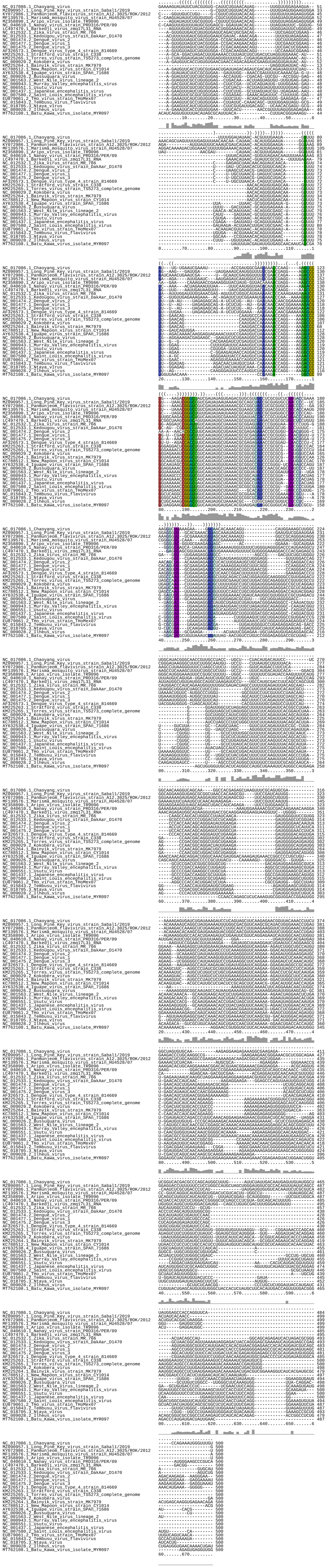

### aln.png

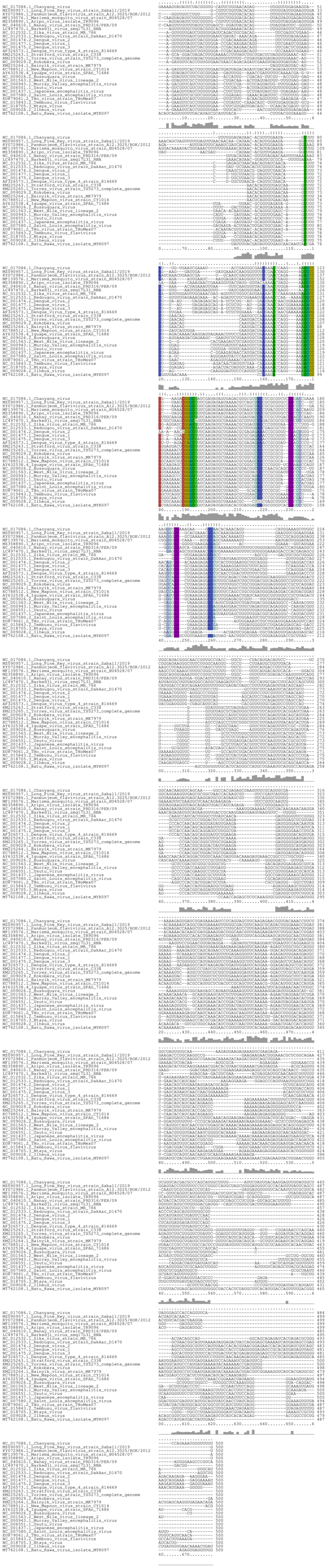

### Figure 1-figure supplement 1

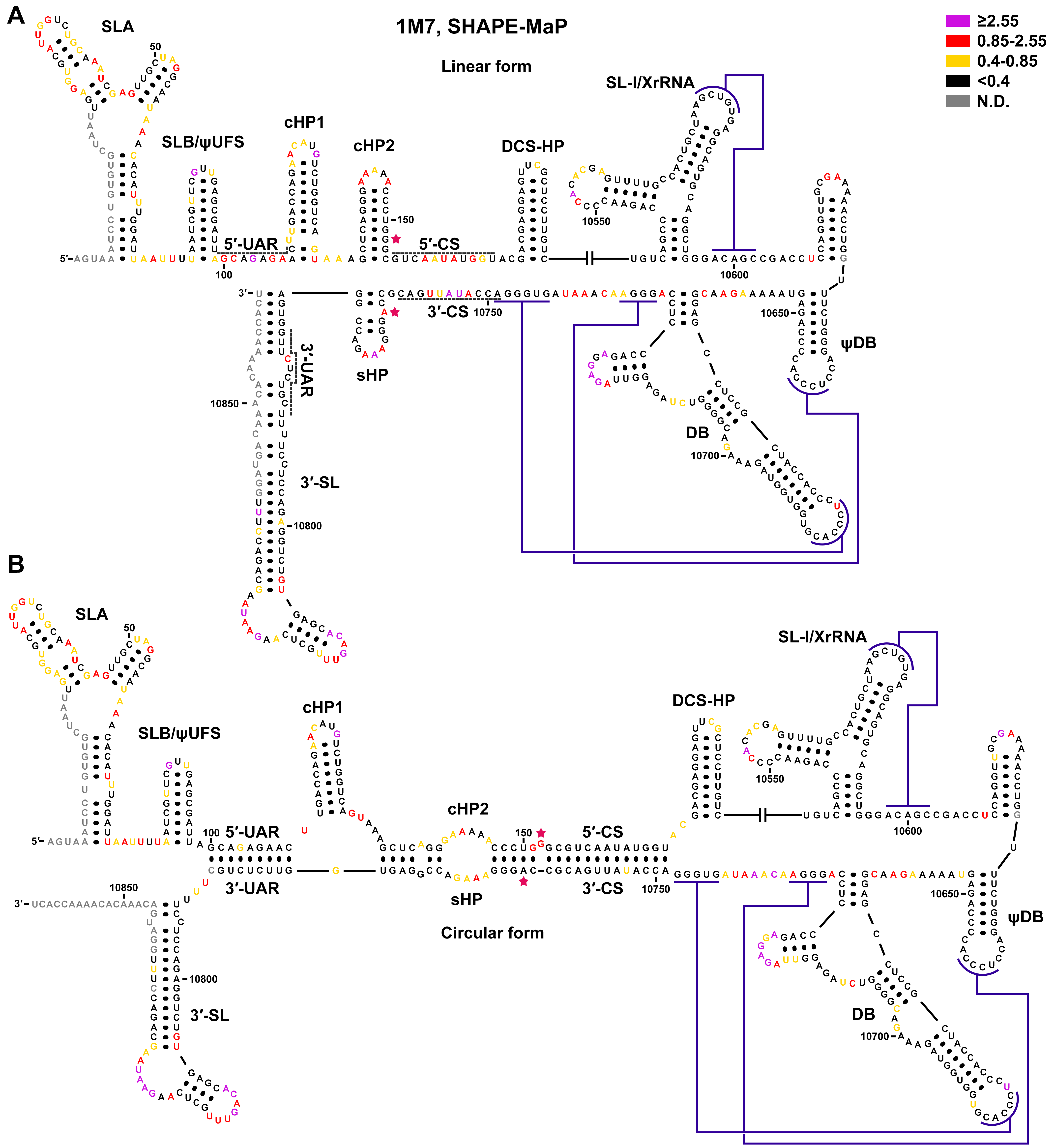

### Figure 1-figure supplement 2

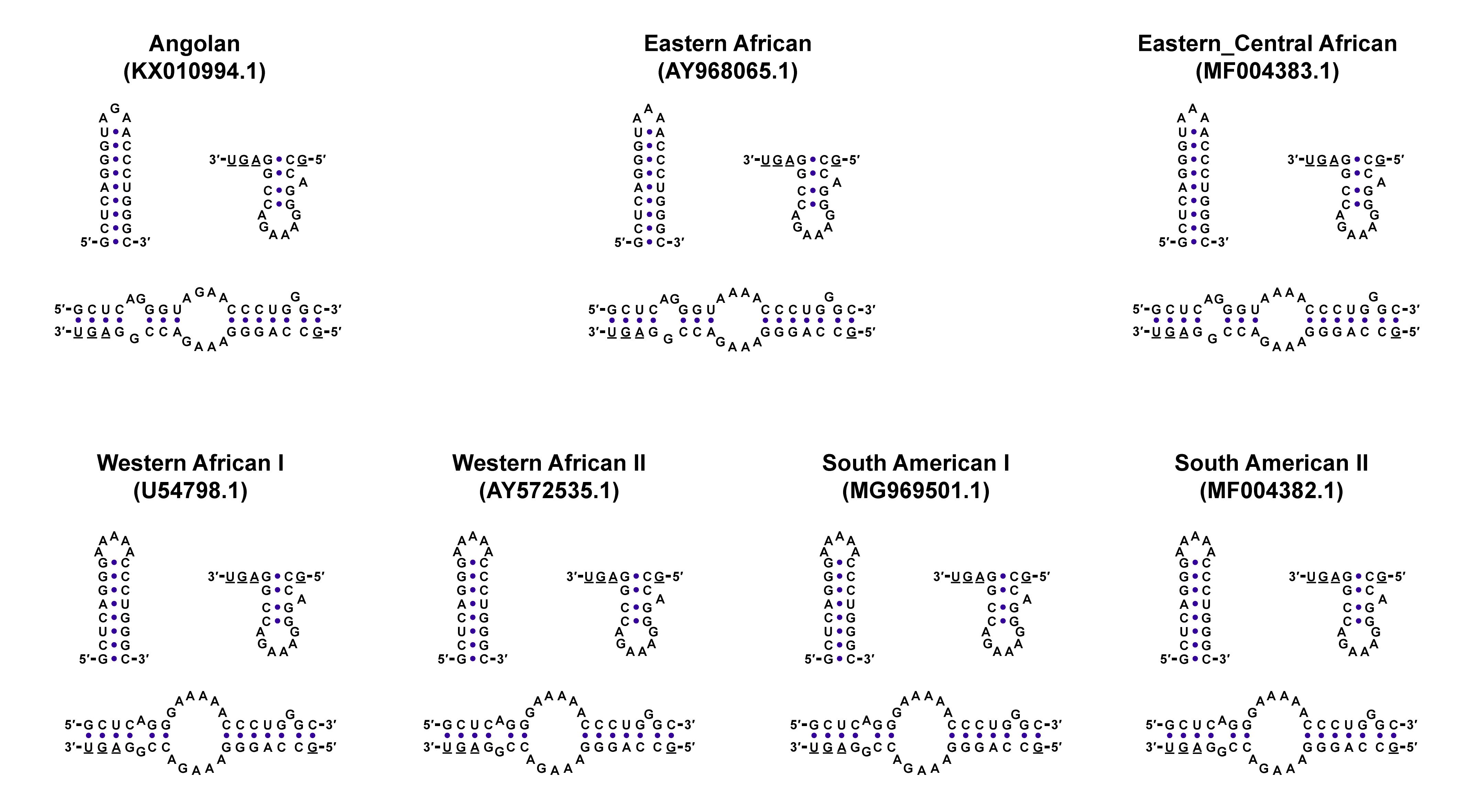

### Figure 2-figure supplement 1

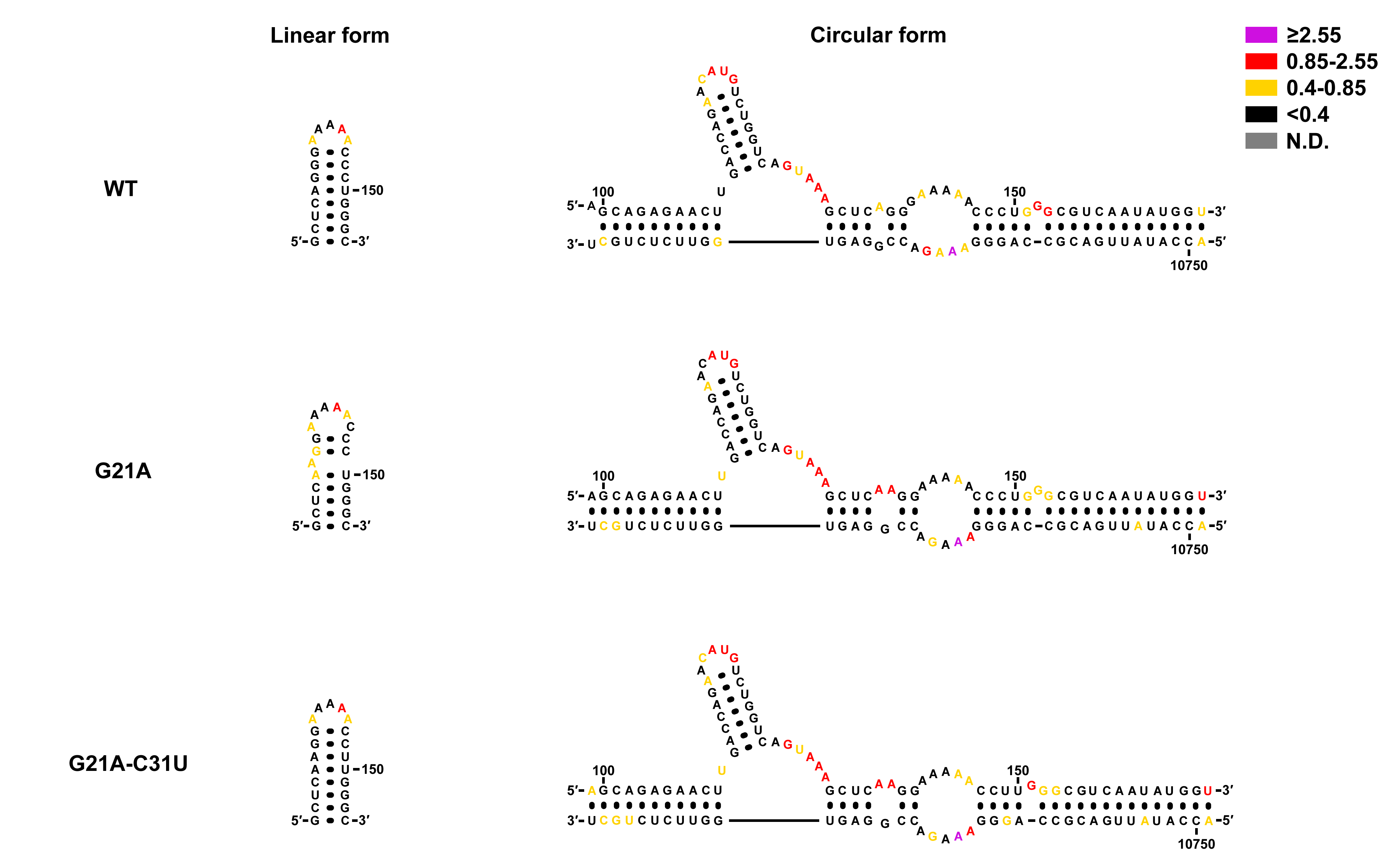

### Figure 2-figure supplement 2

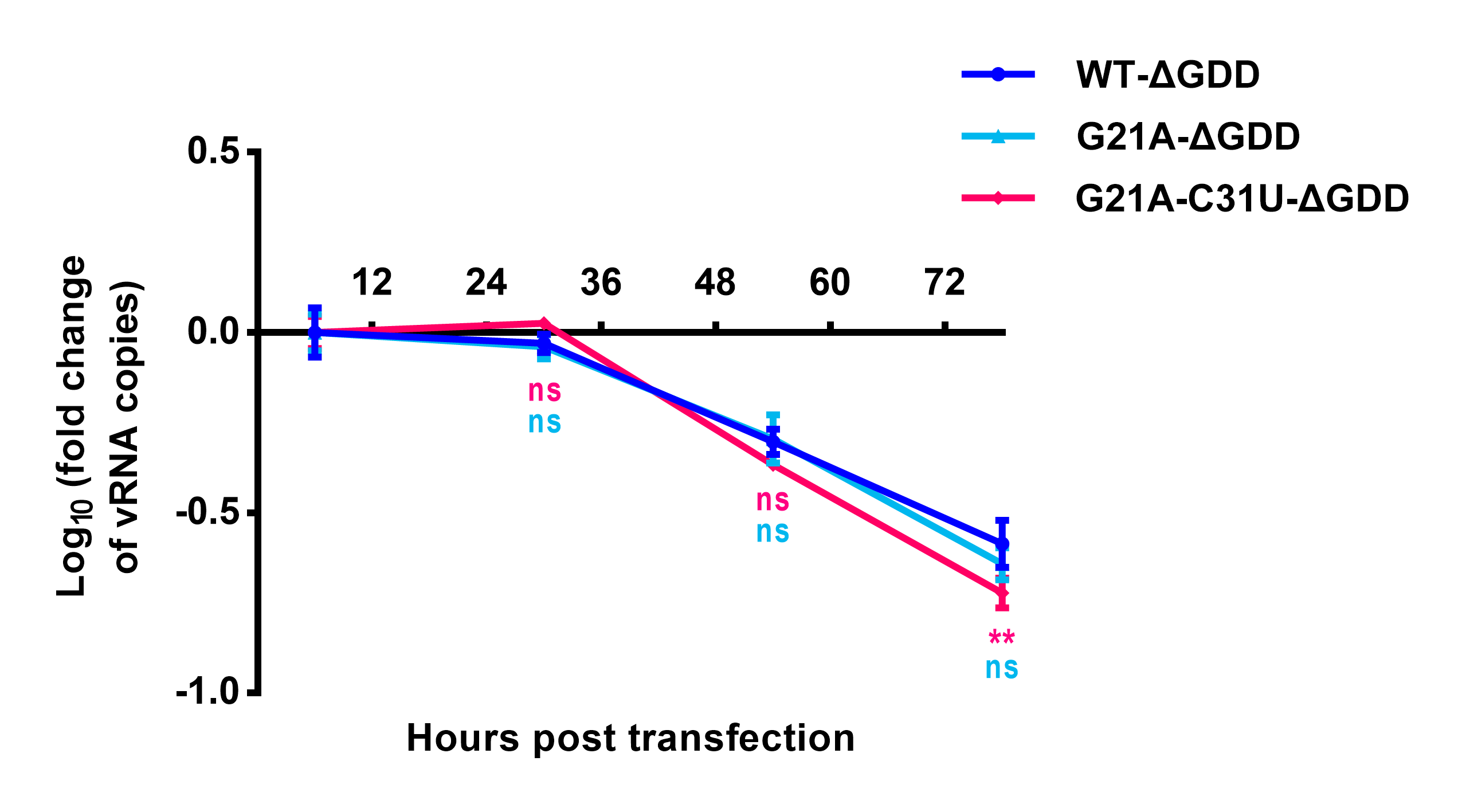

### Figure 6-figure supplement 2

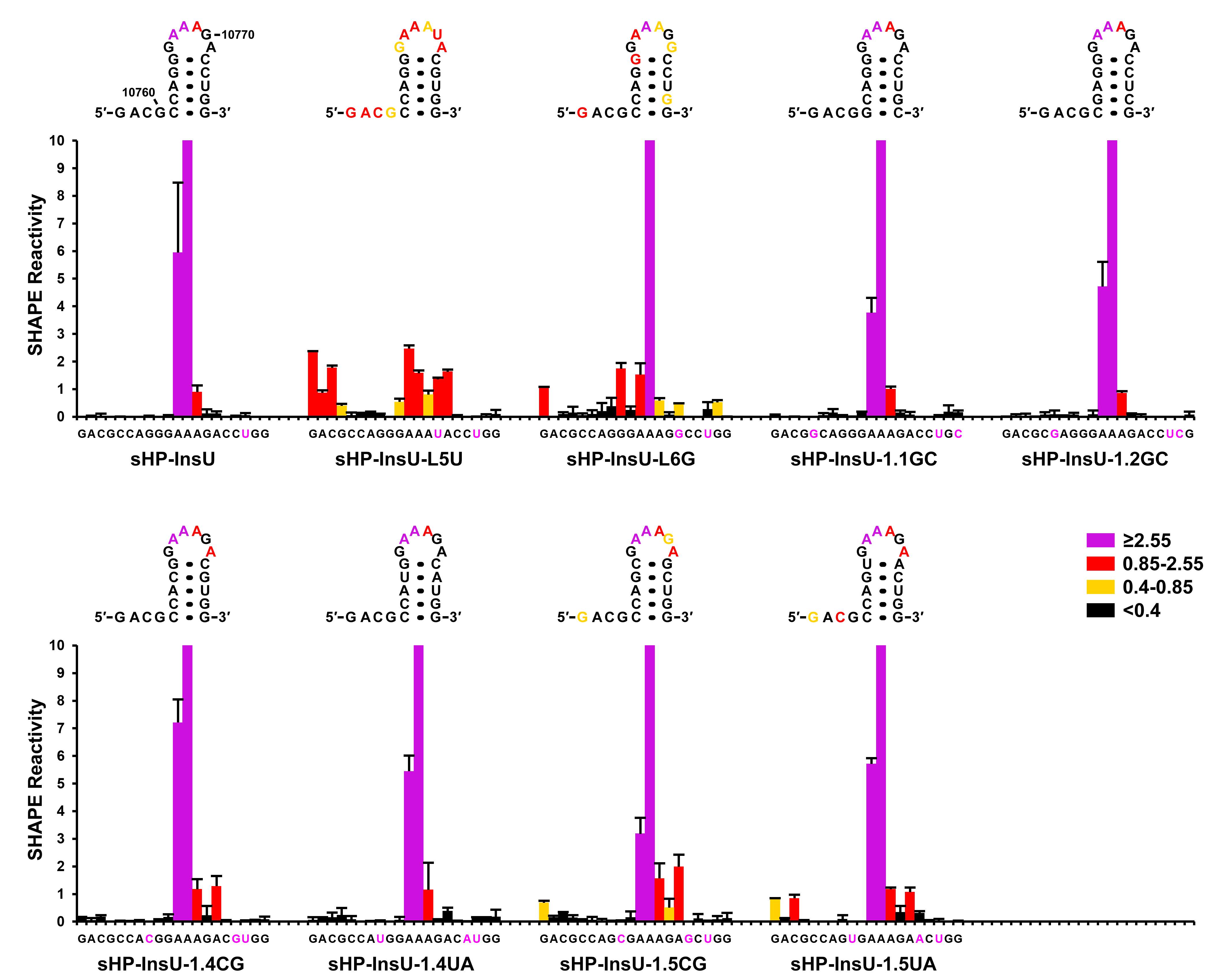

### Figure 6-figure supplement 3

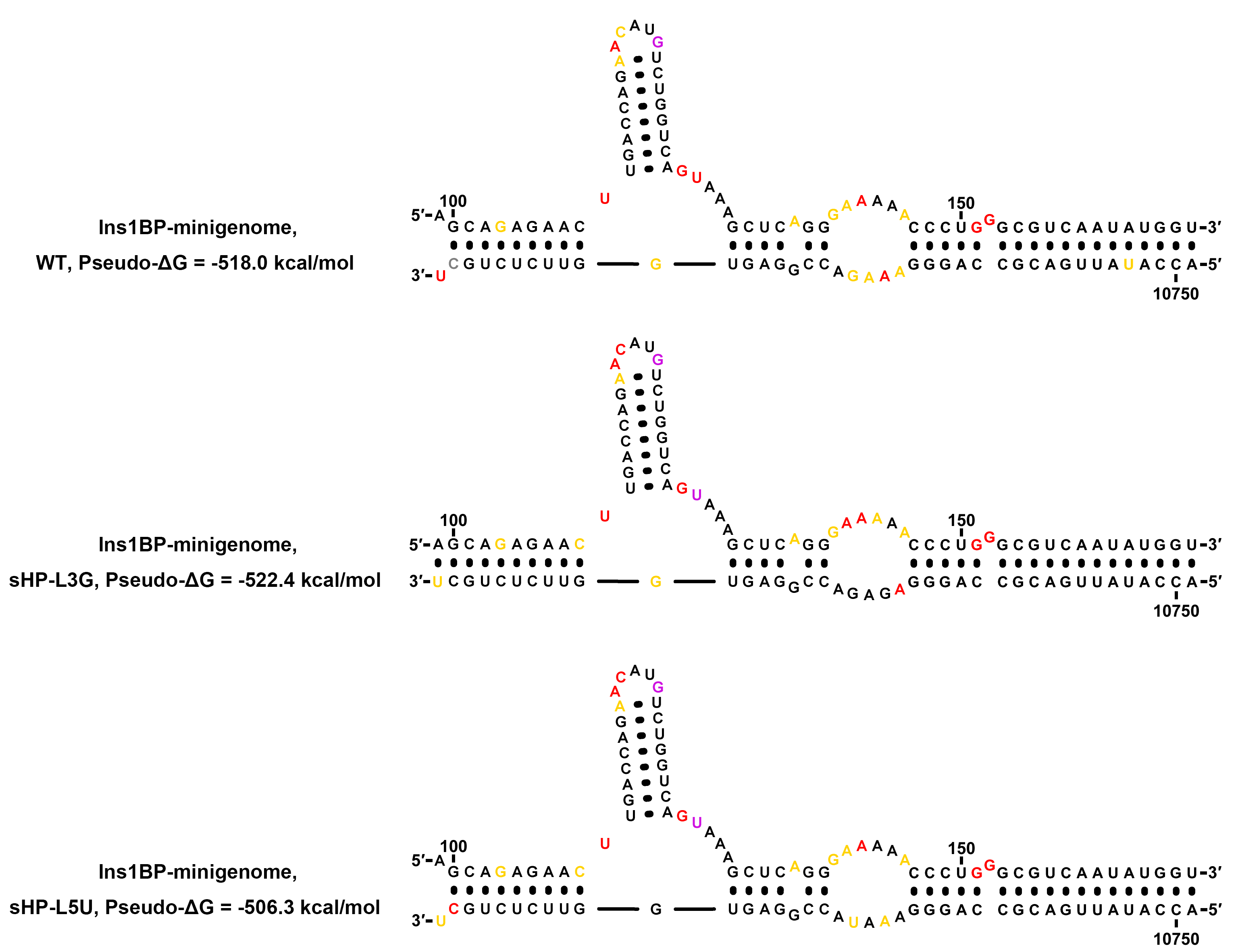

### Figure 7-figure supplement 1

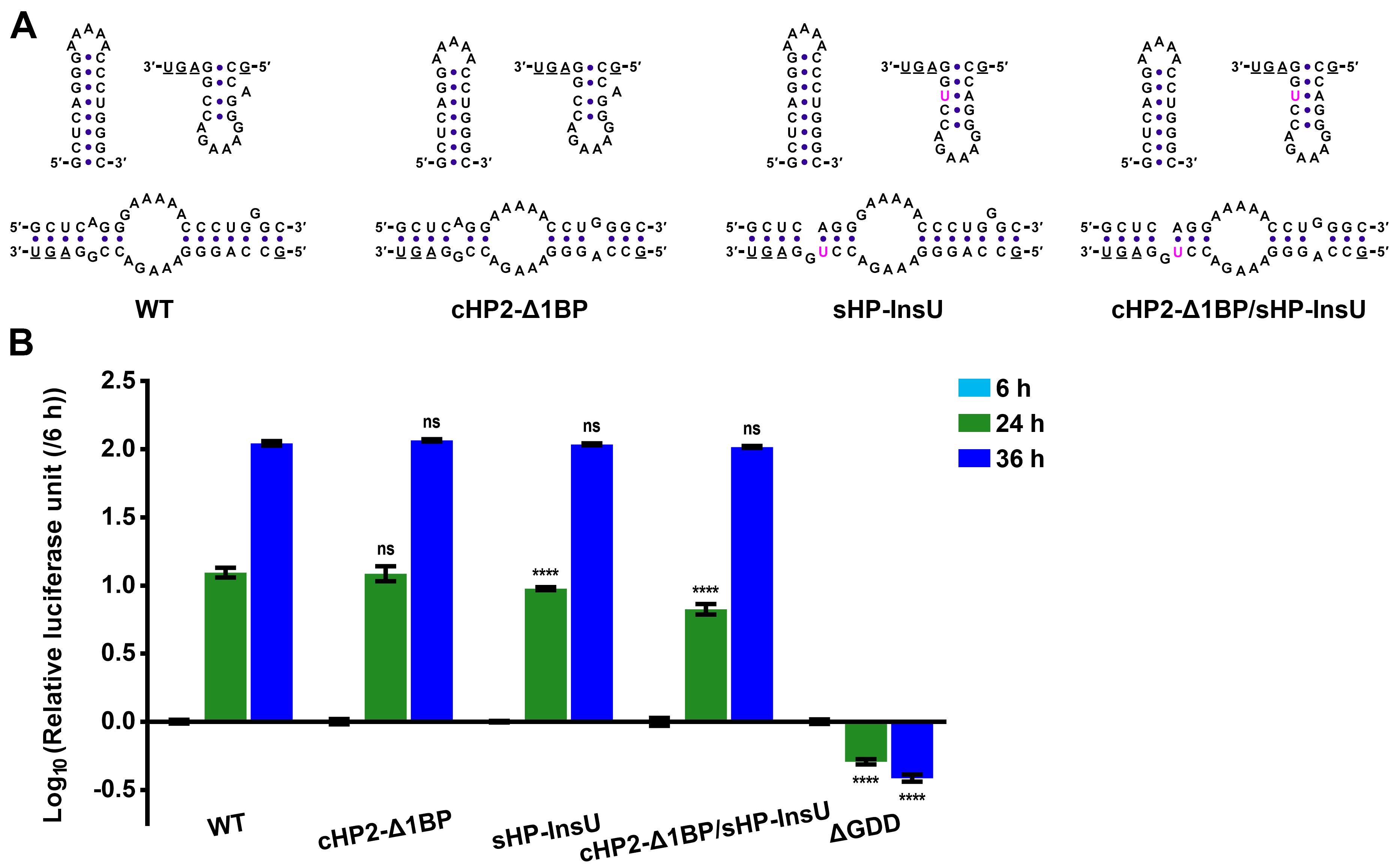

### Figure 8-figure supplement 1

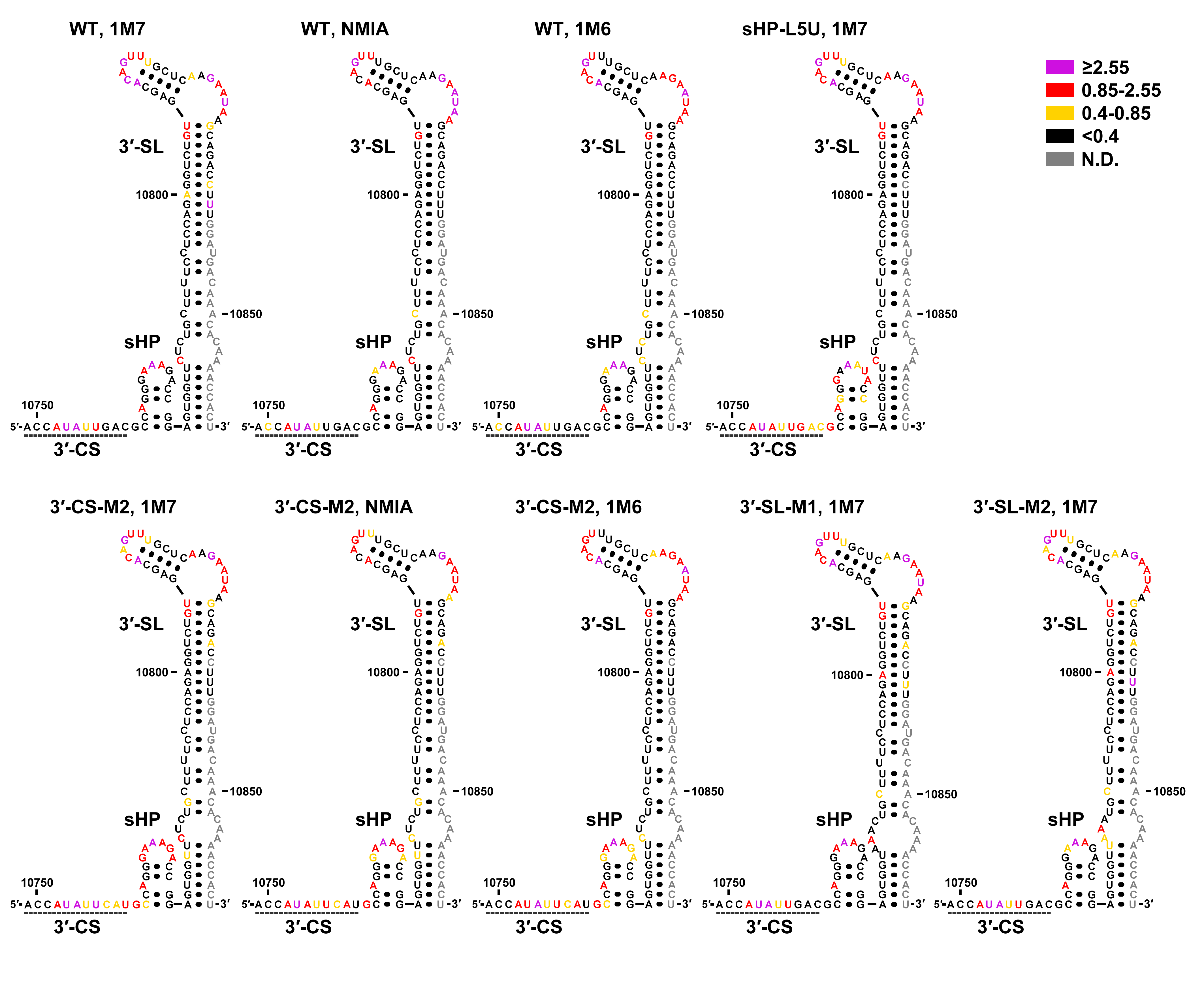

### Figure 8-figure supplement 2

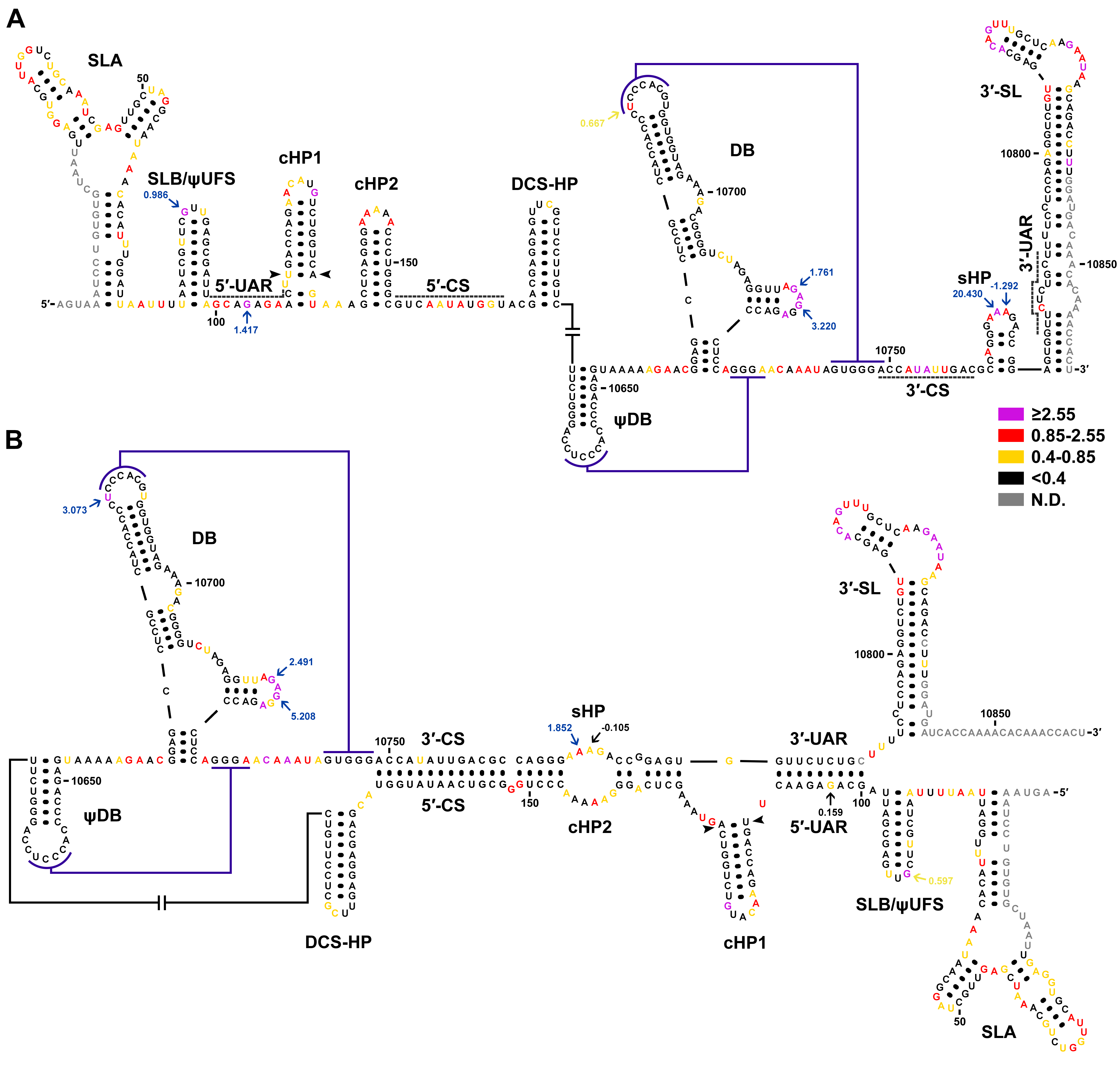

### Figure 8-figure supplement 3

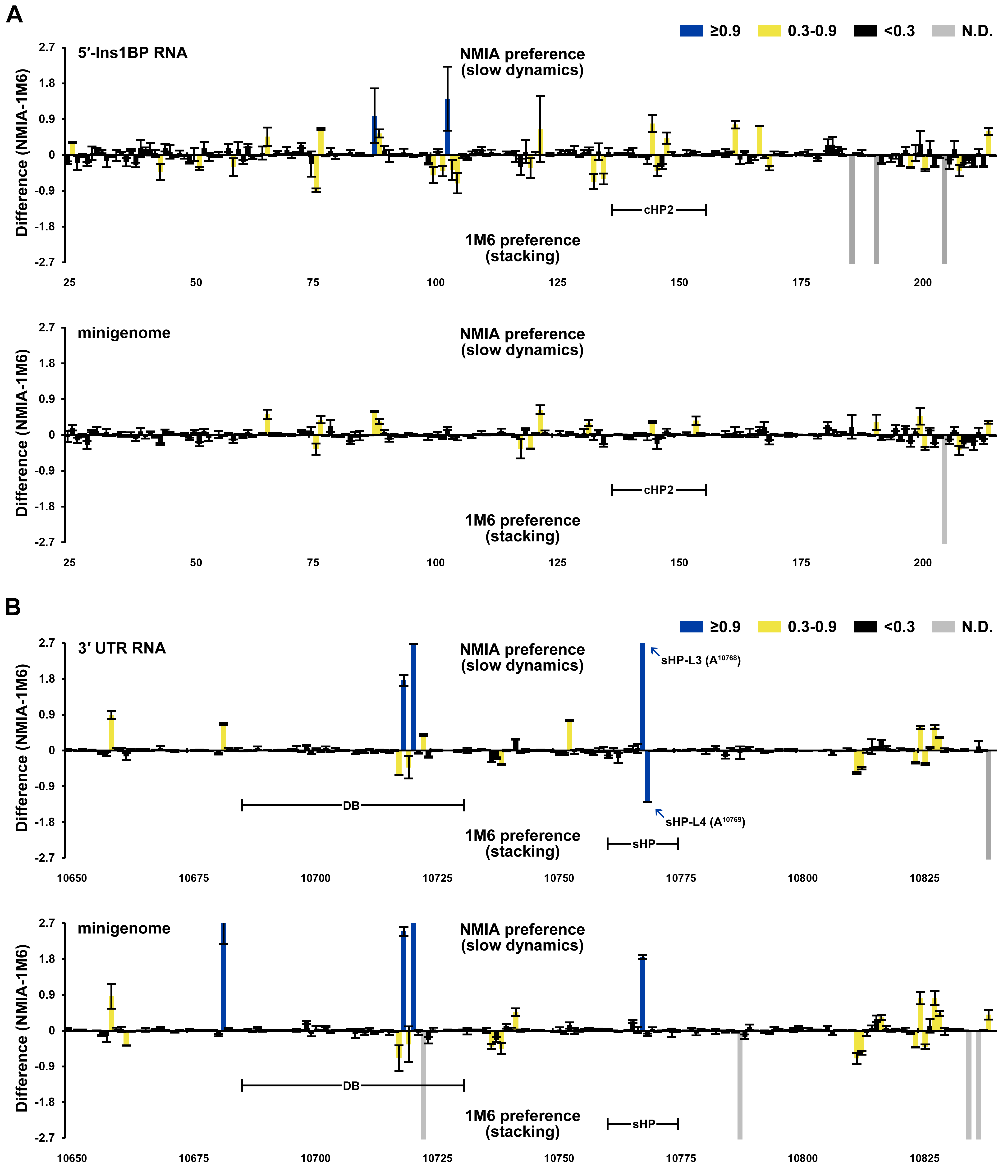

### Figure 9-figure supplement 1

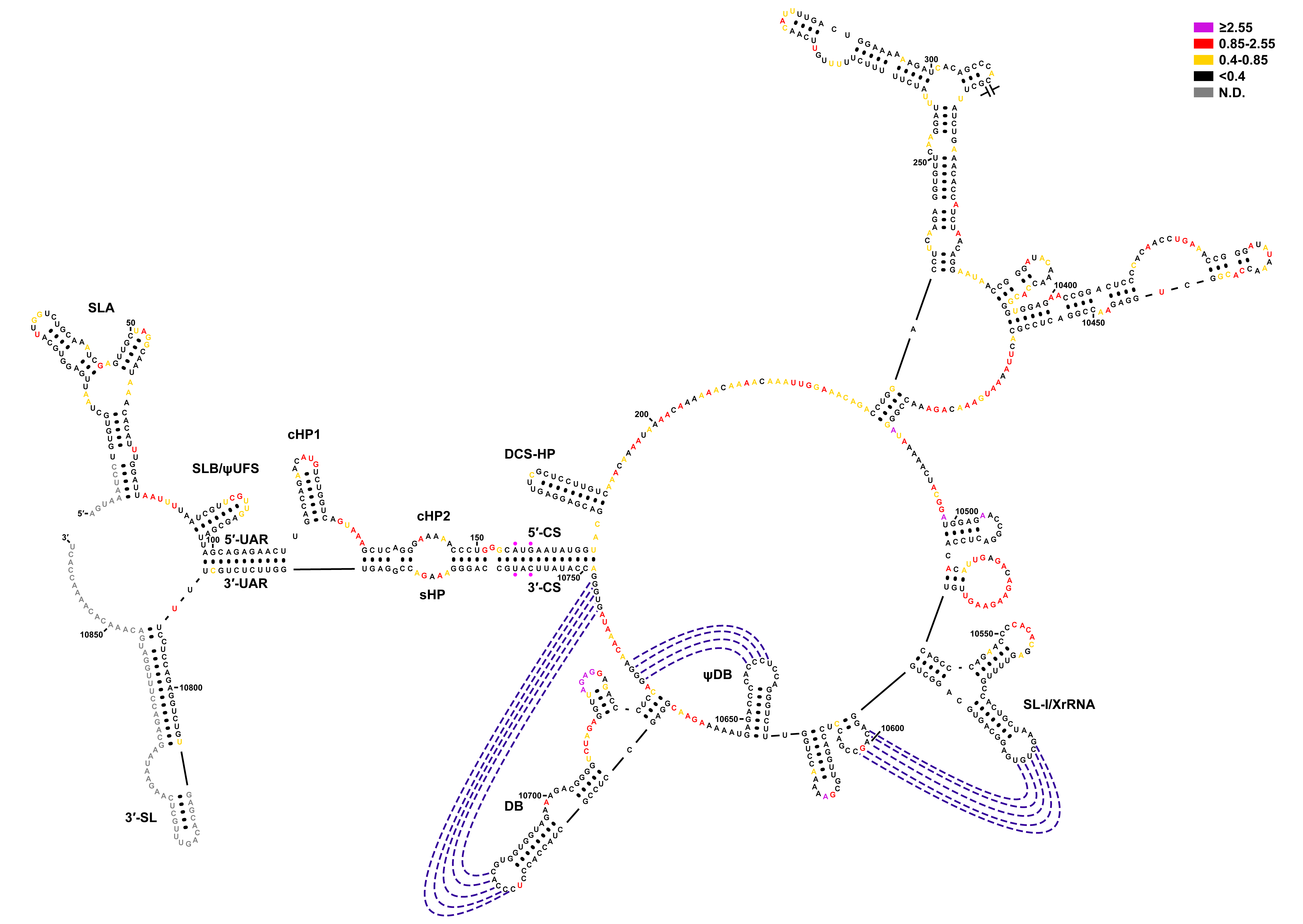

### Figure 11-figure supplement 1

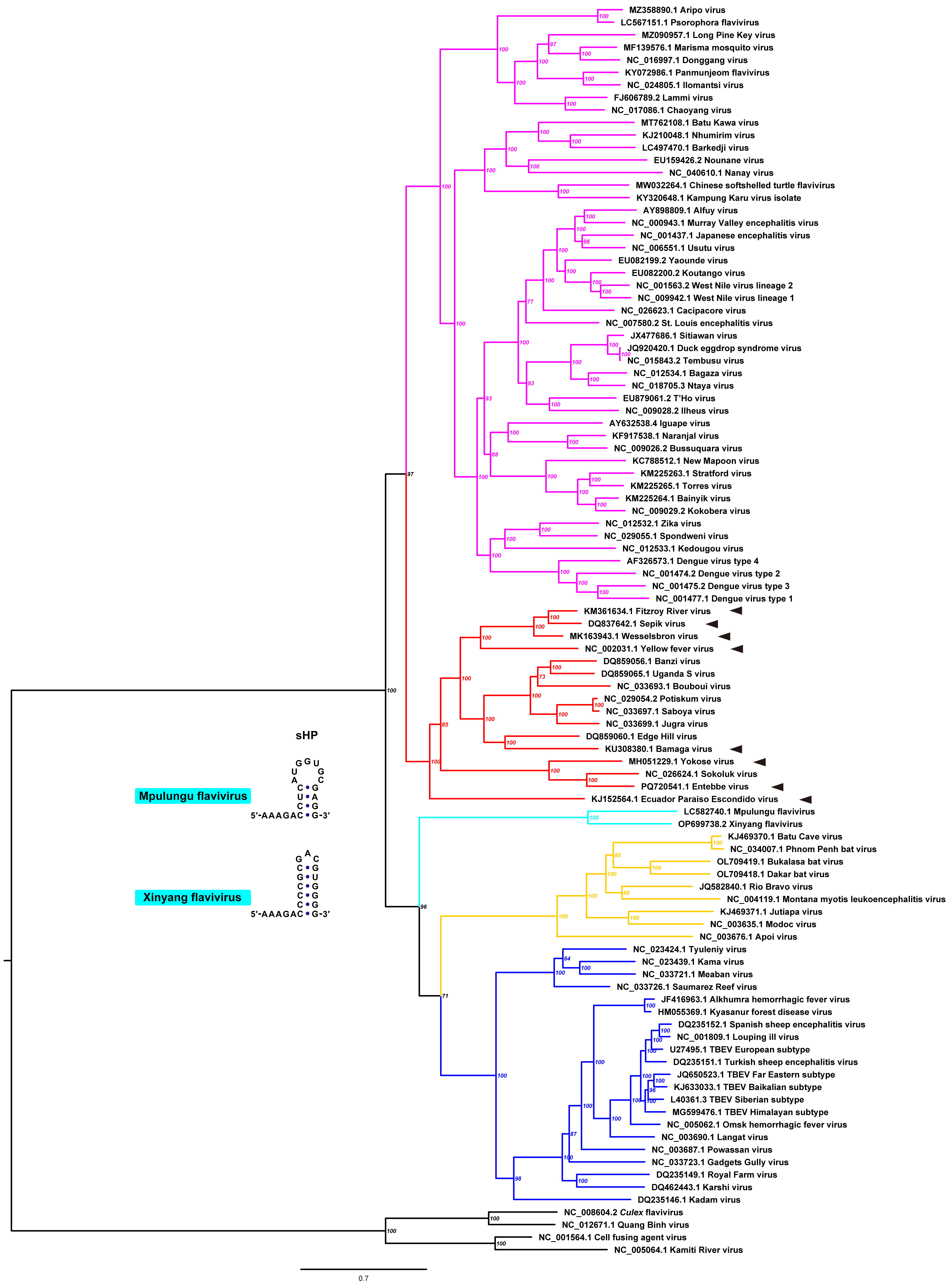

### Figure 11-figure supplement 2

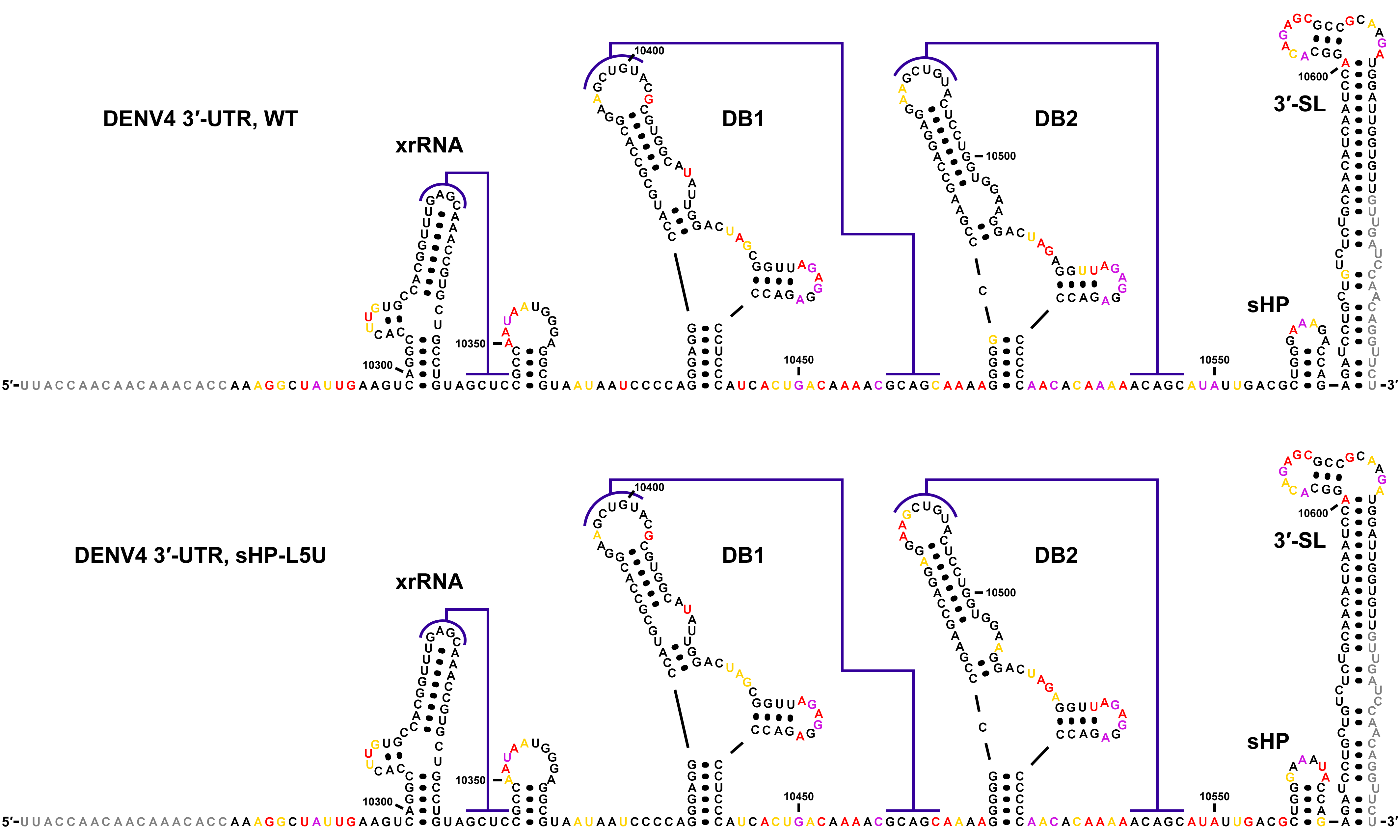

### Figure 11-figure supplement 3

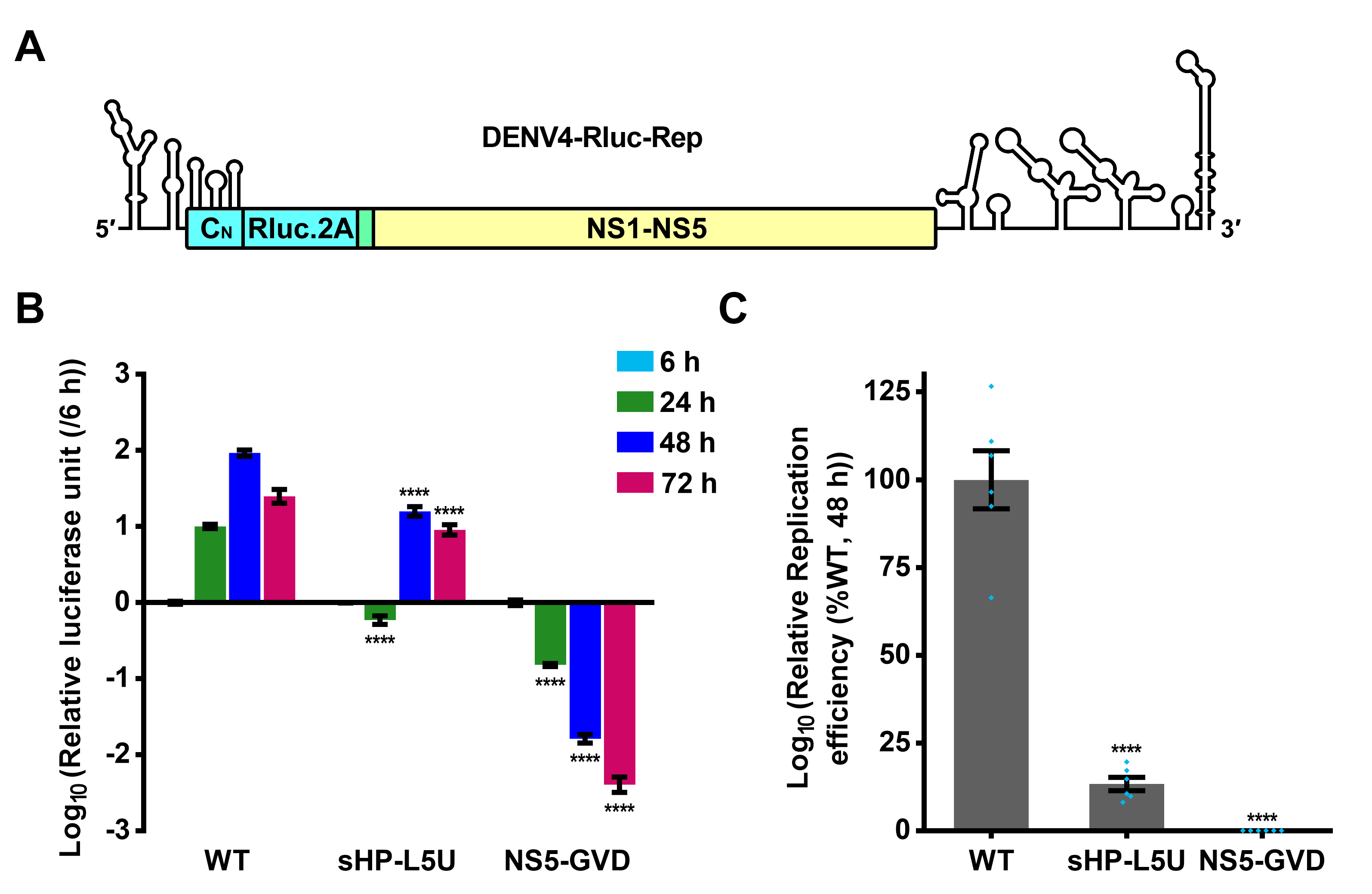

### Figure 11-figure supplement 4

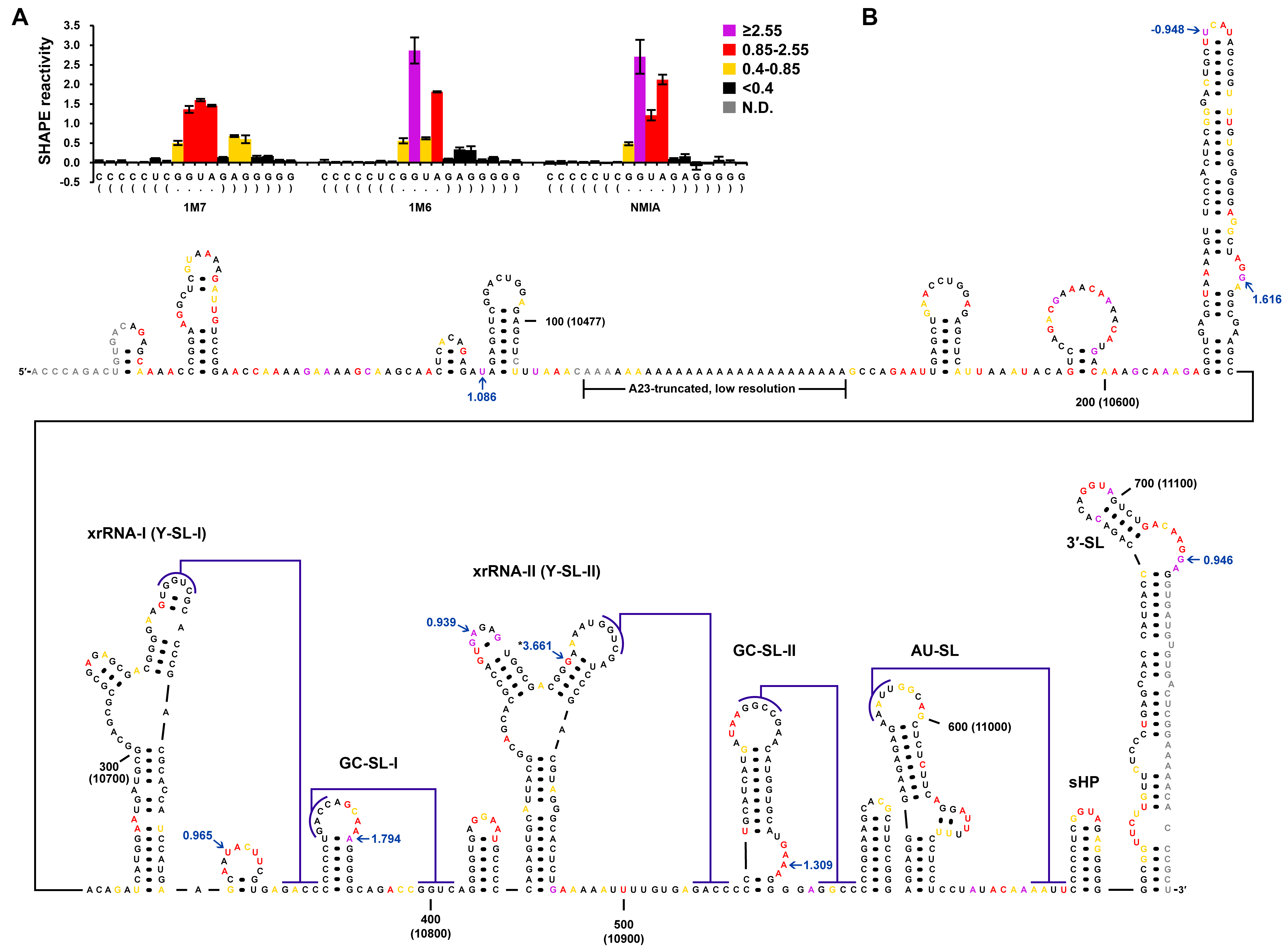

### Figure 11-figure supplement 5

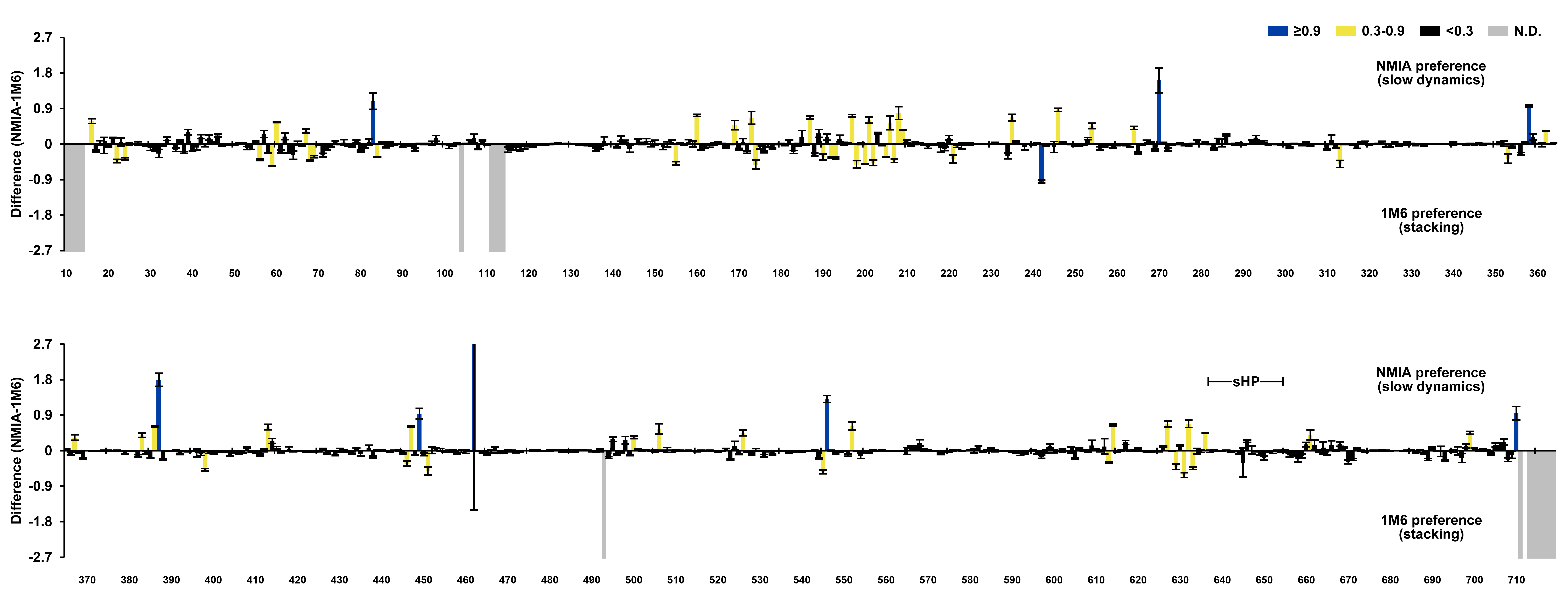

### Ins1BP-21-22-24-30-31-mini.pdf

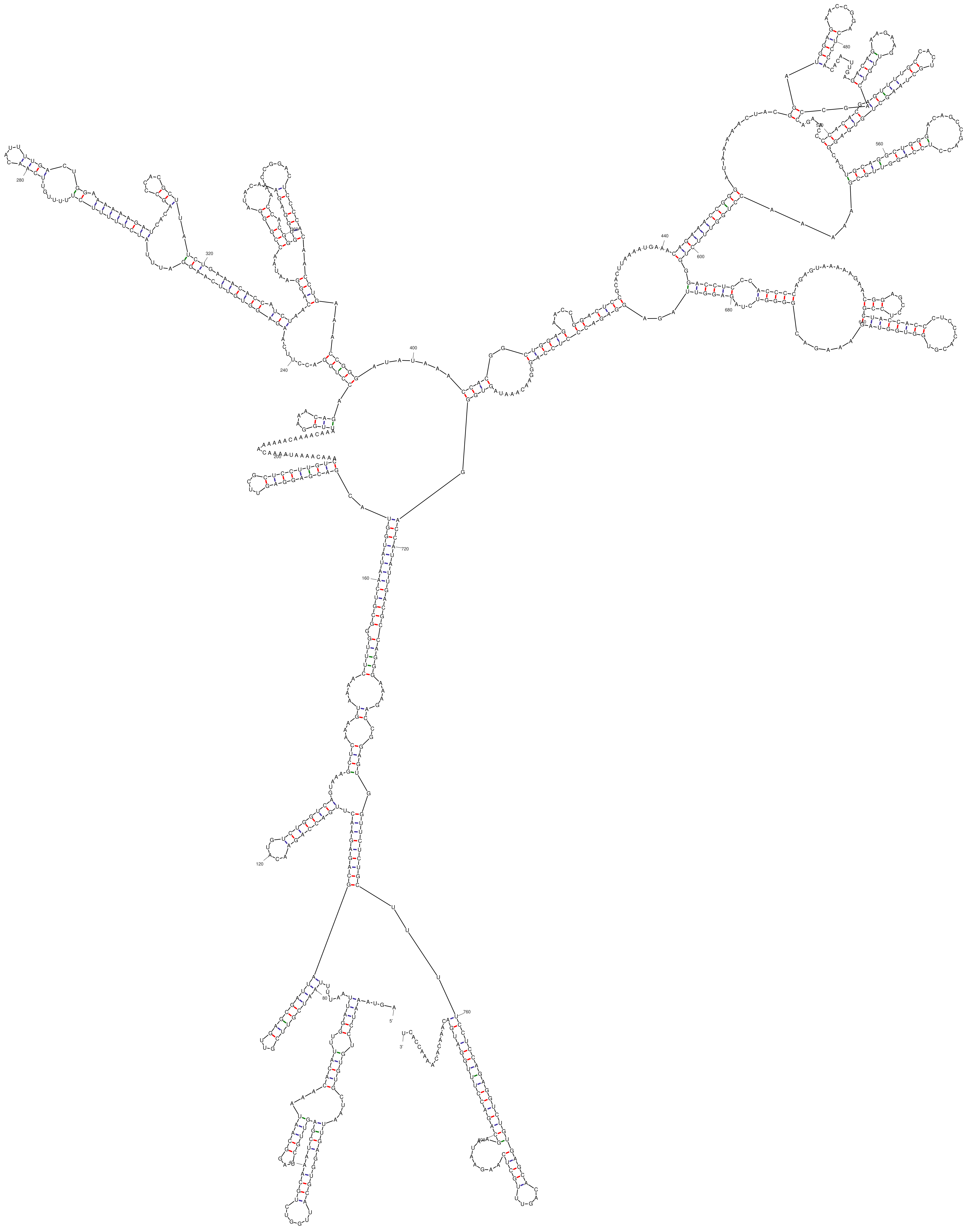

dG = -253.52 [Initially -281.30] 26May09-16-53-36

### Ins1BP-21-22-24-mini.pdf

Output of sir_graph (©)
mfold_util 4.7

Created Sat May 9 16:46:02 2026

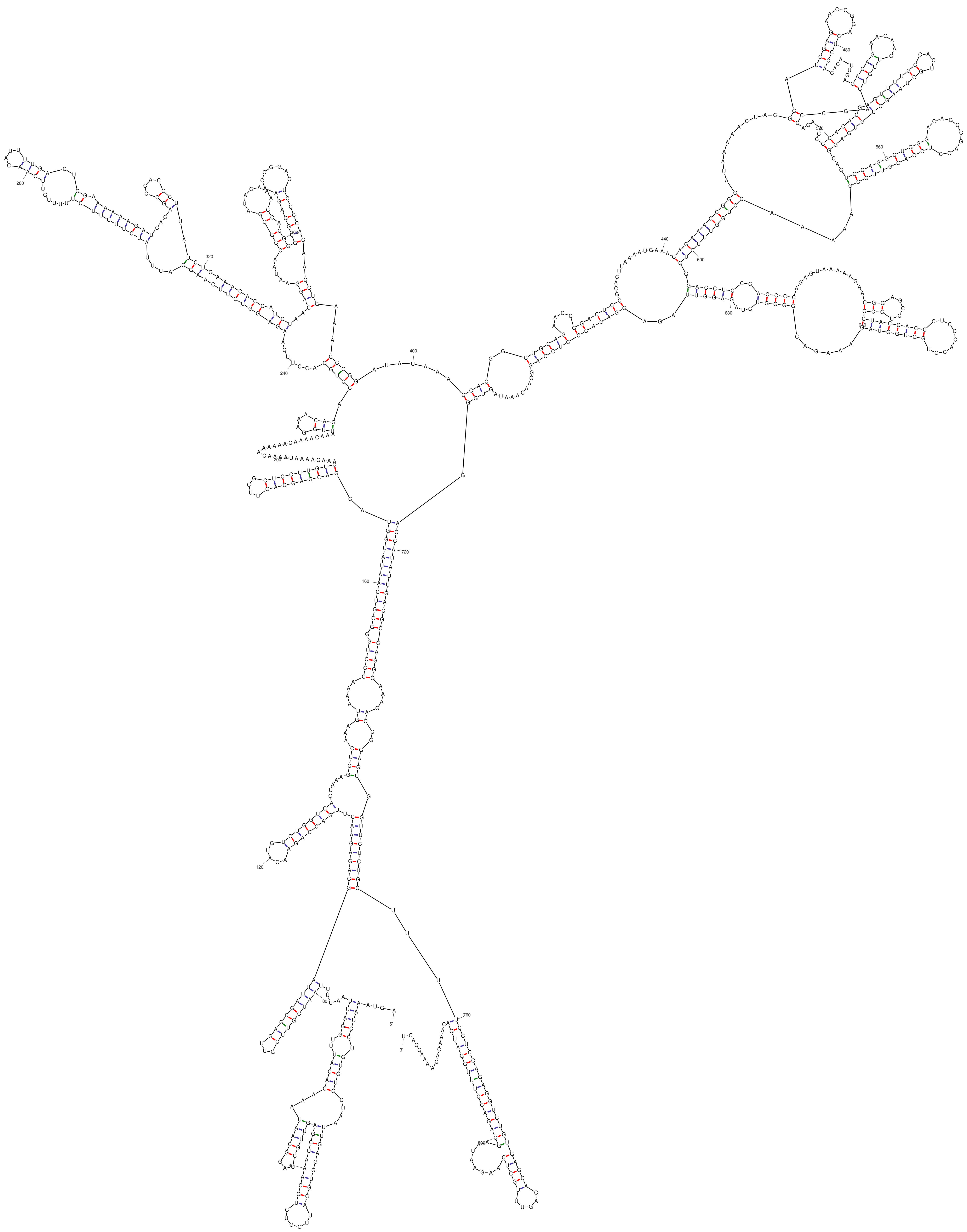


*dG = -259.02 [Initially -286.80] 26May09-16-45-58*

### Ins1BP-30-31-mini.pdf

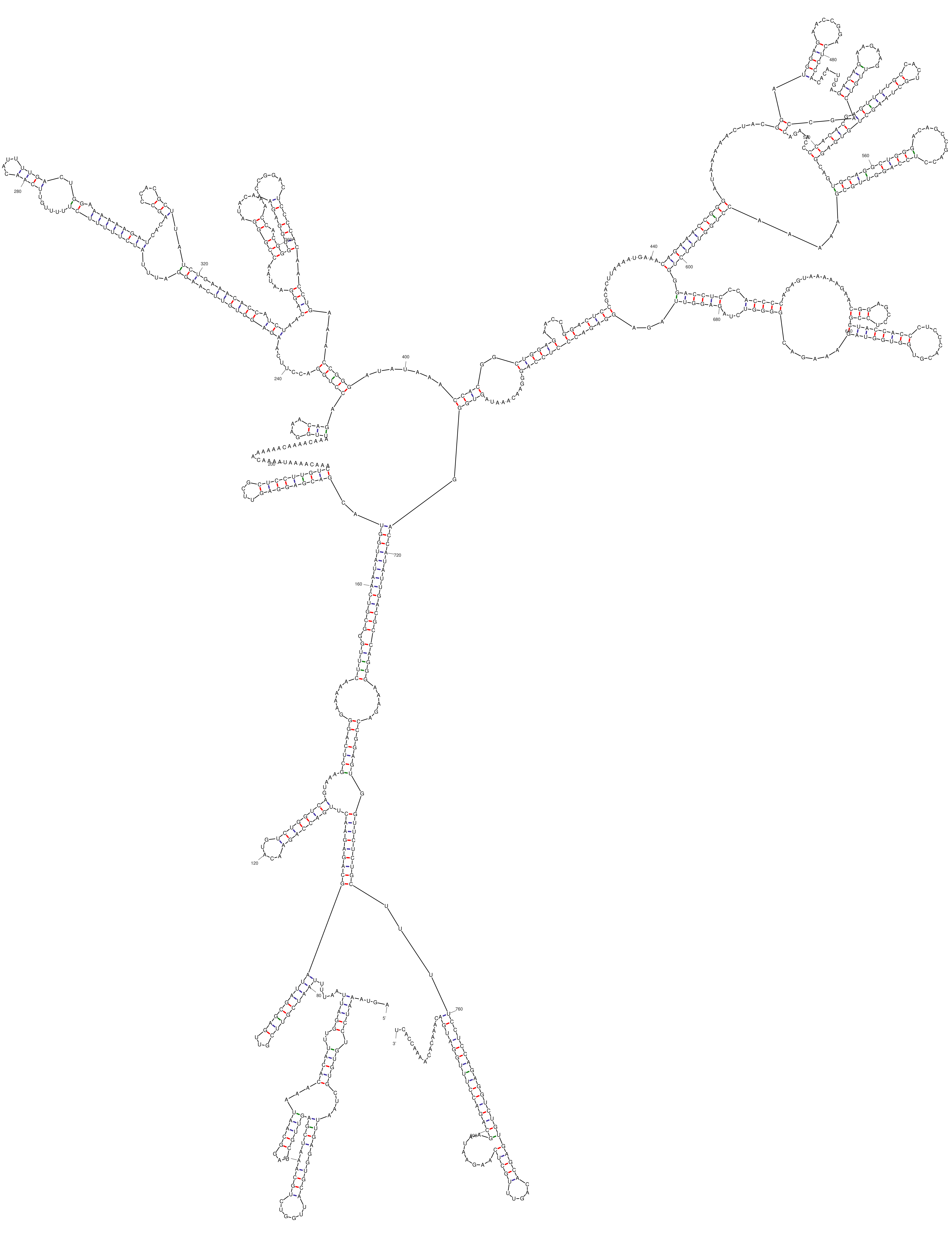


*dG = -255.32 [Initially -283.10] 26May09-16-49-47*

### Ins1BP-G21A-C31U-mini.pdf

Output of sir_graph (©)
mfold_util 4.7

Created Sat May  9 16:44:30 2026

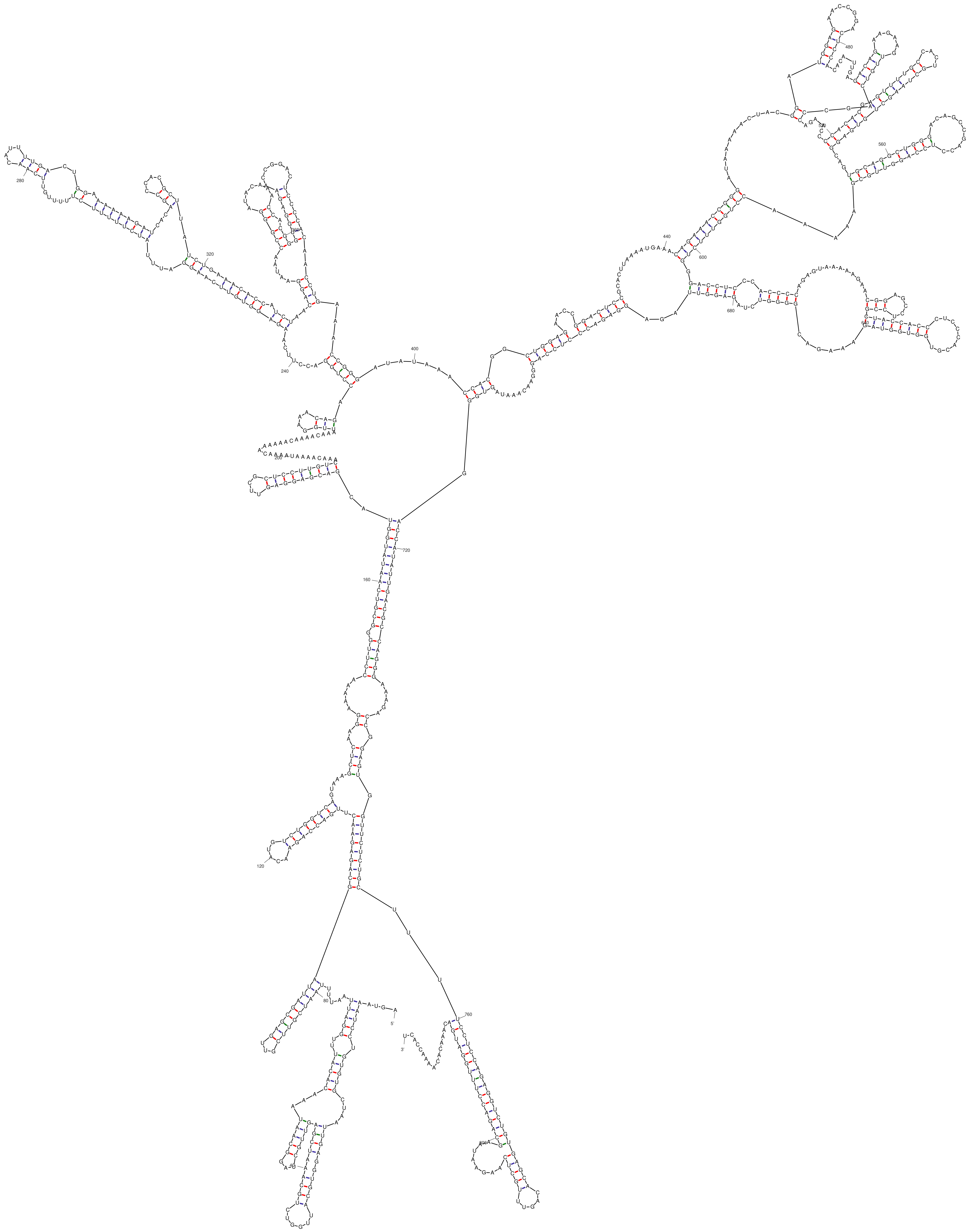

dG = -256.32 [Initially -284.10] 26May09-16-44-27

### Ins1BP-G21A-mini.pdf

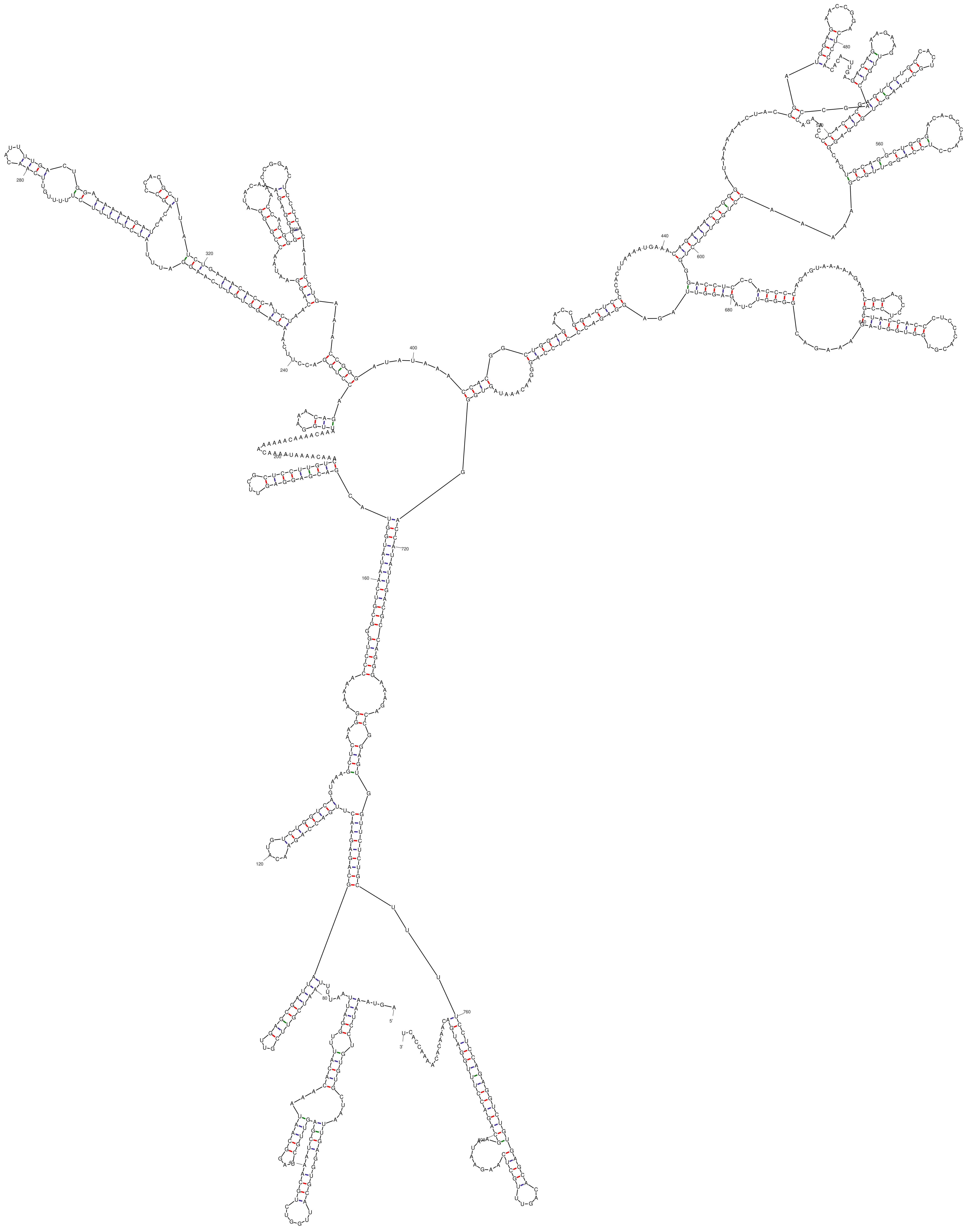

dG = -259.02 [Initially -286.80] 26May09-16-43-12

### Ins1BP-WT-mini.pdf

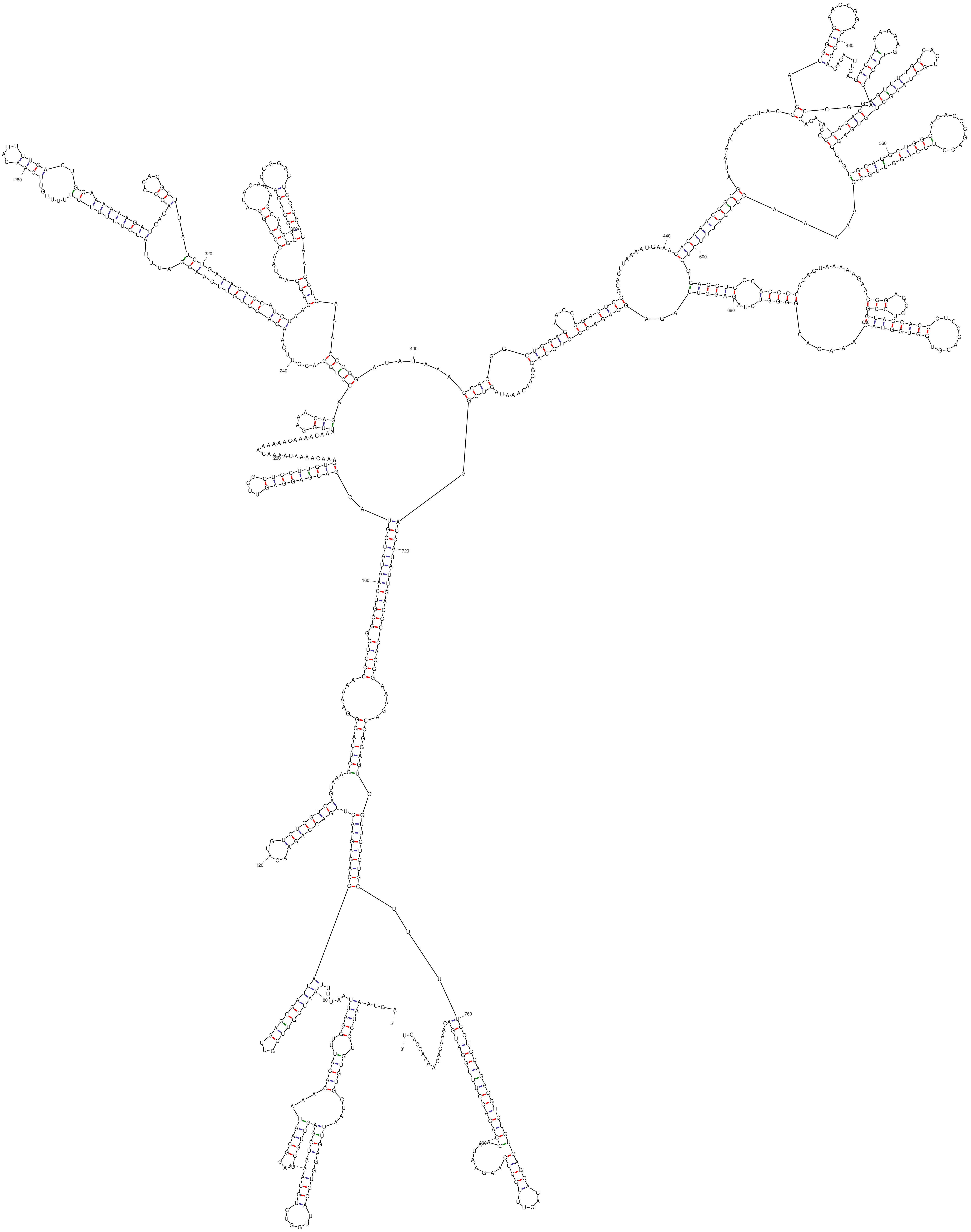


dG = -260.82 [Initially -288.60] 26May09-16-41-06
