## Supplementary material for "Novel conserved RNA structural motifs regulate genome cyclization of yellow fever virus": Figure 6-figure supplement 1

| | |
|---|---|
| | ..............((.((........)).)) |
| NC_002031.1 Yellow fever virus | --ACCAUAUUGACGCCAGGGAA--AGACC-GG |
| MK163943.1 Wesselsbron virus | --ACCAUAUUGACGCCAGGGAA--AGACC-GG |
| NC_008719.1 Sepik virus | --ACCAUAUUGACGCCUGGGAA--AGACC-GG |
| KM361634.1 Fitzroy River virus | --ACCAUAUUGACGCCUGGGAA--AGACC-GG |
| MH257544.1 Bamaga virus 3′-UTR | --AUCACAUUGACACCAGGGAU--AGACC-GG |
| PQ720541.1 Entebbe bat virus | --AAUAUGUUGACACCAGGGAA--AGACC-GG |
| MH051229.1 Yokose virus | --AAUAUGUUGACACCAGGGAA--AGACC-GG |
| NC_027999.1 Ecuador Paraiso Escondido virus | ---UCAUAUUGACACCAGGGAAA-AGACC-GG |
| NC_017086.1 Chaoyang virus | -CACUAUAUCGACACUUGGGAA--AGACC-AG |
| PP329310.1 Lammi virus | -CACUAUAUCGACACUUGGGAA--AGACC-AG |
| MF139576.1 Marisma mosquito virus | UUAUUAUAUCGACACCUAGGAA--AGACU-GG |
| MZ090957.1 Long Pine Key virus | --AUUAUAUCGACACCUAGGAA--AGACU-GG |
| MZ358890.1 Aripo virus | -AAUUAUAUCGACACCUAGGAA--AGACU-GG |
| NC_016997.1 Donggang virus | -UAUUAUAUCGACACCUAGGAA--AGACU-GG |
| LC567151.1 Psorophora flavivirus | -AAUUAUAUCGACACCUAGGAA--AGACU-GG |
| NC_040610.1 Nanay virus | ---GCAUAUUGACACCAGGGAU--AGACC-GG |
| LC497470.1 Barkedji virus | --AGCAUAUUGACACCUGGGAU--AGACC-GG |
| NC_024017.1 Nhumirim virus | -UAGCAUAUUGACACUUGGGAAG-AGACC-AG |
| MT762108.1 Batu Kawa virus | ----CAUAUUGACACCAGGGAA--AGACC-GG |
| NC_001477.1 Dengue virus type1 | -CAGCAUAUUGACGCU-GGGAG--AGACC-AG |
| NC_001474.2 Dengue virus type 2 | -CAGCAUAUUGACGCU-GGGAA--AGACC-AG |
| NC_001475.2 Dengue virus type 3 | -CAGCAUAUUGACGCU-GGGAG--AGACC-AG |
| AF326573.1 Dengue virus type 4 | -CAGCAUAUUGACGCU-GGGAA--AGACC-AG |
| NC_035889.1 Zika virus | --AGCAUAUUGACGCU-GGGAA--AGACC-AG |
| NC_007580.2 St. Louis encephalitis virus | --AGCAUAUUGACACC-UGGAA--AGACA-GG |
| NC_012533.1 Kedougou virus | -CAGCAUAUUGACACCUGGGAA--AGACC-GG |
| NC_009026.2 Bussuquara virus | --AGCAUAUUGACACCUGGGAU--AGACC-GG |
| AY632538.4 Iguape virus | --AACAUACUGACACCUGGGAA--AGACC-GG |
| NC_040776.1 Rocio virus | -CAGCAUAUUGACACCUGGGAA--AGACC-GG |
| NC_009028.2 Ilheus virus | --AGCAUAUUGACGCCUGGGAA--AGACC-GG |
| EU879061.2 T’Ho virus | -CAGCAUAUUGACACCUGGGAU--AGACUGGG |
| NC_015843.2 Tembusu virus | --AGCAUAUUGACACCUGGGAU--AGACUAGG |
| NC_012534.1 Bagaza virus | --AGCAUAUUGACACCUGGGAG--AGACUAGG |
| NC_018705.3 Ntaya virus | --AGCAUAUUGACACCUGGGAA--AGACUAGG |
| NC_009942.1 West Nile virus lineage 1 | --AGCAUAUUGACACCUGGGAU--AGACUAGG |
| NC_001563.2 West Nile virus lineage 2 | --AGCAUAUUGACACCUGGGAU--AGACUAGG |
| NC_006551.1 Usutu virus | -CAGCAUAUUGACACCUGGGAAUUAGACUAGG |
| NC_001437.1 Japanese encephalitis virus | -CAGCAUAUUGACACCUGGGAAU-AGACUGGG |
| NC_000943.1 Murray Valley encephalitis virus | --AGCAUAUUGACACCUGGGAAA-AGACUAGG |
| AY898809.1 Alfuy virus | --AGCAUACUGACACCUGGGAC--AGACUAGG |
| NC_009029.2 Kokobera virus | --AGCAUAUUGACACCUGGGAA--AGACUAGG |
| NC_032088.1 New Mapoon virus | --AGCAUAUUGACACCUGGGAU--AGACUAGG |
