## Supplementary material for "Novel conserved RNA structural motifs regulate genome cyclization of yellow fever virus": Figure 11-Supplementary File 1: LocARNA - Results-5-500-5445874.pdf

### Freiburg RNA Tools
### LocARNA - Results



**Input and runtime details for job 5445874 submitted on 06.07.2026 at 10:40 CEST**

[ Hide Input Parameters ]   [ Hide Job Execution Details ]

**Sequence and Structure**

| | |
|---|---|
| Sequence Input in FASTA Format | [.fa] |

**Alignment**

| | |
|---|---|
| Alignment Type | Global |
| Alignment Mode | no |

**Alignment Scoring**

| | |
|---|---|
| Structure Weight | 200 |
| Indel Opening Score | -800 |
| Indel Score | -50 |
| Use RIBOSUM | yes |
| Match Score | 50 |
| Mismatch Score | 0 |

**RNA Folding**

| | |
|---|---|
| Local folding: max. base pair span | 150 |
| Temperature in °C | 37 |
| Energy parameter set (Vienna package) | Turner model, 2004 |
| Upload dot plots | not provided |

**Heuristics for speed/accuracy tradeoff**

| | |
|---|---|
| Minimal Pair Probability | 0.0005 |
| Minimal Probability for Guide Tree Construction | 0.005 |
| Maximal Difference for Sizes of Matched Arcs | 30 |
| Maximal Difference for Alignment Edges | 60 |

**Other Parameters**

| | |
|---|---|
| Ignore Constraints | no |
| Disallow Lonely Pairs | yes |
| Use alifold consensus dot plots | no |
| Consistency Transformation (LocARNA-P only) | no |
| Keep Sequence Input Order | no |
| Stockholm alignment output | yes |

**Job ID 5445874 (server version 5.0.14)**

| | |
|---|---|
| Job Submitted & Queued | @ Mon Jul 06 10:40:32 CEST 2026 |
| Pre-Processing Started | @ Mon Jul 06 10:40:42 CEST 2026 |
| LocARNA Started | @ Mon Jul 06 10:40:42 CEST 2026 |
| LocARNA Finished & Post-Processing | @ Mon Jul 06 10:43:15 CEST 2026 |
| Post-Processing Finished | @ Mon Jul 06 10:44:17 CEST 2026 |
| Job Completed | @ Mon Jul 06 10:44:20 CEST 2026 |

DIRECT ACCESS: https://rna.informatik.uni-freiburg.de/RetrieveResults.jsp?jobID=5445874&toolName=LocARNA ( 30 days expiry )

**Output** **download complete results** [zip]

**Alignment annotated with RNAalifold consensus structure**

download alignment [ClustalW] [Stockholm] [consensus] [FASTA] [structure] [structure-pairs]   download alignment figure [.pdf] [.ps] [.png]

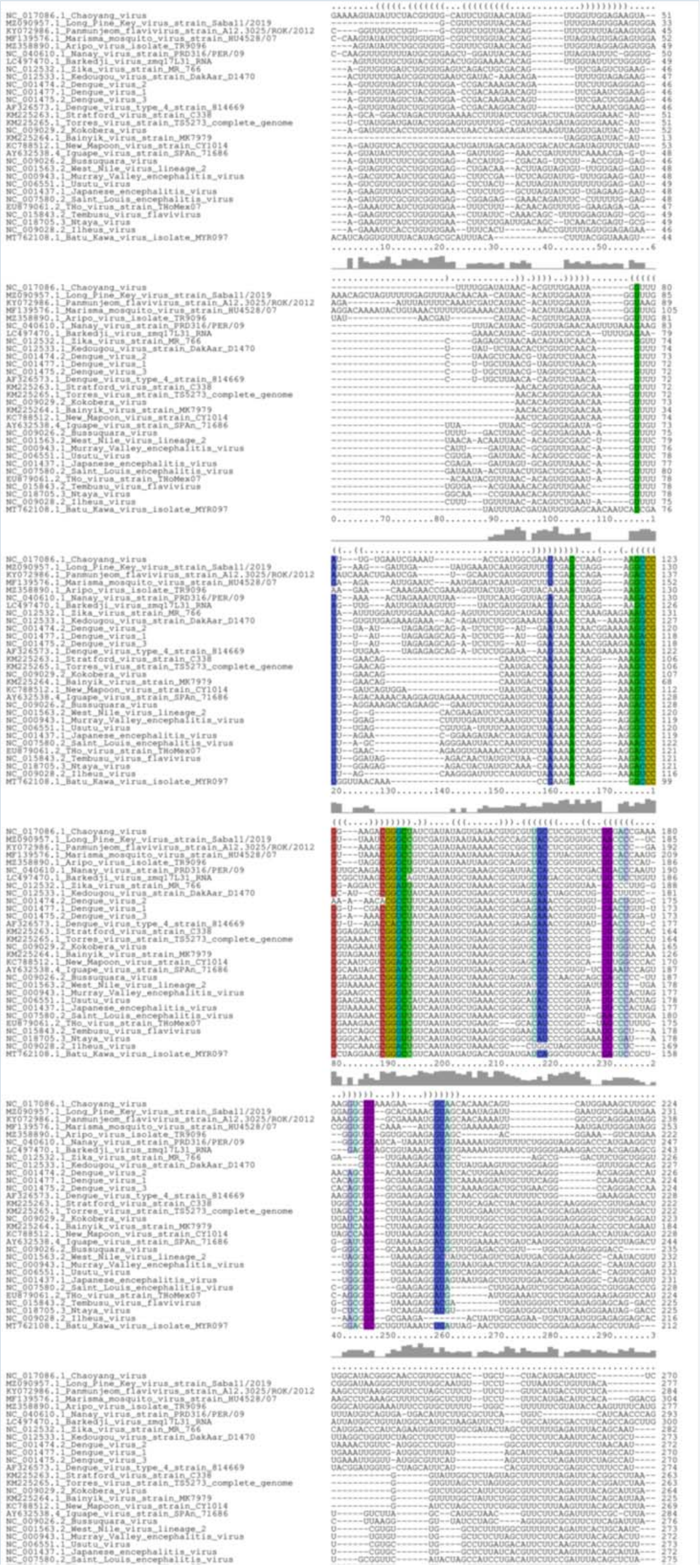

Bases are encoded by IUPAC codes. For interpretation of the colors see this legend. For a detailed description of the output, please see the help page.

#### ? Guide tree and Intermediate Alignments

download guide tree [NEWICK format] [svg] download [similarity matrix]

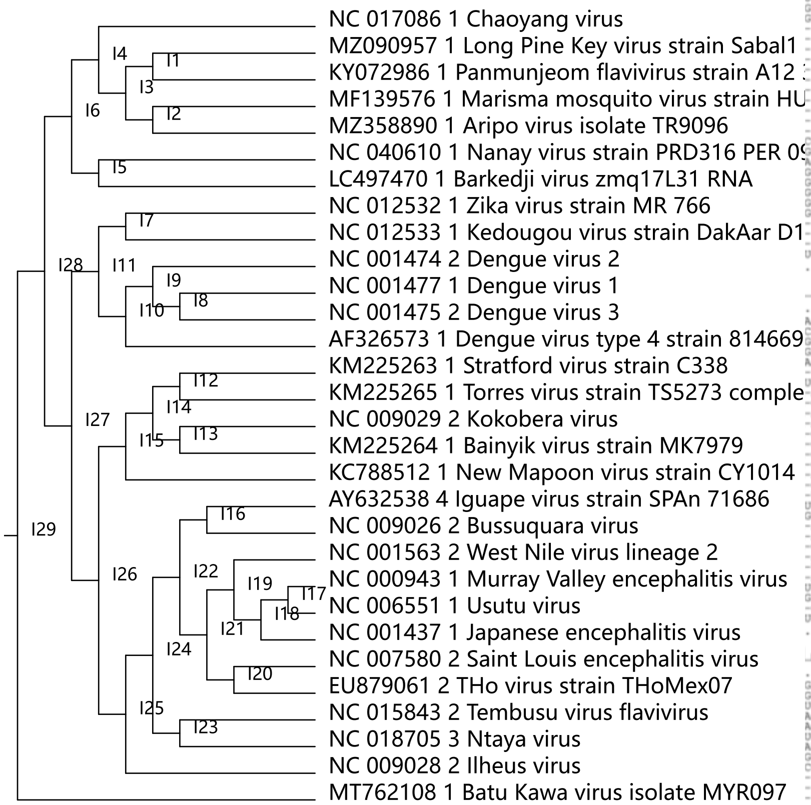


Intermediate Alignments

- intermediates/intermediate1.aln = MZ090957.1_Long_Pine_Key_virus_strain_Sabal1/2019 KY072986.1_Panmunjeom_flavivirus_strain_A12.3025/ROK/2012
- intermediates/intermediate2.aln = MF139576.1_Marisma_mosquito_virus_strain_HU4528/07 MZ358890.1_Aripo_virus_isolate_TR9096
- intermediates/intermediate3.aln = MZ090957.1_Long_Pine_Key_virus_strain_Sabal1/2019 KY072986.1_Panmunjeom_flavivirus_strain_A12.3025/ROK/2012 MF139576.1_Marisma_mosquito_virus_strain_HU4528/07 MZ358890.1_Aripo_virus_isolate_TR9096
- intermediates/intermediate4.aln = NC_017086.1_Chaoyang_virus MZ090957.1_Long_Pine_Key_virus_strain_Sabal1/2019 KY072986.1_Panmunjeom_flavivirus_strain_A12.3025/ROK/2012 MF139576.1_Marisma_mosquito_virus_strain_HU4528/07 MZ358890.1_Aripo_virus_isolate_TR9096
- intermediates/intermediate5.aln = NC_040610.1_Nanay_virus_strain_PRD316/PER/09 LC497470.1_Barkedji_virus_zmq17L31_RNA
- intermediates/intermediate6.aln = NC_017086.1_Chaoyang_virus MZ090957.1_Long_Pine_Key_virus_strain_Sabal1/2019 KY072986.1_Panmunjeom_flavivirus_strain_A12.3025/ROK/2012 MF139576.1_Marisma_mosquito_virus_strain_HU4528/07 MZ358890.1_Aripo_virus_isolate_TR9096 NC_040610.1_Nanay_virus_strain_PRD316/PER/09 LC497470.1_Barkedji_virus_zmq17L31_RNA
- intermediates/intermediate7.aln = NC_012532.1_Zika_virus_strain_MR_766 NC_012533.1_Kedougou_virus_strain_DakAar_D1470
- intermediates/intermediate8.aln = NC_001477.1_Dengue_virus_1 NC_001475.2_Dengue_virus_3
- intermediates/intermediate9.aln = NC_001474.2_Dengue_virus_2 NC_001477.1_Dengue_virus_1 NC_001475.2_Dengue_virus_3
- intermediates/intermediate10.aln = NC_001474.2_Dengue_virus_2 NC_001477.1_Dengue_virus_1 NC_001475.2_Dengue_virus_3 AF326573.1_Dengue_virus_type_4_strain_814669
- intermediates/intermediate11.aln = NC_012532.1_Zika_virus_strain_MR_766 NC_012533.1_Kedougou_virus_strain_DakAar_D1470 NC_001474.2_Dengue_virus_2 NC_001477.1_Dengue_virus_1 NC_001475.2_Dengue_virus_3 AF326573.1_Dengue_virus_type_4_strain_814669
- intermediates/intermediate12.aln = KM225263.1_Stratford_virus_strain_C338 KM225265.1_Torres_virus_strain_TS5273_complete_genome
- intermediates/intermediate13.aln = NC_009029.2_Kokobera_virus KM225264.1_Bainyik_virus_strain_MK7979
- intermediates/intermediate14.aln = KM225263.1_Stratford_virus_strain_C338 KM225265.1_Torres_virus_strain_TS5273_complete_genome NC_009029.2_Kokobera_virus KM225264.1_Bainyik_virus_strain_MK7979
- intermediates/intermediate15.aln = KM225263.1_Stratford_virus_strain_C338 KM225265.1_Torres_virus_strain_TS5273_complete_genome NC_009029.2_Kokobera_virus KM225264.1_Bainyik_virus_strain_MK7979 KC788512.1_New_Mapoon_virus_strain_CY1014
- intermediates/intermediate16.aln = AY632538.4_Iguape_virus_strain_SPAn_71686 NC_009026.2_Bussuquara_virus
- intermediates/intermediate17.aln = NC_000943.1_Murray_Valley_encephalitis_virus NC_006551.1_Usutu_virus
- intermediates/intermediate18.aln = NC_000943.1_Murray_Valley_encephalitis_virus NC_006551.1_Usutu_virus NC_001437.1_Japanese_encephalitis_virus
- intermediates/intermediate19.aln = NC_001563.2_West_Nile_virus_lineage_2 NC_000943.1_Murray_Valley_encephalitis_virus NC_006551.1_Usutu_virus NC_001437.1_Japanese_encephalitis_virus
- intermediates/intermediate20.aln = NC_007580.2_Saint_Louis_encephalitis_virus EU879061.2_THo_virus_strain_THoMex07
- intermediates/intermediate21.aln = NC_001563.2_West_Nile_virus_lineage_2 NC_000943.1_Murray_Valley_encephalitis_virus NC_006551.1_Usutu_virus NC_001437.1_Japanese_encephalitis_virus NC_007580.2_Saint_Louis_encephalitis_virus EU879061.2_THo_virus_strain_THoMex07
- intermediates/intermediate22.aln = AY632538.4_Iguape_virus_strain_SPAn_71686 NC_009026.2_Bussuquara_virus NC_001563.2_West_Nile_virus_lineage_2 NC_000943.1_Murray_Valley_encephalitis_virus NC_006551.1_Usutu_virus NC_001437.1_Japanese_encephalitis_virus NC_007580.2_Saint_Louis_encephalitis_virus EU879061.2_THo_virus_strain_THoMex07
- intermediates/intermediate23.aln = NC_015843.2_Tembusu_virus_flavivirus NC_018705.3_Ntaya_virus
- intermediates/intermediate24.aln = AY632538.4_Iguape_virus_strain_SPAn_71686 NC_009026.2_Bussuquara_virus NC_001563.2_West_Nile_virus_lineage_2 NC_000943.1_Murray_Valley_encephalitis_virus NC_006551.1_Usutu_virus NC_001437.1_Japanese_encephalitis_virus NC_007580.2_Saint_Louis_encephalitis_virus EU879061.2_THo_virus_strain_THoMex07 NC_015843.2_Tembusu_virus_flavivirus NC_018705.3_Ntaya_virus
- intermediates/intermediate25.aln = AY632538.4_Iguape_virus_strain_SPAn_71686 NC_009026.2_Bussuquara_virus NC_001563.2_West_Nile_virus_lineage_2 NC_000943.1_Murray_Valley_encephalitis_virus NC_006551.1_Usutu_virus NC_001437.1_Japanese_encephalitis_virus NC_007580.2_Saint_Louis_encephalitis_virus EU879061.2_THo_virus_strain_THoMex07 NC_015843.2_Tembusu_virus_flavivirus NC_018705.3_Ntaya_virus NC_009028.2_Ilheus_virus
- intermediates/intermediate26.aln = KM225263.1_Stratford_virus_strain_C338 KM225265.1_Torres_virus_strain_TS5273_complete_genome NC_009029.2_Kokobera_virus KM225264.1_Bainyik_virus_strain_MK7979 KC788512.1_New_Mapoon_virus_strain_CY1014 AY632538.4_Iguape_virus_strain_SPAn_71686 NC_009026.2_Bussuquara_virus NC_001563.2_West_Nile_virus_lineage_2

NC_000943.1_Murray_Valley_encephalitis_virus NC_006551.1_Usutu_virus NC_001437.1_Japanese_encephalitis_virus NC_007580.2_Saint_Louis_encephalitis_virus EU879061.2_THo_virus_strain_THoMex07 NC_015843.2_Tembusu_virus_flavivirus NC_018705.3_Ntaya_virus NC_009028.2_Ilheus_virus

- intermediates/intermediate27.aln = NC_012532.1_Zika_virus_strain_MR_766 NC_012533.1_Kedougou_virus_strain_DakAar_D1470 NC_001474.2_Dengue_virus_2 NC_001477.1_Dengue_virus_1 NC_001475.2_Dengue_virus_3 AF326573.1_Dengue_virus_type_4_strain_814669 KM225263.1_Stratford_virus_strain_C338 KM225265.1_Torres_virus_strain_TS5273_complete_genome NC_009029.2_Kokobera_virus KM225264.1_Bainyik_virus_strain_MK7979 KC788512.1_New_Mapoon_virus_strain_CY1014 AY632538.4_Iguape_virus_strain_SPAn_71686 NC_009026.2_Bussuquara_virus NC_001563.2_West_Nile_virus_lineage_2 NC_000943.1_Murray_Valley_encephalitis_virus NC_006551.1_Usutu_virus NC_001437.1_Japanese_encephalitis_virus NC_007580.2_Saint_Louis_encephalitis_virus EU879061.2_THo_virus_strain_THoMex07 NC_015843.2_Tembusu_virus_flavivirus NC_018705.3_Ntaya_virus NC_009028.2_Ilheus_virus
- intermediates/intermediate28.aln = NC_017086.1_Chaoyang_virus MZ090957.1_Long_Pine_Key_virus_strain_Sabal1/2019 KY072986.1_Panmunjeom_flavivirus_strain_A12.3025/ROK/2012 MF139576.1_Marisma_mosquito_virus_strain_HU4528/07 MZ358890.1_Aripo_virus_isolate_TR9096 NC_040610.1_Nanay_virus_strain_PRD316/PER/09 LC497470.1_Barkedji_virus_zmq17L31_RNA NC_012532.1_Zika_virus_strain_MR_766 NC_012533.1_Kedougou_virus_strain_DakAar_D1470 NC_001474.2_Dengue_virus_2 NC_001477.1_Dengue_virus_1 NC_001475.2_Dengue_virus_3 AF326573.1_Dengue_virus_type_4_strain_814669 KM225263.1_Stratford_virus_strain_C338 KM225265.1_Torres_virus_strain_TS5273_complete_genome NC_009029.2_Kokobera_virus KM225264.1_Bainyik_virus_strain_MK7979 KC788512.1_New_Mapoon_virus_strain_CY1014 AY632538.4_Iguape_virus_strain_SPAn_71686 NC_009026.2_Bussuquara_virus NC_001563.2_West_Nile_virus_lineage_2 NC_000943.1_Murray_Valley_encephalitis_virus NC_006551.1_Usutu_virus NC_001437.1_Japanese_encephalitis_virus NC_007580.2_Saint_Louis_encephalitis_virus EU879061.2_THo_virus_strain_THoMex07 NC_015843.2_Tembusu_virus_flavivirus NC_018705.3_Ntaya_virus NC_009028.2_Ilheus_virus
- intermediates/intermediate29.aln = NC_017086.1_Chaoyang_virus MZ090957.1_Long_Pine_Key_virus_strain_Sabal1/2019 KY072986.1_Panmunjeom_flavivirus_strain_A12.3025/ROK/2012 MF139576.1_Marisma_mosquito_virus_strain_HU4528/07 MZ358890.1_Aripo_virus_isolate_TR9096 NC_040610.1_Nanay_virus_strain_PRD316/PER/09 LC497470.1_Barkedji_virus_zmq17L31_RNA NC_012532.1_Zika_virus_strain_MR_766 NC_012533.1_Kedougou_virus_strain_DakAar_D1470 NC_001474.2_Dengue_virus_2 NC_001477.1_Dengue_virus_1 NC_001475.2_Dengue_virus_3 AF326573.1_Dengue_virus_type_4_strain_814669 KM225263.1_Stratford_virus_strain_C338 KM225265.1_Torres_virus_strain_TS5273_complete_genome NC_009029.2_Kokobera_virus KM225264.1_Bainyik_virus_strain_MK7979 KC788512.1_New_Mapoon_virus_strain_CY1014 AY632538.4_Iguape_virus_strain_SPAn_71686 NC_009026.2_Bussuquara_virus NC_001563.2_West_Nile_virus_lineage_2 NC_000943.1_Murray_Valley_encephalitis_virus NC_006551.1_Usutu_virus NC_001437.1_Japanese_encephalitis_virus NC_007580.2_Saint_Louis_encephalitis_virus EU879061.2_THo_virus_strain_THoMex07 NC_015843.2_Tembusu_virus_flavivirus NC_018705.3_Ntaya_virus NC_009028.2_Ilheus_virus MT762108.1_Batu_Kawa_virus_isolate_MYR097

##### RNAalifold consensus structure

download consensus figure [.pdf] [.ps] [.png]

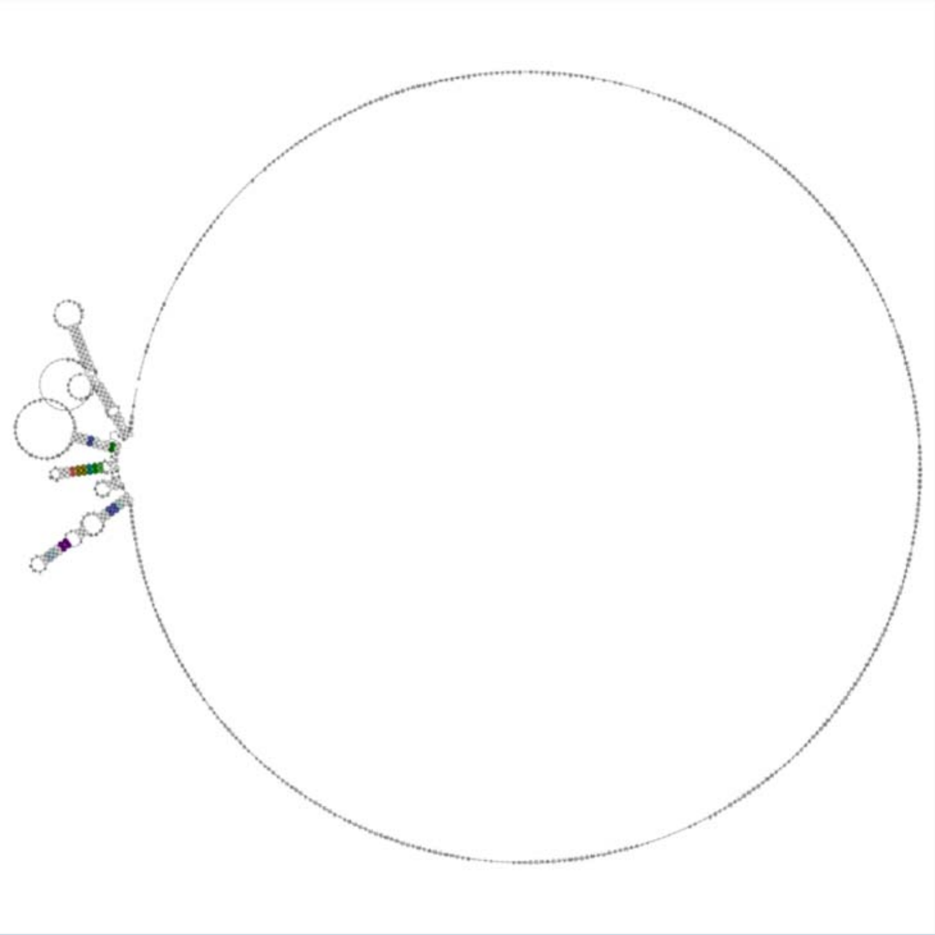

For interpretation of the colors see this legend. For a detailed description of the output, please see the help page.

##### Color Legend

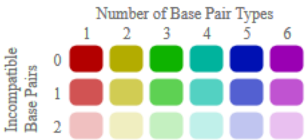


Compatible base pairs are colored, where the hue shows the number of different types C-G, G-C, A-U, U-A, G-U or U-G of compatible base pairs in the corresponding columns. In this way the hue shows sequence conservation of the base pair. The saturation decreases with the number of incompatible base pairs. Thus, it indicates the structural conservation of the base pair.

##### Job resubmission

Use the following button if you want to resubmit the job with altered input or parameterization: RESTART

**de.NBI usability assessment**

Please *support us* by filling this short quality survey for evaluating this de.NBI service.

**When using LocARNA please cite :**


- Sebastian Will, Tejal Joshi, Ivo L. Hofacker, Peter F. Stadler, and Rolf Backofen.
  LocARNA-P: Accurate boundary prediction and improved detection of structural RNAs
  RNA, 18 no. 5, pp. 900-14, 2012
- Sebastian Will, Kristin Reiche, Ivo L. Hofacker, Peter F. Stadler, and Rolf Backofen.
  Inferring non-coding RNA families and classes by means of genome-scale structure-based clustering
  PLoS Computational Biology, 3 no. 4, pp. e65, 2007
- Martin Raden, Syed M Ali, Omer S Alkhnbashi, Anke Busch, Fabrizio Costa, Jason A Davis, Florian Eggenhofer, Rick Gelhausen, Jens Georg, Steffen Heyne, Michael Hiller, Kousik Kundu, Robert Kleinkauf, Steffen C Lott, Mostafa M Mohamed, Alexander Mattheis, Milad Miladi, Andreas S Richter, Sebastian Will, Joachim Wolff, Patrick R Wright, and Rolf Backofen
  Freiburg RNA tools: a central online resource for RNA-focused research and teaching
  Nucleic Acids Research, 46(W1), W25-W29, 2018.


Results are computed with LocARNA version 1.9.1 linking Vienna RNA package 2.3.2
